# Barley malting for beer production – Impact of temperature on the barley-associated microbiome, malt quality and energy expenditure

**DOI:** 10.64898/2026.11.17.767257

**Authors:** Steven Kasozi Galiwango, Eoin Ó Cinnéide, Elif Göksu Kızılyar, Lars Pierco, Wouter De Sutter, Agata Soszka, Jolien Philips, Gust Heuvinck, Gert De Rouck, Ilse Van de Voorde, Florian Weiland

## Abstract

Malt production is an energy-intensive process, and elevated germination temperatures have been proposed to shorten it and thereby lower the energy demand. However, the effect of elevated temperatures on the barley-associated microbiome remains unexplored. Using a multi-metaomics approach, we explored microbial composition and function in four barley varieties malted at pilot scale under standard industrial conditions (IMR) and an accelerated regime using higher temperatures (AMR). AMR yielded malts with excellent friability, resulting in adequate wort viscosity and saccharification times despite lower amylase and β-glucanase activities, with no effect on proteolysis modulation. AMR substantially shifted microbial community composition. Fungi, notably *Alternaria alternata*, contributed xylan- and β-glucan-degrading enzymes, whereas abundant detection of the strong biofilm former *Pseudomonas* coincided with impaired wort filtration. Electrical energy use was lowered in AMR by approximately 5–9%, mainly through lower humidification and cooling demand. Malting with increased temperatures thus promises high-quality malt with time and energy savings, provided that the malting microbiome is monitored and controlled.

**Highlights:**

- Malting at 20°C reduces barley germination time by at least 2 days
- Increased temperatures shifted the malting microbiome towards more commonality
- Fungal glucanases and xylanases contribute to malt modification
- Biofilm-forming bacteria can likely impair lautering performance

## 1. Introduction

Barley (*Hordeum vulgare*) malt is a key raw material for beer brewing, with a production of more than 24 million metric tons in 2024 (Market Reports Word, 2025) and a market size of around 10 billion USD (*Barley Malt-Global Strategic Business Report*, 2026; Market Reports Word, 2025). Malt production is accomplished by controlled germination, which generates the enzymes needed for 1) cell wall breakdown (cytolysis), 2) starch degradation (amylolysis) and 3) the release of low-molecular-weight peptides and amino acids (FAN) for brewing yeast nutrition (proteolysis) (reviewed in (Narziß et al., 2024)). After germination, the kernels undergo drying and kilning to develop aroma and colour compounds while lowering the moisture content for prolonged storage. Modern high-throughput brewing techniques rely on the use of malt which exhibits a high degree of cytolysis and a limited degree of proteolysis: Insufficient cytolysis leads to the release of high-molecular-weight β-glucan from the barley cell walls during mashing, which increases wort viscosity and filtration times (reviewed in (Jin et al., 2004)). Conversely, excessive proteolysis impairs beer foam stability, and the resulting surplus of amino acids increases the concentration of staling off-flavour aldehydes formed via the Ehrlich pathway in yeasts and/or Strecker degradation (Bauwens et al., 2022; Evans and Hejgaard, 1999; Jaskula-Goiris et al., 2011). However, insufficient proteolysis causes fermentation problems as brewing yeasts rely on low-molecular-weight peptides and amino acids for their nitrogen supply (Fernández et al., 2000).

The malting process itself is energy-intensive, requiring about 110–170 kWh of electrical energy per ton of malt, of which approximately 25% is attributed to cooling during germination (Moresi and Cimini, 2025; Pensel, 1986). Germination at higher temperatures was therefore proposed to accelerate the production process and additionally save electricity costs for refrigeration (Müller and Methner, 2015). A further reported benefit of higher germination temperatures is the suppression of proteolysis while cytolytic activity is retained (Müller et al., 2014; Müller and Methner, 2015).

Although the microbial communities associated with barley germination have been described previously (Beccari et al., 2018; Bianco et al., 2018; Justé et al., 2011; Oliveira et al., 2012; Raulio et al., 2009; Tsitko et al., 2018) (also reviewed in (Galiwango et al., 2024)), the effect of an elevated-temperature malting regime on microbial composition, its development over the course of germination, the expressed metaproteome and the resulting malt quality has not yet been described. To address these knowledge gaps, we combined temporally resolved metagenomics, metaproteomics and monitored malt enzyme activities during germination under 1) a standard industrial malting regime and 2) an accelerated malting regime applying higher temperatures.

## 2. Material and Methods

### 2.1. Pilot-scale Malting

Four barley varieties were used: one six-row winter variety (Faro, FAR) and three two-row summer varieties (Fandaga (FDG), Fantex (FTX), and Planet (PLT)). All varieties were from the 2021 crop year (origin: France (FDG, FTX, FAR) and England (PLT)) and were malted in a pilot malting plant at KU Leuven-Ghent, Campus Rabot. The facility consisted of a cylindro-conical steeping tank and a doughnut-shaped germination/kilning vessel suitable for 500 kg of malt (BW Procestechniek NV, Belgium). The unit contains a false bottom which can hold 12 meshed stainless-steel buckets (20 cm diameter × 30 cm height), allowing different varieties to be malted under identical conditions. In this study, 2 kg of each variety (originating from the same barley variety lot) was malted in three replicates under two malting regimes: a standard industrial malting regime (IMR) and an accelerated malting regime (AMR) using elevated temperatures. The two regimes differed in their steeping and germination programmes, whereas the kilning programme was identical. For IMR, steeping consisted of a 4 h wet steep at 12 °C followed by a 9 h dry rest at 19–25 °C, and a second 4 h wet steep at 16 °C, which was followed by another 9 h dry rest at 19–25 °C; germination was then conducted at 16 °C for 120 h. For AMR, the first wet steep was conducted for 6 h at 20 °C, followed by an 18 h dry rest at 21°C and a second wet steep of 1 h at 20 °C; germination was then conducted at 20 °C for 74 h. For both regimes, germination was halted by withering the green malt at 35 °C for 4 h. The green malt was then dried at 45 °C for 2 h, followed by a linear temperature increase to 55 °C over 10 h, and held at 55 °C for 10 h. The temperature was then gradually increased in 5 °C steps of 1 h each from 60 to 75 °C. Finally, the green malt was kilned at 80 °C for 1 h and 85 °C for 4 h (total process time 35 h) to a final moisture content of about 3–4% (HE73 moisture analyser, Mettler-Toledo, Switzerland). The complete malting programme is shown in **Figure 1A and 1B** for IMR and AMR, respectively. After kilning, the malt was deculmed manually by rubbing the kilned grains against a sieve to break off the rootlets, packaged in 3 kg zip bags and stored at room temperature for three to four weeks before malt analysis.

**Figure 1.**
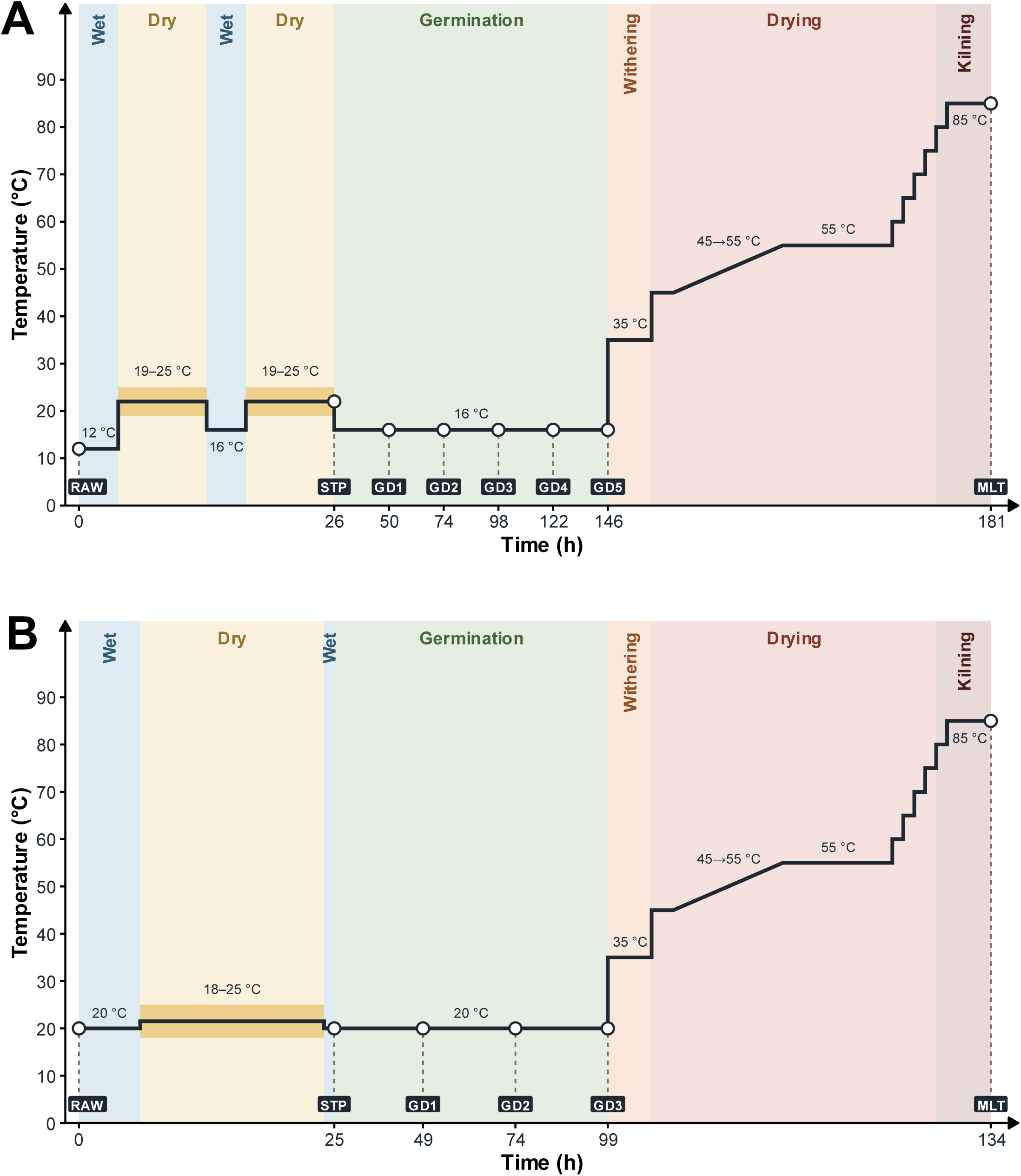
Temperature-time profiles and sampling points of the standard industrial (IMR) and accelerated (AMR) malting regimes. For IMR (**A**), steeping comprised two wet steeps (1^st^ wet steep at 12 °C; 2^nd^ wet steep at 16 °C; both lasting 4 h), each followed by a 9 h dry rest at 19–25 °C, after which the barley grains were germinated at 16 °C for 120 h (∼5 days). For the AMR (**B**), steeping comprised a first wet steep at 20 °C for 6 h, a dry rest at 18–25 °C for 18 h and a second wet steep at 20°C for 1 h, followed by germination at 20 °C for 74 h. In both regimes, the kilning programme was kept constant. Green malts were withered at 35 °C for 4 h and then dried: 2 h at 45 °C, a 10 h linear ramp from 45 to 55 °C, 10 h at 55 °C, and 1 h steps at 60, 65, 70, and 75 °C. Finally, the green malt was kilned at 80–85 °C for 5 h. For both plots, solid lines indicate the temperature programme, while the shaded bands show the temperature ranges during the dry rests, with the line drawn at the midpoint of each range. Labelled markers show the sampling time points: barley before imbibition (RAW); chitted barley at the end of steeping (STP); germination days 1–5 (GD1–GD5, IMR) and 1–3 (GD1–GD3; AMR); finished malt (MLT, kilned malt from which rootlets had been removed).

### 2.2. Sample collection

Samples were collected at the following time points: ungerminated grain before imbibition (RAW), at the end of steeping (STP), after germination days (GD) 1 to 5 and after kilning (MLT). In total, eight time points were sampled for IMR (RAW, STP, GD1–GD5, MLT) and six for AMR (RAW, STP, GD1–GD3, MLT). All samples for each variety and time point were taken from the same meshed stainless-steel bucket, representing the biological replicate per variety × regime combination. Samples for enzyme analysis and metagenomics were vacuum-packed and stored at –20 °C, whereas samples for metaproteomics were stored at –80 °C (all subsamples from the same bucket).

### 2.3. Analysis of acrospires’ growth performance

The MEBAK protocol for acrospire development in malt (MEBAK online, 2016) was employed for the evaluation of acrospire development along the germination timeline. For each sampling time point (GD1–GD5 in IMR, and GD1–GD3 in AMR), 100 barley kernels were randomly selected from each of three replicates per barley variety and boiled for 1 min in a 20 % (wt/vol) copper(II)sulphate solution (Cat. No. 8175.1, Carl Roth, Karlsruhe, Germany), incubated in the same solution for 30 min and afterwards washed with reverse osmosis water. The kernels were then kept at –20 °C until further analysis. The ratio of the acrospire length to kernel length (as measured manually by a ruler) was calculated for each grain for classification and assessing the uniformity of germination as described in the MEBAK protocol R-200.15.733 (revised 2016) (MEBAK online, 2016).

### 2.4. Malt quality and enzymatic analysis

Standard malt quality parameters were determined by Albert Maltings (Ruisbroek, Belgium) using the following methods: moisture content (EBC, 4.2); total protein content (EBC, 4.3.1); fine/coarse extract (EBC 4.5.2); soluble protein (Cat. No. 530-001 issue 122211/MH/99270407, Skalar, Breda, Netherlands); free amino nitrogen, FAN (Cat. No. 149-203 issue 122211/MH/99270407, Skalar); friability (EBC 4.15); final attenuation (EBC 8.6.1); colour (EBC 4.7.2); pH (EBC 4.5.1); viscosity (EBC 4.8); filtration (EBC 4.5.1); and diastatic power (Cat. No. 308-406 issue 122211/MH/99270407, Skalar). Grain moisture content was monitored during steeping and germination using a rapid moisture analyser (HE73, Mettler Toledo, Greifensee, Switzerland), and acrospire growth was determined according to MEBAK method R-200.15.733 (MEBAK online, 2016).

Enzyme activities and carbohydrate contents were determined with the following Megazyme kits (Dublin, Ireland), according to the manufacturer’s protocols: endo-β-glucanase (Cat. No. T-BGZ-200T, β-glucazyme tablets, lot: 110901); α- and β-amylase (Malt Amylase assay kit, Cat. No. K-MALTA); endo-protease (Cat. No. T-PRAK-200T, SKU:700005108, Protazyme AK tablets, lot:140401); β-glucan (β-Glucan assay kit, Cat. No. K-BGLU, SKU: 700004269); endo-1,4-β-xylanase (T-XAX, SKU 7000051100); and starch (Total Starch assay kit, Cat. No. K-TSTA-100A, SKU:700004351). Total arabinoxylan and water-extractable arabinoxylan contents were determined based on the gas chromatography method described by Gebruers, Courtin and Delcour (Gebruers et al., 2009). Malt quality parameters of IMR and AMR were compared using the Student’s t-test, as implemented in the rstatix package (Kassambara, 2026) in R (version 4.5.1;) (R Core Team, 2023) and resulting *P*-values were adjusted using the Holm-Bonferroni method (Holm, 1979) with a *P-*value cut-off of 0.05. The data analysis scripts for malt quality and enzymatic analyses, along with their raw data, are available from Zenodo under the URL https://doi.org/10.5281/zenodo.23890431.

### 2.5. Assessment of the lautering performance of malt

IMR and AMR malt samples were coarsely milled (1.0 mm) using a DLFU disc mill (Bühler company, Braunschweig, Germany) and the ground malts were mashed according to the EBC congress mash protocol (EBC 4.5.1, (EBC Analytica, 2004)) using a laboratory-scale mashing unit (LB8, Lochner, Labor und Technik GmbH, Berching, Germany). Lautering performance was evaluated as described by (Evans et al., 2011) by measuring the weight of wort filtered through a 270 mm Whatman folded filter paper (Grade 597 ½; Cat. No. 10311852, Cytiva, Marlborough, MA, USA) after 25 min.

### 2.6. Microbial DNA extraction for metagenomic analysis

DNA for metagenomic analysis was extracted from microbes colonising the surfaces of barley grains. Approximately 50 g of grains were immersed in 100 mL phosphate-buffered saline (137 mM sodium chloride (Cat. No.: 15628170, Fisher Scientific, Pittsburgh, PA, USA), 2.7 mM potassium chloride (Cat. No. 1.04936.0500, Merck KGaA, Darmstadt, Germany), 10 mM disodium hydrogen phosphate (Cat. No. 1.06580.1000, Merck KGaA), 1.8 mM potassium dihydrogen phosphate (Cat. No. 1.04873.1000, Merck KGaA)) and washed on an orbital shaker (PSU-20i, Grant Instruments, Royston, UK) at 200 rpm for 1 h at room temperature (RT). The wash solution was then centrifuged at 3000 × g for 30 min (4 °C) using a refrigerated centrifuge (5910 Ri, Eppendorf, Hamburg, Germany). The supernatant was discarded and the resulting pellets were resuspended in 2.5 mL Carlson lysis buffer (2% (wt/vol) cetyltrimethylammonium bromide (Cat. No. 9161.1, Carl Roth), 100 mM Tris-base (Cat. No. 93350, Sigma-Aldrich, St. Louis, MO, USA), 1.37 M sodium chloride (Cat. No. 15628170, Fisher Scientific), 20 mM disodium ethylenediaminetetraacetate dihydrate (Cat. No. 324503-100GM, MilliporeSigma, Burlington, MA, USA), 1% (wt/vol) polyethylene glycol (PEG) 8000 (Cat. No. BP233-100, Fisher Scientific) and 7 μL β-mercaptoethanol (Cat. No. M3148, Sigma-Aldrich)) and incubated at 65 °C in a temperature-controlled water bath for 1 h to lyse the cells. An equal volume of chloroform:isoamyl alcohol (24:1, vol/vol; Cat. No. A1935.0500, VWR Life Science, Radnor, PA, USA) was added to the lysates and the tubes were gently inverted ten times to mix the phases. The samples were then centrifuged at 3000 × g for 10 min (4 °C), and the upper aqueous phase containing DNA was transferred to a new 15 mL conical tube and the chloroform:isoamyl alcohol enrichment was repeated once to improve DNA quality and purity. The aqueous phase was then transferred to a 15 mL DNA LoBind tube, and a 0.1× volume of 3 M sodium acetate (i.e., 1⁄10 of the volume of the aqueous phase collected after the second chloroform: isoamyl alcohol wash; Cat. No. 1062680250, Merck KGaA) was added, followed by a 1× volume of room-temperature isopropanol (i.e., the volume of the aqueous phase including the added sodium acetate; Cat. No. P/7500/15, Fisher Scientific). The tubes were gently inverted 3–5 times and stored at –20 °C overnight to precipitate the DNA. The next day, the samples were centrifuged at 3000 × g for 10 min (4 °C), the supernatant was discarded, and the resulting DNA pellet was washed twice with 4 mL of 70% (vol/vol) ethanol (Cat. No. E/06665DF/17, Fisher Scientific). After air-drying to evaporate the ethanol, the DNA pellet was dissolved in 500 μL Tris-EDTA buffer (10 mM Tris-HCl (Cat. No. 93350, Sigma-Aldrich), 1 mM EDTA (Cat. No. 324503-100GM, Sigma-Aldrich), pH 8.2), 1 μL RNase A (20 mg/mL; Cat. No. T3018L, New England Biolabs, Ipswich, MA, USA) was added, and the samples were incubated at 37 °C for 1 h. Next, 2.5 μL of Proteinase K (20 mg/mL; Cat. No. 19131, QIAGEN GmbH, Hilden, Germany) was added, and the samples were incubated in a water bath at 55 °C for 15 min. To remove the added enzymes, the samples were then washed twice with chloroform: isoamyl alcohol as described above, and the final aqueous phase (approx. 300 μL) was transferred to a new 1.5 mL DNA LoBind tube to which a 0.1× volume of 3 M sodium acetate, followed by a 1× volume of ice-cold (–20 °C) isopropanol, were added. The contents were mixed by gently inverting the tubes 3–5 times and stored at –20 °C overnight to precipitate the DNA. The next day, the samples were centrifuged at 3000 × g for 10 min (4 °C) (CD-3124R, Phoenix Instruments, Garbsen, Germany), the supernatant was discarded, and the DNA pellet was washed twice with 1.5 mL of cold (4 °C) 70% (vol/vol) ethanol. After the second ethanol wash, the supernatant was discarded, the DNA pellet was air-dried and resuspended in 20–50 μL Tris-EDTA buffer (depending on the pellet size). DNA integrity was evaluated by agarose gel electrophoresis (see **Supplementary Figure 1**). Gels were prepared by dissolving 0.4 g agarose (Cat. No. CLOO.2762.0100, Chem-Lab NV, Zedelgem, Belgium) in 100 mL of Tris-acetate EDTA buffer (40 mM Tris (Cat. No. 93350, Sigma-Aldrich), 20 mM acetic acid (Cat. No. 20104.298, VWR Chemicals, Radnor, PA, USA), 1 mM EDTA (Cat. No. 324503-100GM, Sigma-Aldrich)) by boiling in a microwave, and GelRed was added at a 0.5× concentration from a 10,000× stock (Cat. No. BIOT41003, Biotium, Fremont CA, USA). An aliquot of each sample (25 ng DNA) was mixed with 2 μL of 6× purple loading dye (Cat. No. B7025S, New England Biolabs), adjusted to 12.5 μL with ultrapure water and electrophoresed at 55 V for 120 min. DNA was visualised using an iBright FL1500 imager (Cat. No. A44115, Thermo). DNA purity was assessed by A_260_/_A280_ and A_260_/_A230_ readings using a NanoPhotometer (NP80, Implen, Munich, Germany) and DNA concentration was determined using a Quantus fluorometer (Cat. No. E6150, Promega Corporation, Madison, WI, USA) applying the QuantiFluor®ONE dsDNA System (Cat. No. E4870, Promega) according to the manufacturer’s protocol.

#### 2.6.1. Metagenomic DNA purification and size-selection

Long DNA fragments were enriched by size-selection based on solid-phase reversible immobilisation (SPRI), as described by Stortchevoi and colleagues (Stortchevoi et al., 2020; Stortchevoi and Levine, 2024). Briefly, the required volumes of SPRI beads (Hieff NGS®DNA selection beads, 12601ES56, YEASEN, Shanghai, China), determined from the target size selection range of > 4 kbp, (Stortchevoi and Levine, 2024) and the DNA yield of each sample, were exchanged in nuclease-free water (Cat. No. PD092, Omega Bio-tek, Norcross, GA, USA) using two rounds of magnetic binding and washing. Metagenomic DNA of each sample was conditioned by dilution to 105 μL in Tris-EDTA buffer (10 mM Tris-HCl (Cat. No. 93350), 1 mM EDTA (Cat. No. 324503-100GM, Sigma-Aldrich), pH 8.2), and 100 μL was added to the corresponding volume of exchanged beads, mixed by pipetting 12–15 times through a 200 μL wide-bore pipette tip (Cat. No. 19917744, Fisher Scientific) and briefly spun down (mySPIN™6 Mini Centrifuge, Fisher Scientific). To each bead/DNA mixture, 65 μL of the customised binding buffer (NaPEG; 750 mM NaCl (Cat. No. 15628170, Fisher Scientific); 20% (wt/vol) PEG 8000 (Cat. No. BP233-100, Fisher Scientific)) was added, mixed by gently pipetting 12–15 times through a 200 μL wide-bore tip, briefly spun down and incubated at room temperature for 10 min. The bead pellet was then collected and the supernatant discarded. The beads were washed twice with freshly prepared 80%(vol/vol) ethanol, air-dried for 5–10 min, and then DNA was eluted with 10–30 μL Tris-EDTA buffer (10 mM Tris-HCl (Cat. No. 93350), 1 mM EDTA (Cat. No. 324503-100GM, Sigma-Aldrich), pH 8.0). DNA concentration and purity were determined using a Quantus fluorometer (Promega Corporation) and a NanoPhotometer (NP80, Implen), respectively, as described in section 2.6. The remaining size-selected DNA was checked using a 0.4% agarose gel image (see **Supplementary Figure 2**) and was briefly stored at –20 °C until library preparation for sequencing.

### 2.7. DNA library preparation for Nanopore longread sequencing

Whole-metagenome sequencing was performed using samples from four barley varieties (crop year 2021) germinated by IMR or AMR (see **Figure 1A** and **1B**), including 19 samples from crop year 2023 of the Planet variety germinated using IMR (n = 173 in total). The 2021 crop samples were subsample from the same lot (Bucket) as those used for enzymatic and metaproteomics analysis. Libraries were prepared in two cohorts using a Rapid Barcoding Kit 96 V14 (Cat. No. SQK-RBK114.96, Oxford Nanopore Technologies, Oxford, UK) according to the manufacturer’s manual, with two minor deviations: the DNA library was eluted from the AMPure XP beads (Cat. No. A63880, Beckman Coulter, Brea, CA, USA) at 37 °C instead of room temperature and bovine serum albumin was omitted from the flow cell priming mix. Library DNA concentration was determined using a Quantus fluorometer with the QuantiFluor® ONE dsDNA assay kit as described in section 2.6. Sequencing was performed on a MinION Mk1B device (Oxford Nanopore Technologies) using two R10.4 flow cells (FLO-MN114), with 77 samples multiplexed in the first run (BarleyMetagenomics_R10_Run01), and 96 in the second (BarleyMetagenomics_R10_Run02). To enrich the sequencing of metagenomic DNA, barley host DNA was depleted by adaptive sampling in both runs using the barley reference genome downloaded from NCBI (Genome assembly MorexV3_pseudomolecules_assembly; genome accession: GCA_904849725.1). Reads were generated for 72 h with the high-accuracy (HAC) model in MinKNOW (version 25.03.9).

### 2.8. Metagenomic data processing

After sequencing, the raw nanopore reads were re-basecalled with Dorado (version 1.0.0, https://github.com/nanoporetech/dorado), in super-accuracy mode (SUP) using the dna_r10.4.1_e8.2_400bps_sup@v5.2.0 model. Downstream processing was performed on two versions of the same reads: (1) demultiplexed reads, for which SUP-basecalled reads were demultiplexed for per-sample quantification, and (2) non-demultiplexed reads, comprising the combined reads of all samples sequenced in this study for microbial identification. In both cases, adapters were removed using Dorado; the resulting per-sample BAM files were deposited on the NCBI Sequence Read Archive (SRA) under BioProject ID **PRJNA1541012**. The BAM files of both versions were converted to FASTQ files using SAMtools (Danecek et al., 2021) and human DNA contamination was removed by aligning the reads to the *Homo sapiens* genome (“hsa” FASTA file Homo_sapiens.GRCh38.dna.primary_assembly, downloaded from Ensembl) using CLEAN (version 1.1.2) (Lataretu et al., 2025). The decontaminated FASTQ files were then filtered with Chopper (version 0.8.0) (De Coster and Rademakers, 2023) to retain only reads with a Q-score ≥ 9 and a minimum read length of 500 bp. Nanoplot (version 1.43.0) and NanoStat (version 1.6.0) (De Coster et al., 2018) were used to diagnose the quality of the sequencing data, evaluate the efficiency of the DNA extraction process and also check the effectiveness of the applied filters in removing low-quality reads (for the diagnostic plots, see **Supplementary Figures 3 and 4**).

#### 2.8.1. DIAMOND and MEGAN analysis

All reads passing the Chopper quality filter were aligned against the NCBI non-redundant (nr) protein database (downloaded from ftp://ftp.ncbi.nih.gov/blast/db/FASTA/nr/gz_on_April_9_2025) using DIAMOND (version 2.1.9.163, with default alignment settings) following the procedure described by Bağci, Patz and Huson (Bağcı et al., 2021). The resulting DIAMOND alignment archive (DAA) files were “meganized” (i.e., all sequences and alignments in the DAA file were subjected to taxonomic and functional binning (Bağcı et al., 2021)) using the command-line tool daa-meganizer (version 6.25.9, with default settings). The “meganized” DAA files were analysed in the graphical user interface of MEGAN (‘MetaGenome ANalyser’) Community Edition (version 6.25.10) to obtain the taxonomic profiles at the genus and species levels. Read alignment and meganization of the aligned reads were performed on Linux using the KU Leuven High Performance Computing infrastructure.

### 2.9. Statistical analysis of metagenomic data

#### 2.9.1. Data preparation for alpha and beta diversity analysis

Species-level read counts for all IMR and AMR samples obtained using the DIAMOND-MEGAN pipeline were analysed in R (version 4.5.1) (R Core Team, 2023). Species with zero reads in all samples were discarded. For downstream analyses, each biological replicate followed over time was treated as one experimental unit (BucketID), giving 24 BucketIDs ( 2 regimes × 4 varieties × 3 replicates). To evaluate the effects of the regime on the barley-associated microbiome during germination, alpha and beta diversity analyses were performed on two datasets since germination lasted longer in IMR (GD1–GD5) than in AMR (GD1–GD3): 1) a dataset composed of time points shared in both regimes (STP and GD1– GD3), which allowed us to compare the regimes on the same germination day while accounting for time; and 2) a dataset composed of the last germination days of both regimes, when grain modification was optimum (IMR GD5 vs AMR GD3). As a quality control check, the sequencing depth (total species-level per sample) of both regimes was compared with a two-sided Wilcoxon rank-sum test, with an additional assessment of its association with the observed species richness using Spearman’s rank correlation (see **Supplementary Figure 5** for diagnostic plots).

#### 2.9.2. Alpha diversity

Microbial alpha diversity was assessed using observed species richness (the number of detected species), the Shannon index and the Gini-Simpson index. Due to the sensitivity of observed richness to the sequencing depth (Lundin et al., 2012; Schloss, 2024), richness was estimated by rarefaction (using the rarefy () function from the *vegan* R package) as the expected number of species in a random subsample of (N) reads, the smallest library size (Hurlbert, 1971). Shannon and Simpson indices were calculated from the full count data.

##### 2.9.2.1. Alpha diversity within a malting regime

Temporal changes in alpha diversity were investigated using linear mixed-effects models, with variety, time point and their interactions as fixed effects. Bucket identity /biological replicate was included as a random intercept to account for repeated sampling of the same replicate over time (lme4, version 2.0.6; (Bates et al., 2015)). Model assumptions were checked using residual diagnostics, including assessment of residual normality using quantile-quantile plots (see **Supplementary Figure 6**). Differences between varieties and sampling time points were evaluated using estimated marginal means and post-hoc pairwise comparisons, with *P*-values adjusted for multiple comparisons using the Benjamini-Hochberg method (Benjamini and Hochberg, 1995).

##### 2.9.2.2. Alpha diversity between malting regimes

Each alpha index was also analysed using linear mixed-effects models with regime, variety, time point, and their interactions as fixed effects, and the biological replicate (bucket identity) as a random intercept. Fixed effects were tested with the type III F tests using sum-to-zero contrasts and Satterthwaite’s approximation for denominator degrees of freedom (lmerTest version 3.2.1, (Kuznetsova et al., 2017)). Regime contrasts (AMR vs. IMR) were estimated from estimated marginal means (emmeans version 2.0.4, (Lenth and Piaskowski, 2026)) within each variety (averaged over time points), within each time point, and within each variety × time point combination. *P*-values were adjusted with the Benjamini-Hochberg procedure within each family of contrasts. Normality of residuals was checked with normal q-q plots. For comparisons using the end-of-germination dataset, alpha indices were analysed by two-way ANOVA (regime × variety). The overall regime contrast and the regime contrast within each variety were obtained from estimated marginal means; within-variety *P*-values were Benjamini-Hochberg adjusted across the four varieties. Statistically significant results were considered at *P* < 0.05.

##### 2.9.2.3. Taxonomic barley microbiome composition

Differences in community composition were tested by permutational multivariate analysis of variance (PERMANOVA; (Anderson, 2017)) on Bray-Curtis dissimilarities calculated on reads converted to relative abundances per sample (total-sum scaling), using the adonis2 function of the vegan package (version 2.7.6 (Oksanen, 2025); 999 permutations) with permutation schemes defined using the permute R package (version 0.9.10 (Simpson, 2026). For common time points between the two regimes, the model included regime, variety, time point and all interactions. To account for the repeated measurements on the same biological replicate over time, terms that vary between replicates (regime, variety and regime x variety) were tested with permutations restricted within time points. Time point and its interactions were tested by permuting samples within replicates. For AMR and IMR comparisons within each variety, the model included time point and regime × time point, with permutations restricted within time points. With three replicates per regime, permuting whole replicates would only allow 20 distinct arrangements (minimum *P* = 0.05), so the time-stratified scheme was used, and R^2^ were reported as the effect size. *P*-values were Benjamini-Hochberg adjusted across the four varieties. For assessment of the regime effect at each time point and the end of germination, the model consisted of variety × regime effects, with unrestricted permutations run separately for each time point. Regime *P*-values were Benjamini-Hochberg adjusted. For the end-of-germination dataset, regime was also tested within each variety (3 vs 3 samples, complete enumeration of permutations; minimum attainable *P* = 0.05). Multivariate homogeneity of dispersions was tested with PERMDISP using the betadisper function followed by permutest with 999 permutations. This was done for regime and for regime × variety groups (pairwise) at the common time points, and for regime in each per-time-point and end-of-germination data set. The microbial community composition was visualised using principal coordinate analysis on the Bray-Curtis distances of the respective sample set. All scripts used for the alpha and beta diversity analyses are available from Zenodo via the URL https://doi.org/10.5281/zenodo.24121796, and the corresponding metagenomics raw data (in BAM output files per sample) can be accessed via the following SRA link: https://www.ncbi.nlm.nih.gov/sra/PRJNA1541012.

#### 2.9.3. Differential abundance testing

Analysis of Compositions of Microbiomes with Bias Correction 2 (ANCOM-BC2) was used to test for differentially abundant taxa between the two malting regimes in 1) all common time points, 2) at the end of germination, 3) at each specific sampling time point and 4) within a single barley variety, all performed at the species level. Taxa were considered robustly different if they were reported as differentially abundant (Holm-adjusted *P*-value < 0.05) and additionally passed the sensitivity test for pseudo-count addition (Lin and Peddada, 2024). Differentially abundant taxa were visualised as log fold changes (AMR vs IMR) at the species level using ggplot2 barplots. Variables included in the models were the same as those described for beta diversity analysis in section 2.9.2.3.

##### 2.9.3.1. Software used for metagenomics data analysis and data visualisation

All metagenomics statistical analyses were performed in R (version 4.6.1; (R Core Team, 2023)) with vegan (version 2.7–6; (Oksanen, 2025), permute (version 0.9–10; (Simpson, 2026)), lme4 (version 2.0–6; (Bates et al., 2015)), lmerTest (version 3.2–1; (Kuznetsova et al., 2017)), emmeans (version 2.0.4; (Lenth and Piaskowski, 2026)) and tidyverse (version 2.0.0; (Wickham et al., 2019)). Figures were produced using ggplot2 (version 4.0.3, (Wickham, 2016)) and patchwork (version 1.3.2, (Pedersen, 2025)). All the analysis scripts used are available from Zenodo under the URL https://doi.org/10.5281/zenodo.24121796.

### 2.10. Microbial cell and protein extraction

Germinated barley grains (approx. 3 g) were thawed at room temperature and submerged in 3 mL of 137 mM sodium chloride solution (Cat. No. 15628170, Fisher Scientific), and ultrasonicated in a water bath (Cat. No. 960733205, SONOREX DIGITEC DT 52, BANDELIN electronic GmbH, Germany) at room temperature for 15 min. The wash was decanted into a 50 mL conical tube, and the grains were washed a second time. The combined washes were mixed with 4 volumes of ice-cold (–20 °C) 20% (wt/vol) trichloroacetic acid (Cat. No.10775151, Fisher Scientific B.V., Boulevard Saint-Lazare, Belgium) in 80% (vol/vol) acetone (Cat. No. 10098181, Fisher Scientific B. V., Boulevard Saint-Lazare, Belgium) and incubated at –20 °C overnight to ensure complete protein precipitation. The precipitate was pelleted by centrifugation at 8,960 × g for 20 min (–10 °C), the supernatant was discarded, to the pellet 1 mL ice-cold (–20 °C) acetone was added and the samples were incubated for 30 min at –20 °C with intermittent vortexing every 10 min. The samples were then centrifuged at 8,960 × g for 20 min (– 10 °C), the supernatant was discarded, and the samples were air-dried for 10–15 min. The samples were then resolubilised in 8 M urea (Cat. No. 1.08487.0500, Merck KGaA, Darmstadt, Germany) in 50 mM ammonium bicarbonate (AmBiC, Cat. No. 09830-500G, Sigma-Aldrich) by incubation for 1 h at room temperature, assisted by ultrasonication in a water bath (DT 52, Bandelin, Berlin, Germany) for 15 min. Afterwards, the samples were transferred to 1.5 mL Protein LoBind tubes (Eppendorf) and their protein concentration was determined using Bicinchoninic Acid (BCA) Protein Assay (Cat. No. 23225, Thermo Fisher Scientific, Waltham, MA, USA) according to the manufacturer’s manual.

### 2.11. Sample preparation for proteome analysis

Proteins (50 μg per sample) were reduced and alkylated for 45 min at room temperature in the dark with 10 mM tris(2-carboxyethyl) phosphine (TCEP, Cat. No. C4706-2G, Sigma-Aldrich) and 25 mM chloroacetamide (CAA, Cat. No. C0267-100G, Sigma-Aldrich). The samples were diluted with 50 mM AmBiC to a urea concentration of 2 M, and 1 μg of trypsin (1 μg/μL in 50 mM AmBiC; Cat. No. 90057, Thermo Fisher) was added. Proteins were digested for 21 h at 24 °C, and the resulting peptide mixture was acidified with trifluoroacetic acid (TFA, Cat. No. 1.08262.0025, Merck) to a final concentration of 1% (vol/vol) to stop the enzymatic reaction. Ten micrograms of the acidified tryptic peptides were desalted on home-made C18 columns (SPE discs, 3M, Saint Paul, MN, USA, Cat. No. 66883-U), as described previously (Jadav et al., 2024) and freeze-dried (Alpha 2–4 LSC, Martin Christ Gefriertrocknungsanlagen GmbH, Osterode, Germany). Peptides were resuspended in 71.4 mM HEPES (pH 8.5, Cat. No. 54457-10G-F, Sigma-Aldrich), ultrasonicated in a water bath for 15 min, and labelled with 4 µL of the corresponding TMTpro 18-plex reagent (Cat. No. A44521 and Cat. No. A52046, Thermo Fisher Scientific) reconstituted to 13 mM in anhydrous acetonitrile (Cat. No. 271004, Sigma-Aldrich) as described in (Zecha et al., 2019). After 2 h of incubation at room temperature, the reaction was quenched by adding hydroxylamine (Cat. No. 467804-10ML, Sigma-Aldrich) to a final concentration of 0.5% (wt/vol), followed by incubation at room temperature for 15 min. An aliquot (1 μL) of each quenched reaction was pooled and freeze-dried. The TMT-labelled peptides were resuspended in 2% (vol/vol) acetonitrile (Cat. No. 1.00029.1000, Merck) and 0.1% (vol/vol) formic acid (Cat. No. 5.33002.0050, Merck), ultrasonicated in a water bath for 15 min, transferred to autosampler vials and submitted to the VIB Proteomics Core (Ghent, Belgium) for LC-MS/MS analysis.

### 2.12. Liquid chromatography tandem mass spectrometry (LC-MS/MS)

An aliquot of 10 µL (approx. 4–5 μg) of the tryptic peptides was analysed by LC-MS/MS on an UltiMate 3000 RSLCnano system (Thermo Fisher Scientific) coupled to a Q Exactive HF mass spectrometer (Thermo Fisher Scientific) equipped with a pneu-Nimbus dual ion source (Phoenix S&T, Chadds Ford, PA, USA). Peptides were trapped at 20 µL/min for 2 min in loading solvent A (0.1% (vol/vol) trifluoroacetic acid in water/acetonitrile, 99.5:0.5, vol/vol) on a PepMap™Neo Trap column (Thermo Fisher Scientific, 300 µm internal diameter, 5 µm beads) and separated on a 250 mm Odyssey Ultimate column (1.7 µm C18 resin, 75 µm inner diameter; IonOpticks, Melbourne, Australia) kept at 45 °C. Peptides were eluted with linear gradients reaching 33% MS solvent B (0.1% (vol/vol) formic acid in water/acetonitrile, 20:80, (vol/vol)) at 135 min, 55% at 155 min, and 70% at 160 min, at a constant flow rate of 300 nL/min, followed by a 5-min wash at 70% MS solvent B and re-equilibration with MS solvent A (0.1% (vol/vol) formic acid in water). The mass spectrometer was operated in data-dependent mode, automatically switching between MS and MS/MS acquisition for the 12 most abundant ion peaks per MS spectrum. Full-scan MS spectra (375–1500 *m/z*) were acquired at a resolution of 60,000 in the Orbitrap mass analyser after accumulation to a target value of 3 million. The 12 most intense ions above a threshold of 13,000 were isolated (isolation window of 1.5 *m/z*) and fragmented at a normalised collision energy of 33% after filling the trap to a target value of 100,000 for a maximum of 80 ms. MS/MS spectra (fixed first mass 120 *m/z*) were acquired at a resolution of 45,000 in the Orbitrap analyser. The polydimethylcyclosiloxane background ion at 445.120028 Da was used for internal calibration, and QCloud was used to monitor the longitudinal performance of the instrument (Chiva et al., 2018).

### 2.13. Construction of metaproteome protein sequence database

The microbial taxonomy identifiers at the genus and species levels detected by metagenomics were used to download the corresponding (pan)proteome FASTA files from the UniProt Knowledgebase (UniProtKB; retrieved on 26^th^ June 2025), which were concatenated into a single FASTA file via the command line. The *Hordeum vulgare* TrEMBL FASTA file (UniProtKB, downloaded 6^th^ February 2023) and the common Repository of Adventitious Proteins (cRAP) database of common laboratory contaminants (The Global Proteome Machine, ftp://ftp.thegpm.org/fasta/cRAP) were added, giving a total of 2,253,236 protein sequences. Decoy sequences were then added using FragPipe (version 23.1) to complete the metaproteome protein sequence database. The script to download the relevant FASTA files is available from Zenodo under the URL: https://doi.org/10.5281/zenodo.23995407.

### 2.14. Data processing and analysis of metaproteomic data

Mass spectrometry raw files were converted to centroided mzML files using MSConvert (version 3.024310) from the ProteoWizard software suite (Chambers et al., 2012) and searched with MSFragger within the FragPipe computational platform (version 23.1) (Kong et al., 2017) against the *in-house* constructed FASTA database described in section 2.13. The TMT-18 workflow was used for peptide identification, with carbamidomethylation of cysteine and TMT labelling of the N-terminus as fixed modifications, and methionine oxidation, peptide N-terminal acetylation and TMT-labelling of lysine as variable modifications. The metaproteomics database generation module was enabled, with the minimum number of unique peptides set to 1 and the host name set to *Hordeum vulgare*. Mass calibration and parameter optimisation were disabled to prevent out-of-memory errors. Peptide spectrum matches (PSMs) were rescored using MSBooster (Yang et al., 2023) (version 1.3.17) and validated using Percolator (The et al., 2016), with false discovery rates set at 5% at both the peptide and protein levels. Isobaric labelling quantification, was performed with IonQuant (Yu et al., 2020) with the label type set to “TMT-18”, normalisation set to “none,” and the log2 transformation of the intensities was disabled. All other parameters were kept at the default settings of FragPipe’s TMT18 workflow. The FragPipe output pep.xml files were then rescored after the database search using MS^2^Rescore (Buur et al., 2024; Malliet et al., 2026) (version 3.1.5). The details of the configuration settings used for MS^2^Rescore search can be found in **Supplementary Figure 7**. The resulting .pin files from MS^2^Rescore were rescored separately using Percolator (The et al., 2016) (version 3.07.1, default settings) and only PSMs with a q-value (false discovery rate) ≤ 0.05 were retained. Unique peptides were imported into the Unipept web application (version 6.3.4, https://unipept.ugent.be/, based on UniProt 2025.04) for taxonomic and functional annotation. To increase the specificity of peptide annotation in Unipept, a targeted protein reference database from the species identifiers obtained by metagenomics was constructed in Unipept using its custom database function. Single peptide analysis of each metaproteomic MS run was performed in Unipept with 1) equating isoleucine (I) with leucine (L) and 2) advanced missed-cleavage handling enabled. The resulting taxonomic matches (peptide-taxon tables) were downloaded from Unipept and filtered using in-house R scripts to retain only peptides matched to bacterial and fungal taxa. The data analysis scripts are available from Zenodo: https://doi.org/10.5281/zenodo.23995407.

### 2.15. Statistical analysis of Metaproteomics data

TMT reporter ion intensities were extracted from the centroided mzML files in R (version 4.5.2) using in-house scripts adapted from a previous study (Khanam et al., 2021). All extracted TMT reporter intensities were transformed by variance stabilising normalisation (VSN) (Huber et al., 2002) (see **Supplementary Figures 7A–8B** for diagnostic plots) using the vsn R package (v3.80.0) and differentially abundant peptides between two selected groups (contrasts), were identified using linear models and moderated t-statistics based on an empirical Bayes method, as implemented in the R package limma (v.36.3) (Ritchie et al., 2015). Enrichment of Gene Ontology (GO) terms was tested using Fisher’s exact test and q-values were calculated to control the false discovery rate (FDR) (Storey et al., 2024). GO terms with q-value < 0.05, were considered significantly overrepresented in the gene set relative to the background. All MS raw data, search results and metadata have been deposited to the ProteomeXchange Consortium via the PRIDE (Perez-Riverol et al., 2025) partner repository with the dataset identifier PXD086227, and the data analysis scripts are available from https://doi.org/10.5281/zenodo.23995407.

## 3. Results

### 3.1. Malt Production and Sampling

The study setup is shown in **Figure 1**. Four barley varieties (Fandaga (FDG), Faro (FAR), Fantex (FTX) and Planet (PLT)) were malted in triplicate using the programme parameters shown in **Figure 1A and 1B** (for grain moisture contents monitored during germination, see **Supplementary Table 1**). During malt production, samples were taken from raw barley (before imbibition), after steeping, and every 24 h thereafter for three germination days in AMR and five in IMR, with a final malt sample taken after kilning.

### 3.2. Chemical-Technical and Enzymatic Analysis of IMR and AMR Malts

#### 3.2.1. Amylolytic enzymes and starch degradation

AMR resulted in a higher residual starch content (**Figure 2A**), which we attribute to the shorter germination time and the lower α- and β-amylase activities (see **Figure 2B and C**). Surprisingly, Fandaga already showed α-amylase activity in raw barley kernels which is generally considered a result of pre-harvest sprouting (reviewed in (Orth et al., 2025)). However, we exclude pre-harvest sprouting here as it causes rapid losses of barley viability (Moor, 1987) which is not evident in the Fandaga malting results. Although the germinative energy of Fandaga (87.5%) was lower than that of Planet (97.5%), Fantex (99.5%) and Faro (99.0%) (**Figure 2D**), Fandaga generally germinated faster (based on acrospire length distribution) than the other varieties under both regimes (**Supplementary Figure 9)** and even produced the highest number of “huzzars” (kernels with an acrospire length longer than the kernel) in the last germination days (**Supplementary Table 2**). In addition, the very high percentage of friability and low wort viscosity indicate strong cytolysis (**Figure 4A and 4B**), which argues against a high proportion of pre-harvest-sprouted (and thus non-viable and unmodified) kernels. Moreover, the residual β-glucan content of Fandaga malt was not elevated relative to other varieties (**Figure 4C**), which is inconsistent with a loss of viability. We therefore speculate that the high initial α-amylase activity could be a result of late-maturity α-amylase production, as described for wheat (Mares and Mrva, 2008, 2014). As for the malt after kilning, β-amylase shows strong heat sensitivity, resulting in nearly equal activities in IMR and AMR malts (**Figure 2C**). The effect of kilning on α-amylase activity was variety dependent: Fandaga and Planet showed higher activity in IMR malt than in AMR malt (IMR vs AMR: Fandaga: 288.2 U/g vs 185.2 U/g, *P*_adj_ = 0.0147, Planet: 239.5 U/g vs 146.2 U/g, *P*_adj_ = 0.0025), whereas no difference was detected for Faro and Fantex (**Figure 2B**). From these results, the lower enzyme activities of AMR malts, together with their consistently lower diastatic power (**Figure 2E**), could adversely affect wort production. However, no defects in saccharification time were detected (< 10 min for all malts), indicating that the enzyme content of the here produced AMR malts is still sufficient for brewing.

**Figure 2.**
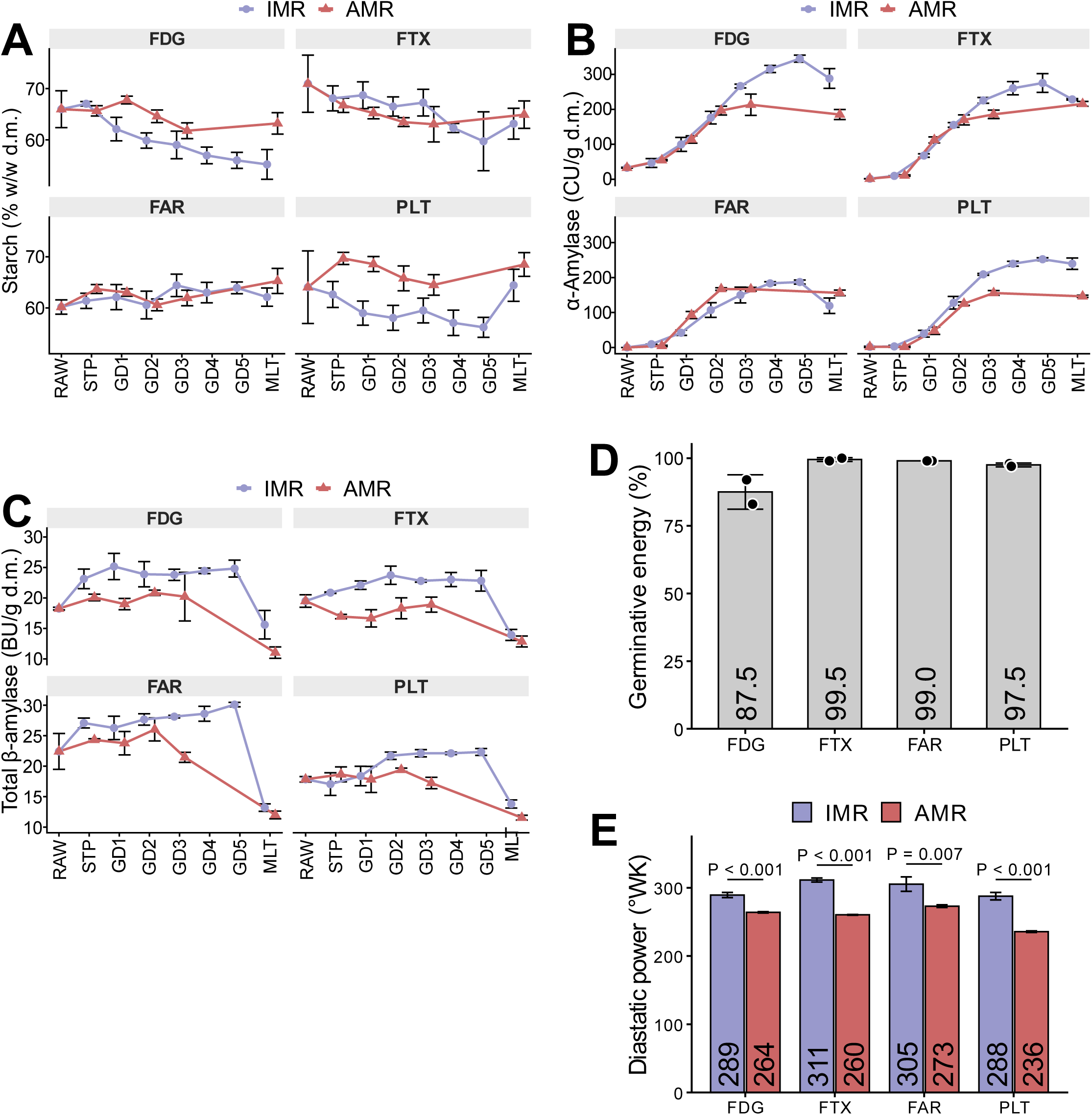
Variety-wise comparison of the amylolytic malt parameters of AMR and IMR malts, coupled with the germinative energy of the raw barley grains. **(A)** Temporal changes in the starch content during malting under AMR and IMR in each barley variety. **(B, C)** Temporal development of the α-amylase activity **(B)** and β-amylase activity **(C)** during malting under AMR and IMR for each barley variety. **(D)** Germinative energy of raw barley grains for each barley variety, determined using the Brewing Research Foundation method (BRF, EBC 3.6.2). **(E)** Comparison of the diastatic power of the AMR and IMR malts by variety. For each barley variety, *P*-values for differences between the two regimes were calculated using a Student’s t-test and Bonferroni-Holm adjusted to account for multiple testing. For all plots (A–E), data are shown as the mean ± standard deviation (n = 6 for A, n = 3 for **B**, **C**, and **E**, and n = 2 for **D**). FDG, Fandaga; FTX, Fantex; FAR, Faro; PLT, Planet; RAW (Ungerminated barley grain); STP (end of steeping or start of germination); GD (Germination day); MLT (finished malt; malt after kilning with rootlets removed).

#### 3.2.2. Proteolytic modification

In stark contrast to earlier research (Müller et al., 2014; Müller and Methner, 2015), AMR did not suppress proteolysis and led to even increased Kolbach indices (KI) in Fandaga (51.9% in IMR vs 56.8% in AMR, *P*_adj_ = 0.0036) (**Figure 3A**). Regardless of malting regime, most KI values were at or above the upper limit of the usually acceptable range of 35–45%, indicating potential to further shorten AMR malt production time. This was especially true for Fandaga, which also had the highest free amino nitrogen (FAN) concentration of all varieties (239 mg/L; **Figure 3B**), far exceeding the recommended minimum concentration of 150 mg/L (Beckerich and Denault, 1987). Further, endo-protease activity did not show a consistent difference between AMR and IMR; thus, we speculate that, in combination with earlier results (Müller et al., 2014; Müller and Methner, 2015) this is an effect of the individual variety, and a suppression of proteolysis by increased germination temperatures cannot be generalised.

**Figure 3.**
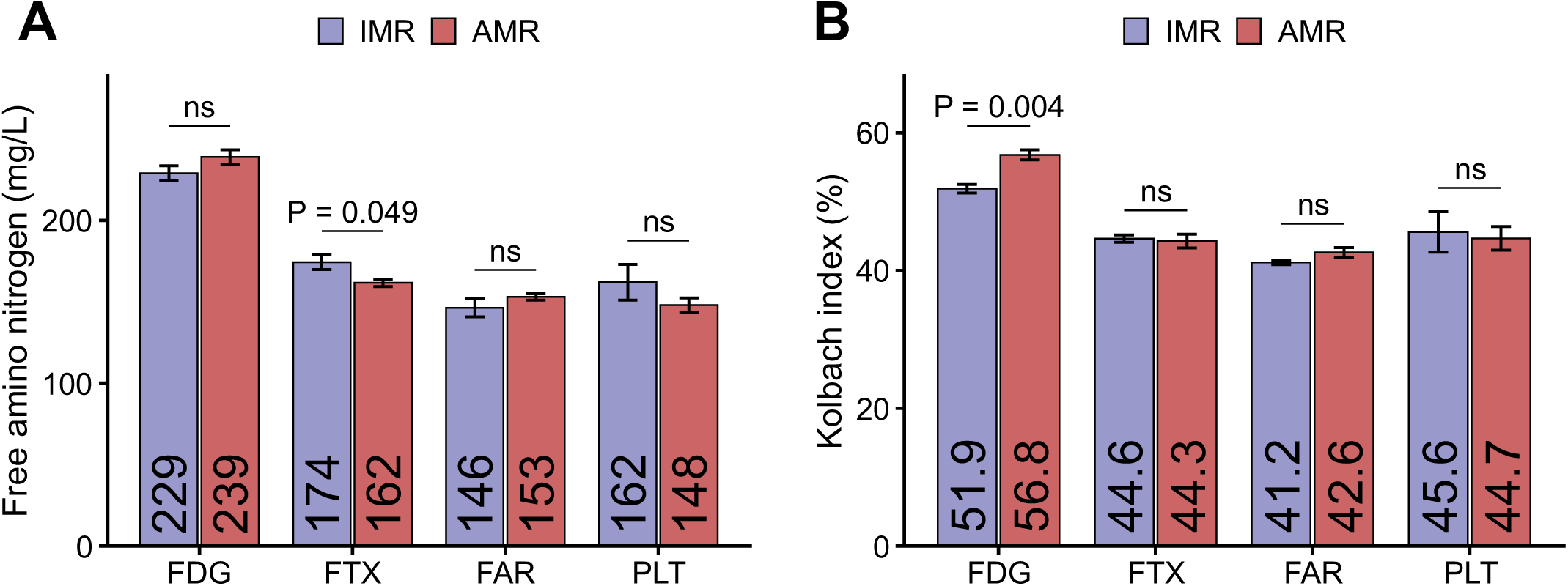
Variety-wise assessment of the level of proteolytic modification in AMR and IMR malts. **(A, B)** Comparison of the Kolbach index **(A)** and free amino nitrogen **(B)** of AMR and IMR malts by barley variety. For each quality attribute, we calculated *P*-values for differences between regimes within each variety using a Student’s t-test and Bonferroni-Holm adjustment for multiple testing (ns: adjusted *P*-value > 0.05). The bar plots show the mean values from three biological replicates, and the error bars indicate the standard deviation. FDG, Fandaga; FTX, Fantex; FAR, Faro; PLT, Planet.

#### 3.2.3. Cytolytic modification

We were also interested in how the two malting regimes influence cell wall degradation (cytolysis), as this reflects β-glucan degradation, an important indicator of expected mash filterability and wort viscosity. Consistent differences in malt friability were not detected between the two malting regimes. IMR resulted in higher friability in Fantex (AMR: 93.6%, IMR: 97.9%, *P*_adj_ = 0.001) and Planet (AMR: 87.8%, IMR: 96.7%, *P*_adj_ = 0.001), while AMR resulted in a higher friability in Faro (AMR: 89.3%, IMR: 79.4%, *P*_adj_ = 0.012) (**Figure 4A**). A friability of > 85% is generally considered acceptable (Jacob, 2016; Kreisz, 2009; O’Rourke, 2002), and as such most malts exceeded this value regardless of regime, with the exception of Faro with a friability of 79.4% after IMR. We therefore interpret friability in response to malting regime as an effect of barley variety. The friability differences were mirrored in wort viscosity, where a higher friability was associated with lower wort viscosity (**Figure 4B**), most likely owing to more extensive β-glucan degradation in more friable malts. Nevertheless, all malts had an acceptable wort viscosity below the recommended upper limit of 1.55–1.65 mPa·s (O’Rourke, 2002). Interestingly, β-glucan degradation progressed more slowly in IMR than in AMR (**Figure 4C**), but in most cases reached the same degree of more than 90% by the end of malting (except Faro in IMR). High germination temperatures have been shown to cause high initial enzyme production during seedling growth followed by a plateau, resulting in lower total enzyme levels at the end of malting than germination at low temperatures (Müller et al., 2014). Consistent with this, AMR resulted in lower enzyme activities, particularly of β-glucanase (**Figure 4D**), however, this is not mirrored in the residual β-glucan content (**Figure 4C**). Relating to this, the lautering performance of IMR and AMR malts was assessed in a wort filtration test. All AMR malts yielded markedly lower wort volumes after 25 min of filtration, with a reduction of 11.4% in Faro (*P*_adj_ = 0.0468), 13.0% in Fantex (*P*_adj_ = 0.0468), 22.2% in Fandaga (*P*_adj_ = 0.0468) and 17.4% in Planet (*P*_adj_ = 0.0337) (**Figure 5A**). These results contradict those of the other malt quality tests, and we tested high-molecular-weight arabinoxylan content, an additional source of wort viscosity (M. Gastl et al., 2021; Sadosky et al., 2002). However, arabinoxylan content showed no statistically significant differences in the resulting malt after kilning between both malting regimes (**Figure 4E**). To exclude a barley batch effect, we therefore conducted a repeat of the IMR and AMR in a different crop of Planet (2023). In contrast to the 2021 crop, the 2023 crop yielded higher filtration volumes with AMR (224 mL) than with IMR (169 mL, *P*_adj_= 0.0001, **Figure 5B**). AMR filtration volumes of the 2023 crop were also higher than IMR volumes of the 2021 crop (224 mL (AMR 2023) vs 177 mL (IMR 2021), *P*_adj_ = 2.08×10^-4^), whereas IMR volumes did not differ significantly between the 2023 and 2021 crops (*P*_adj_ = 0.478). We therefore conclude that AMR does not *per se* result in poorer lautering performance than IMR and turned our attention to the malting microbiome as a potential explanation for the differences in lautering performance.

**Figure 4.**
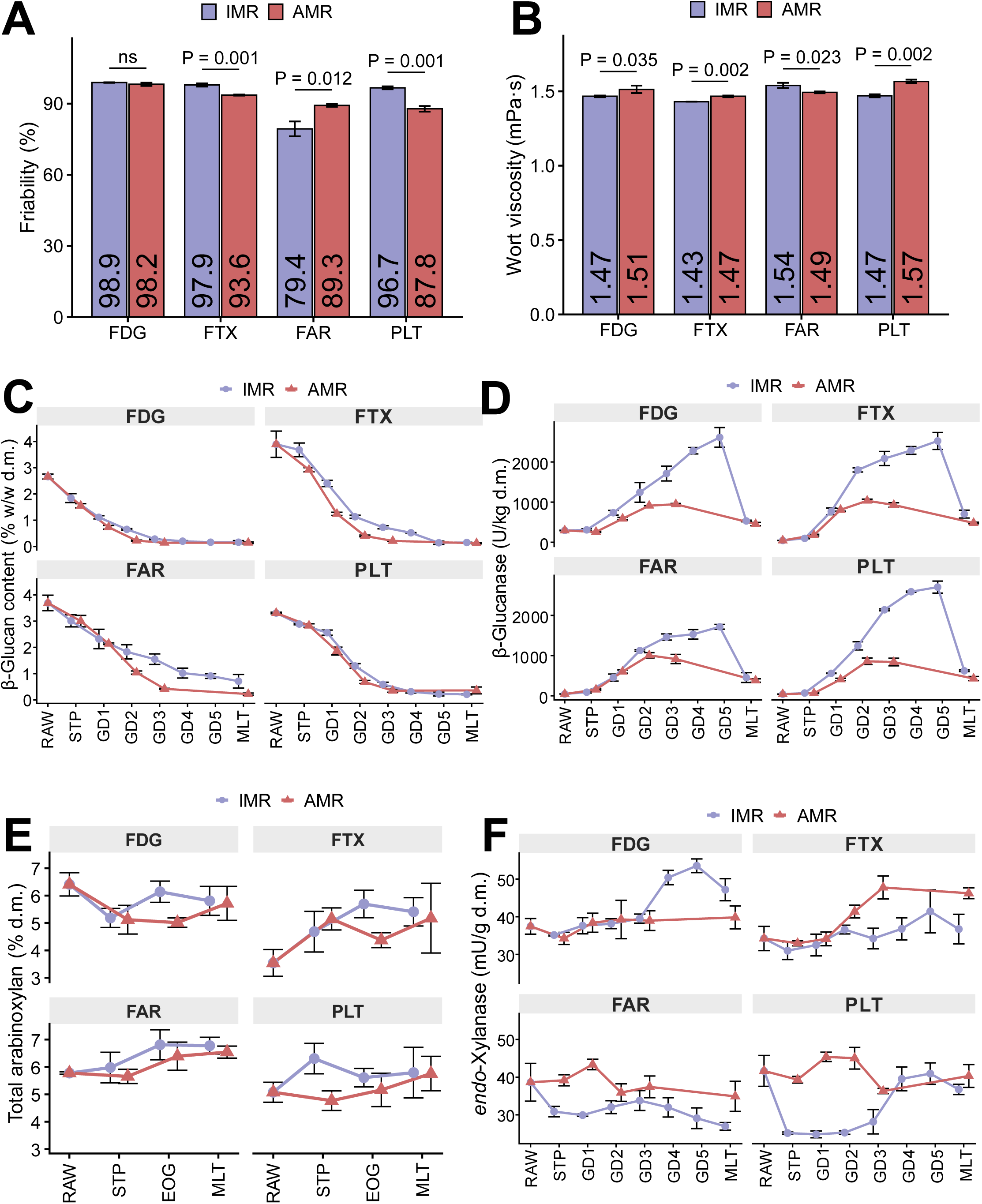
Variety-wise comparison of the cytolytic malt quality attributes, β-glucanase activity and endo-1,4-β-xylanase activity of AMR and IMR malts. **(A–E)** Malt quality attributes used to assess cell wall hydrolysis: **(A)** Friability (n = 3), **(B)** Wort viscosity (n = 3), **(C)** (1→3; 1→4)-β-glucan content (n = 3), and (E) total arabinoxylan content (n = 3). **(D, F)** The development pattern of β-glucanase activity **(D)** and endo-1,4-β-xylanase activity **(F)** in each variety × time point × regime combination (For both D and F, n = 3). The bar plots and line graphs show means, and error bars denote the standard deviation. *P*-values (IMR vs. AMR) were calculated using a Student’s t-test with Bonferroni-Holm adjustment to account for multiple testing and are given in the respective plots (ns: adjusted *P*-value > 0.05). The meaning of the abbreviations is as follows: FDG, Fandaga; FTX, Fantex; FAR, Faro; PLT, Planet; RAW (Ungerminated barley grain); STP (end of steeping or start of germination); GD (Germination day); MLT (finished malt; malt after kilning with rootlets removed); d.m., measurement on a dry-matter basis; EOG (end of germination), meaning that only the last days of germination (GD3 for AMR and GD5 for IMR) in both regimes were considered at this time point.

**Figure 5.**
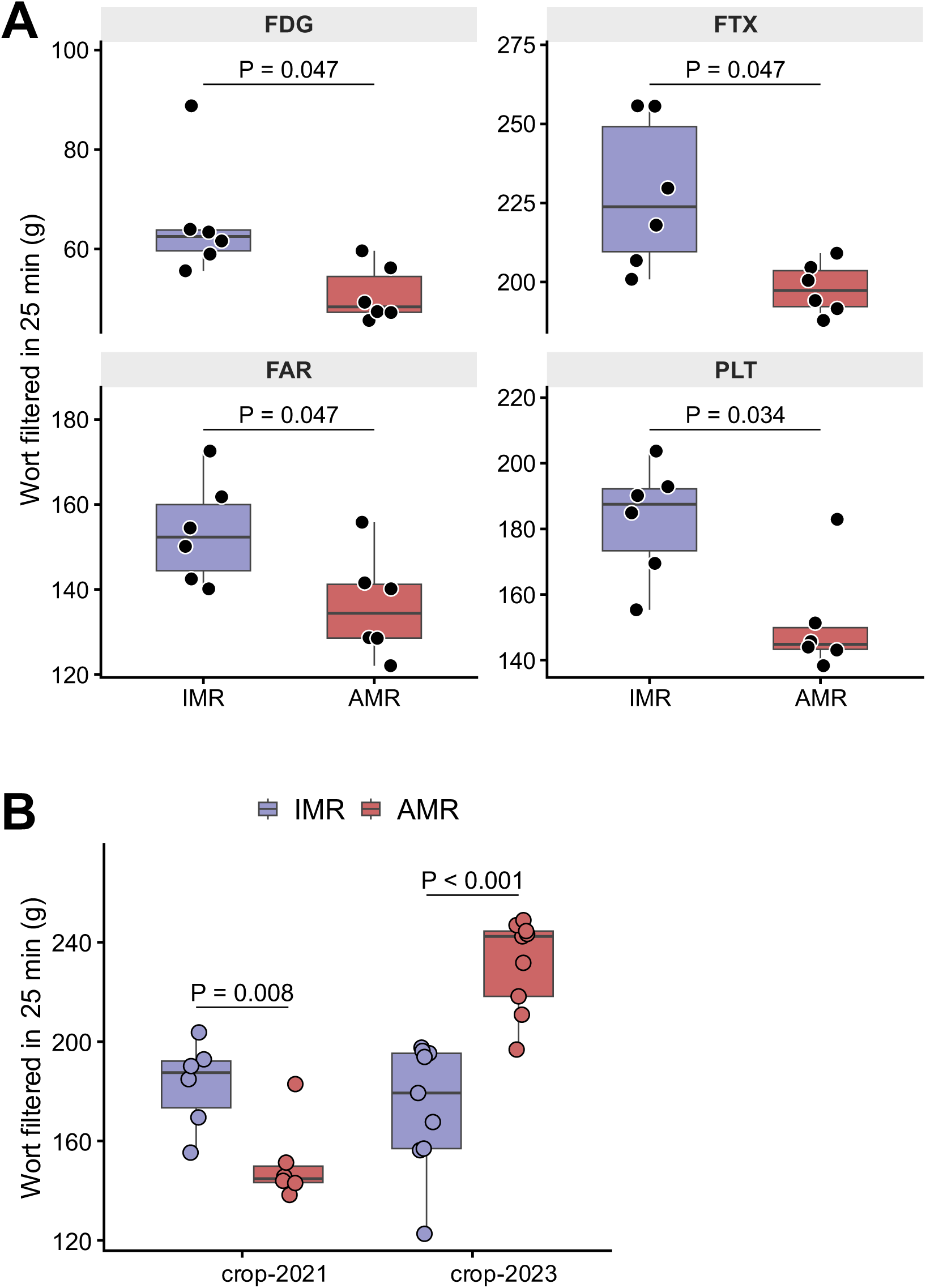
Lautering performance of AMR and IMR malts. (**A**) Comparison of the lautering performance of the AMR and IMR malts per barley variety (n = 6). (**B**) Comparison of the lautering performance of the AMR and IMR malts of the crop year 2021 and crop year 2023 both of the Planet cultivars (2021 crop: n = 6; 2023 crop: n = 9). The boxplots represent the amount of wort yielded after 25 minutes of filtration per group, summarised by the first and third quartiles as hinges of the box, the median as the horizontal line inside the box and the whiskers, which extend to the most extreme values within 1.5 × their interquartile range. Points beyond the whiskers are outliers. *P*-values shown on top of the boxplots indicate significant differences between the tested groups (Student’s t-test with Bonferroni-Holm adjustment to account for multiple testing; Significant difference = adjusted *P*-value ≤ 0.05). FDG, Fandaga; FAR, Faro; FTX, Fantex; PLT, Planet.

### 3.3. Temporal metagenomic and metaproteomic analysis of malt produced via IMR or AMR

#### 3.3.1. Metagenomics-informed composition and diversity of the barley microbiome

For microbiome analysis, barley was sampled at the same time points as for the chemical-technical and enzymatic analyses (**Figure 1A and 1B**), except that samples of the 2023 Planet crop were additionally included for shotgun metagenomics, totalling 173 samples. Metagenomic DNA was extracted from the barley kernel surfaces, meta-barcoded and analysed by Nanopore long-read sequencing. After removal of reads shorter than 500 bp, reads with a Q-score < 9 and human contaminants, a total of 2.7 million reads (5.9 Gbp) with an N50 of 3.67 kbp and a median Q-score of 21.4 were obtained. As it is recommended to have obtained at least 0.5 million reads for taxonomic classification (Hillmann et al., 2018), we performed taxa identification using the non-demultiplexed FASTQ file (2.7 million reads), while for quantification of the microbes in the individual samples the de-multiplexed version was used. Using this workflow, 275 species (246 bacteria, 15 fungi, 14 viruses; **Supplementary Table 3**) in 132 genera (99 bacteria, 20 fungi, 13 viruses; **Supplementary Table 3**) were identified and quantified. In general, each raw barley variety hosted its own microbial community in terms of members and relative abundancies, and this community remained variety-specific at each sampling time point and under both malting regimes (**Figure 6A and B**). Principal coordinate analysis of the raw barley varieties based on Bray-Curtis dissimilarities revealed a clear separation of Faro and Fandaga, while Fantex and Planet overlapped (**Figure 8A**), indicating that the initial microbial composition of Faro and Fandaga was more distinct from that of Fantex and Planet. This pattern was also confirmed by pairwise comparisons; Faro differed most from the other varieties (R^2^ = 0.53–0.71), Fandaga was intermediate (R^2^ = 0.32–0.36 against Fantex and Planet), and Fantex and Planet were the most similar pair (R^2^ = 0.20). PERMANOVA (permutational multivariate analysis of variance) analysis of the varietal effect on the observed variation in Bray-Curtis dissimilarities between the raw barley grain samples was significant (F-statistic = 3.40, *P* = 0.002; **Figure 7A**), while the multivariate dispersion was not (PERMDISP: F-statistic = 1.24, *P* = 0.356; **Supplementary Table 4**), indicating that the PERMANOVA result reflected differences in microbial community composition between the raw barley varieties rather than within-variety variability.

**Figure 6.**
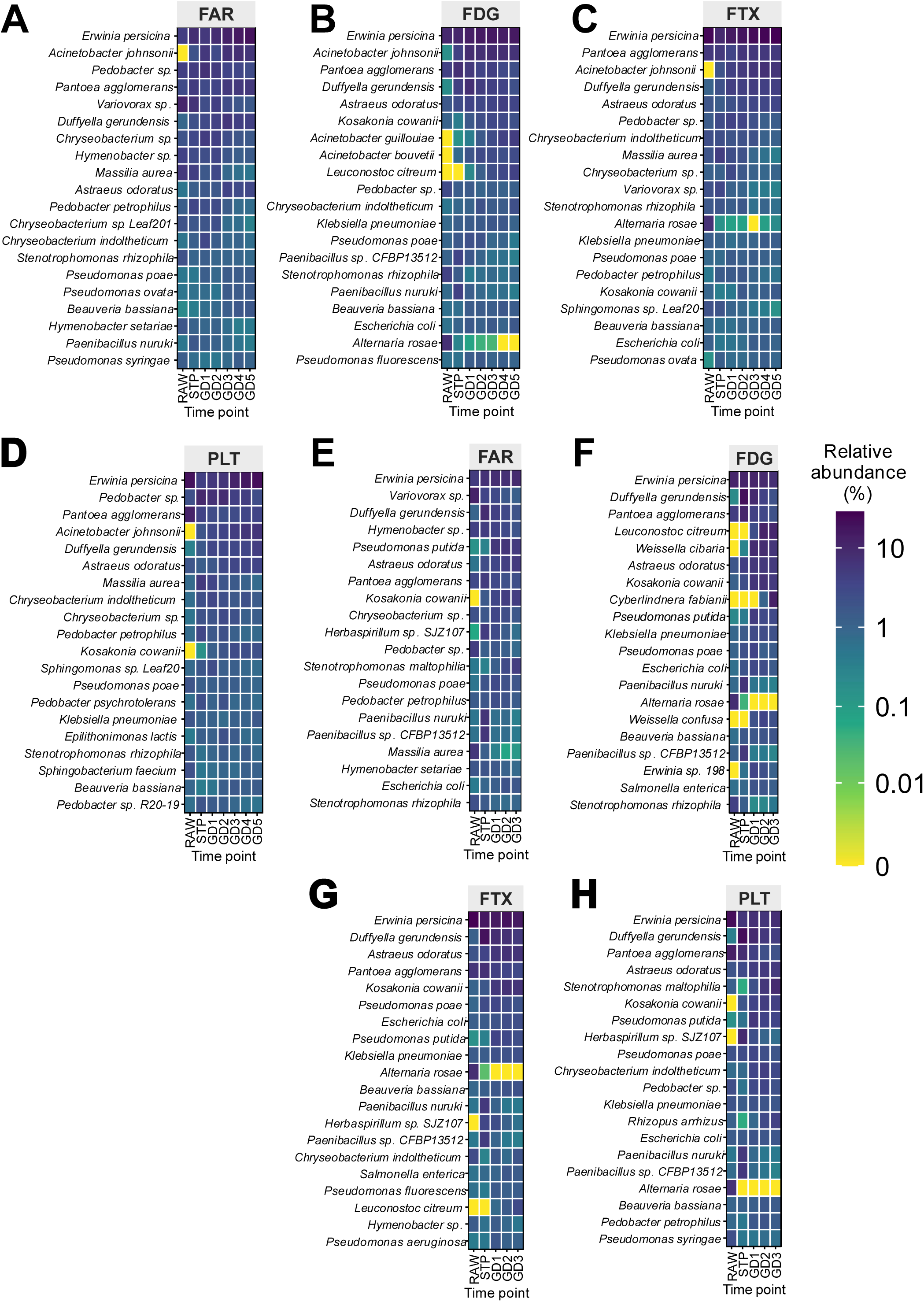
Metagenomics-informed species-level composition of the barley-associated microbiome during malting under the industrial and accelerated malting regimes. The heatmaps show the top 20 most abundant species in each barley variety during malting under the industrial malting regime **(A–D)** and the accelerated malting regime **(E–H)**. The colour indicates the mean relative abundance (% of reads per sample, averaged over n = 3 biological replicates per sampling time point) on a log_10_ scale. The species are arranged from top to bottom by decreasing total abundance. FDG, Fandaga; FTX, Fantex; FAR, Faro; PLT, Planet; RAW (Ungerminated barley grain); STP (end of steeping or start of germination); GD (Germination day).

**Figure 7.**
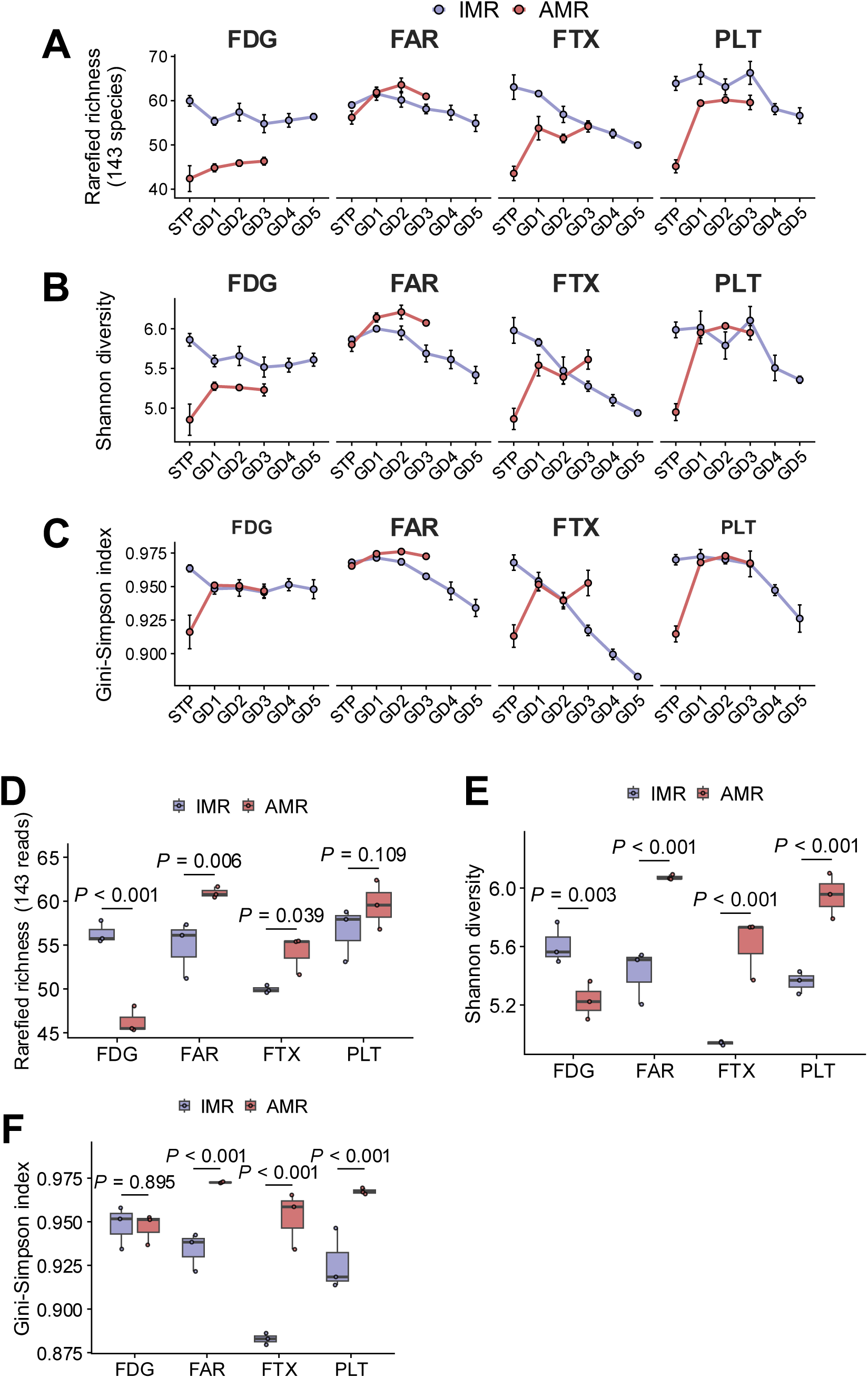
Impact of the accelerated malting regime on the alpha diversity of the barley-associated microbiome during malting. (**A**, **B**, **C**) Temporal changes in (**A**) rarefied species richness (143 reads per sample), (**B**) Shannon diversity, and (**C**) Gini-Simpson index for each barley variety under IMR and AMR. Points show the mean of the corresponding alpha index ± standard error of three biological replicates, with the exception of Fantex IMR at GD1 and Planet IMR at GD3, which had only two replicates. (**D**, **E**, **F**) Comparison of the alpha diversity indices of the IMR and AMR at the end of germination (IMR GD5 vs AMR GD3; n = 3 per malting regime and variety), per variety. *P*-values show comparisons between IMR and AMR within a variety, using estimated marginal means from a two-way ANOVA (regime × variety), corrected using the Benjamini-Hochberg false discovery rate procedure across the four varieties. Boxplots show the median (horizontal line inside the box) and interquartile range. Whiskers denote the lowest and highest values within 1.5 × the interquartile range. Points indicate individual samples tested. STP, end of steeping; GD, germination day; FDG, Fandaga; FAR, Faro; FTX, Fantex; PLT, Planet.

**Figure 8.**
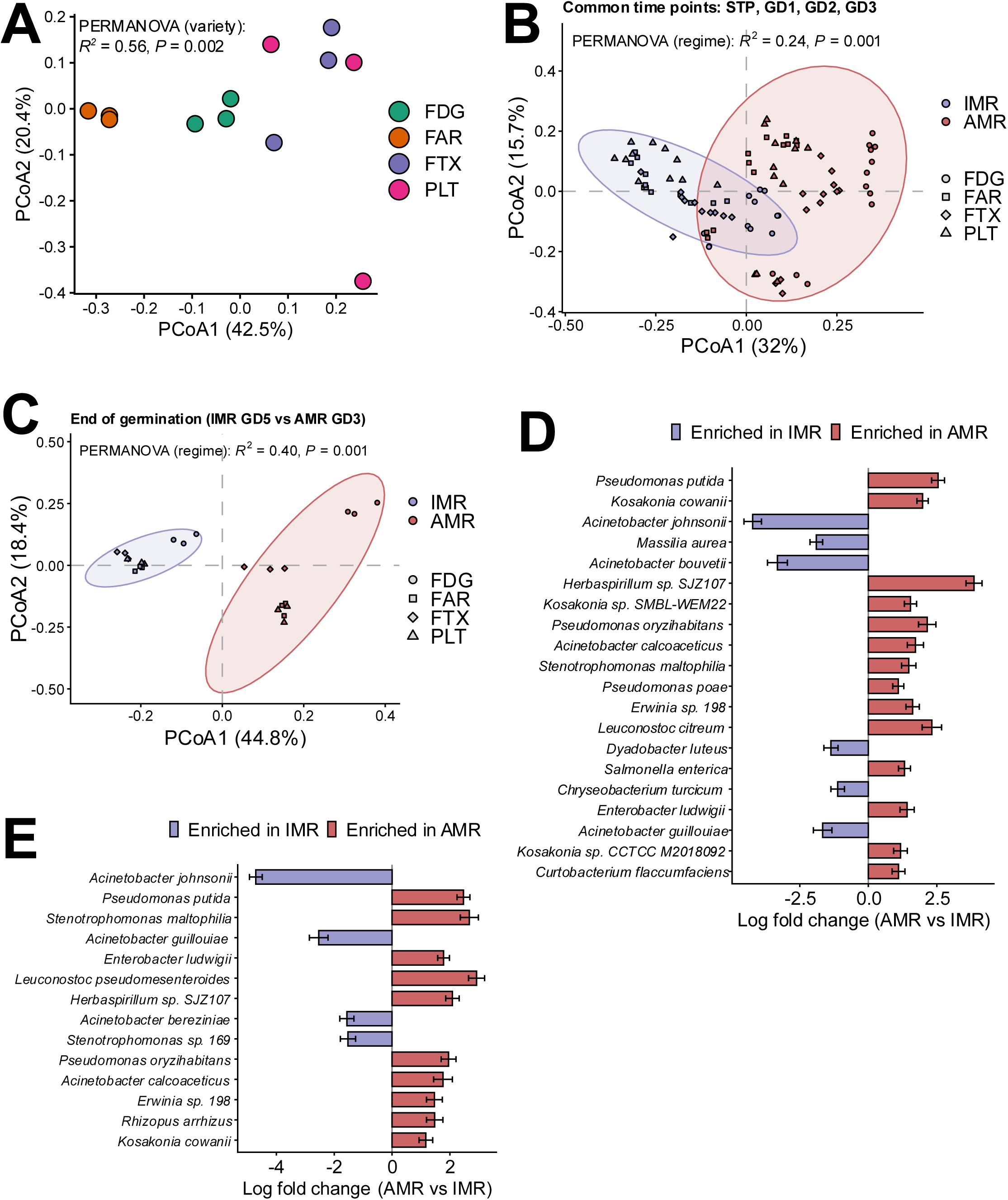
Raw barley varieties hosted distinct microbial communities, and the accelerated malting regime restructured the composition of the barley-associated microbiome during malting. **(A)** Principal coordinate analysis (PCoA) of the Bray-Curtis dissimilarities between raw barley samples of the four varieties before steeping (n = 3 biological replicates per variety). Faro and Fandaga formed clear clusters separated from Fantex and Planet, which overlapped. PERMANOVA analysis of the varietal effect was significant (F-statistic = 3.40, *P* = 0.002), while the multivariate dispersion did not differ between the varieties (PERMDISP: F-statistic = 1.24, *P* = 0.356), indicating that the separation reflected by the PERMANOVA result was due to differences in microbial composition rather than variability in dispersion. **(B)** PCoA of samples from time points during germination shared between IMR and AMR (STP, GD2–GD3, n = 94). Here, we also see a clear separation of the AMR samples from the IMR samples, with a significant PERMANOVA (F-statistic = 91.4, *P* = 0.001) and homogeneous multivariate dispersion (PERMDISP: F-statistic = 0.198, *P* = 0.665), implying that the separation observed between AMR and IMR samples in the PCoA biplot is due to differences in microbial composition between the two regimes. **(C)** PCoA of samples at the end of germination for both regimes (IMR GD5 vs AMR GD3; n = 24). Here, regime explained 40% of the compositional variation (PERMANOVA on variety × regime, R^2^ = 0.40, *P* = 0.001, 999 permutations; PERMDISP *P* = 0.532). Altogether, these results (**B** and **C**) suggest that AMR shifted the composition of the barley-associated microbiome during germination. For the PCoA biplots, ellipses represent a 95% confidence interval for each regime. Axis labels give the percentage of variation explained by each coordinate. (**D, E**) Species robustly differentially abundant between AMR and IMR as identified by ANCOM-BC2 analysis on sample datasets from **(D)** common time points (fixed effects: regime, variety and time point; biological replicate as a random effect) and **(E)** end of germination (fixed effects: regime and variety). The bar plots show the log fold change (AMR vs IMR) ± standard error. The bars are colour-coded to indicate species enriched either in AMR (▪, with positive log fold changes) or in IMR (▪, with negative log fold changes). Only species with a Holm-adjusted *P-*value < 0.05 that also passed the pseudo-count sensitivity analysis are shown, ordered by decreasing –log_10_ adjusted *P-*value. FDG, Fandaga; FTX, Fantex; FAR, Faro; PLT, Planet; STP (end of steeping/start of germination); GD (Germination day).

##### 3.3.1.1. Impact of AMR on alpha diversity of the barley germination microbiome

To assess the impact of the malting regime on the observed species richness, Shannon diversity and Simpson index, we fitted linear mixed-effects models (LMEM) with bucket identity (or the biological replicate) as a random effect, on two sets of data: 1) subsampled from germination time points shared by the two regimes (STP and GD1–GD3), and 2) based on the last germination days of both regimes (IMR, GD5 and AMR, GD3). Our results showed that although AMR during steeping lowered the alpha diversity, this difference largely disappeared during germination (see **Figure 7A–C**). At the end of steeping, mean rarefied richness in AMR was 46.8 species compared with 61.5 in IMR, while by the end of GD3, the difference had reduced to 3.1 species (**Supplementary Table 5**). Both Shannon and Simpson diversity were significantly lower in AMR at STP (Contrast AMR-IMR; Shannon, –0.804, *P*_adj_ = 8.08×10^-15^; Simpson, −0.04, *P*_adj_ = 1.96×10^-16^) but slightly higher than in IMR by GD3 (Shannon,0.07, *P*_adj_ = 0.481; Simpson 0.013, *P*_adj_ = 8.45×10^-4^; **Supplementary Table 5**) However, the steeping effect was variety-specific. In Fandaga, Fantex and Planet, AMR reduced richness by 17.6–19.5 species and Shannon diversity by 1.01–1.11 (all *P* < 0.001), whereas Faro was unaffected (richness, –2.8, *P* = 0.232; Shannon –0.06, *P* = 0.682). Averaged over time, Faro was the only variety whose richness did not differ between regimes (+0.9 species, *P* = 0.474), while Fandaga showed the largest reduction (–12.0 species, *P*_adj_ = 1.14×10^-7^) and remained lower in AMR at every time point (see **Supplementary Table 5)**. When we considered only the last germination days of both regimes, our results showed that AMR green malts had higher Shannon and Simpson diversity than IMR green malts, but similar richness. The mean Shannon diversity we found was 5.72 in AMR and 5.33 in IMR (difference +0.38, *P*_adj_ = 3.06×10^-6^), whereas the Simpson diversity was 0.960 and 0.923 (+0.037, *P*_adj_ = 3.06×10^-7^), respectively. Rarefied richness did not differ overall (55.2 vs 54.4 species, *P*_adj_ = 0.371) because varieties responded in opposite directions (regime × variety, F (3,16) = 17.5, *P*_adj_ = 2.65×10^-5^). For all comparisons at the end of germination, see **Supplementary Table 5**. For comparisons per variety (**Figure 7D**-**F** **and Supplementary Table 5**), Faro (+0.65, *P*_adj_ = 4.13×10^-5^), Fantex (+0.67, *P*_adj_ = 4.13×10^-5^) and Planet (+0.59, *P*_adj_ = 8.32×10^-5^) were more diverse (based on Shannon) at the end of germination in AMR, whereas Fandaga was less rich (–10.04, *P*_adj_ = 1.28×10^-4^) and diverse (–0.38, *P*_adj_ = 3.31×10^-3^) and did not differ in its Simpson diversity (–0.001, *P*_adj_ = 0.895). Overall, in IMR, Shannon diversity declined steadily from GD1 to GD5 (mean 5.86–5.33), whereas in AMR, it stayed relatively constant from GD1 to GD3 (mean 5.72–5.73).

##### 3.3.1.2. Impact of AMR on the barley germination microbial community composition

PERMANOVA analyses based on the Bray-Curtis dissimilarity metric revealed the regime as the strongest single driver of the barley microbial composition during germination. Using the common time points dataset, the malting regime main effect explained 23.9% of the variation in Bray-Curtis dissimilarities (PERMANOVA: F-statistic = 91.4, *P* = 0.001; **Figure 8B**), followed by the variety main effect at 21.9% (F-statistic = 27.9, *P* = 0.001) and time point at 15.6% (F-statistic = 19.9, *P* = 0.001). Regime also interacted with time point (7.9%, *P* = 0.001) and with variety (4.9%, *P* = 0.001), whereas the three-way interaction was not significant (3.8%, *P* = 0.063). All results can be seen in **Supplementary Table 6**. Dispersion did not differ between the two regimes (PERMDISP: F-statistic = 0.20, *P* = 0.665) nor among the eight regime × variety groups (F-statistic = 0.86, *P* = 0.532), indicating that the observed shift in microbial community composition was a result of the malting regime effect itself rather than a result of the variability among samples within a regime. Further principal coordinate analysis at sampling time points showed a clear separation of the regimes at each time point, with regime explaining 34–43% of the total variation (**Supplementary Figure 10**). Interestingly, the separation between the two regimes was largest at the end of steeping (R^2^ = 0.43, n = 24), although its dispersion and that at GD1 also differed significantly between regimes (STP, *P* = 0.021; GD1, *P* = 0.028; **Supplementary Table 6**), indicating that a part of the separation at these two time points also reflected differences in within-regime variability. However, from GD2 onward and at the end of germination, dispersion was homogeneous, indicating that separation reflected differences in microbial community composition. Within each variety, the malting regime explained 35–38% of the compositional variation over the common time points (Fandaga R^2^ = 0.38, Faro 0.35, Fantex 0.37, Planet 0.35; all with *P*_adj_ = 0.001), with the regime × time interaction also significant in each variety (R^2^ = 0.12–0.17, *P* = 0.005–0.025). At the end of germination, regime separated the communities strongly within every variety (Fandaga R^2^ = 0.77, Faro 0.74, Fantex 0.76, and Planet 0.68; **Supplementary Table 6**). However, these within-variety effects are only reported here descriptively, because of the limited number of replicates per regime that were compared, which could only guarantee the smallest attainable permutation *P*-values of 0.10, which all four comparisons reached (see **Supplementary Table 6**). In sum, our results show that the accelerated malting regime restructured the barley-associated microbiome more than barley variety or germination time did.

##### 3.3.1.3. Taxonomic barley germination microbiome differential abundance analysis

To identify the taxa driving microbial compositional differences between AMR and IMR observed by PERMANOVA analysis, we performed differential abundance analysis using ANCOM-BC2 at the species level using two sets of data composed of: 1) common time points between the two regimes (STP, GD1–GD3), and 2) end of germination time points (GD3 for AMR, and GD5 for IMR). Based on the common time points dataset, 30 out of the 261 species tested significantly differed in abundance between AMR and IMR (Holm-adjusted *P* < 0.05; see **Supplementary Table 7** for full list), of which only 20 were robust to the pseudo-count test (14 species enriched in AMR and 6 in IMR; **Supplementary Table 8**). For comparisons performed using the end of germination dataset (AMR-GD3 vs IMR-GD5), 18 out of 254 species were significantly different (**Supplementary Table 7**), of which only 14 passed the pseudo-count sensitivity test (10 enriched in AMR and 4 in IMR; **Supplementary Table 8**). Overall, the two comparisons largely agreed. Ten species were robust in both, and every species robust in one comparison changed in the same direction in the other. In total, 24 species were robustly differentially abundant in at least one comparison (16 enriched in AMR and 8 in IMR; see **Table 1**). Interestingly, AMR generally favoured the proliferation of Gram-negative species of *Pseudomonas*, *Kosaknia*, *Enterobacter, Erwinia, Herbaspirillum and Stenotrophomonas*, together with lactic acid bacteria; *Leuconostoc citreum* and *L. pseudomesenteroides* (**Figure 8D**), whereas IMR favoured several *Acinetobacter* species and the Bacteroidota *Dyadobacter luteus and Chryseobacterium turcicum*, together with *Massilia aurea* (**Figure 8D**). The most striking difference observed between AMR and IMR was the near-disappearance of *Acinetobacter johnsonii* under AMR. It was detected in every IMR sample but in only 13% of AMR samples, and its mean relative abundance fell from 5.4% to 0.006% (log fold change (logFC) –4.22, approx. 68-fold, *P*_adj_ = 1.31×10^-9^; **Supplementary Table 9**). Other notable differences include *Acinetobacter bouvetii*, which was detected in 96% of IMR samples but only in 10% of the AMR samples (logFC –3.32, *P*_adj_ = 4.83×10^-9^); *Herbaspirillum sp. SJZ107* and *Pseudomonas oryzihabitans* that were detected in 17–20% of IMR samples but in nearly all AMR samples (**Supplementary Table 9**). When we tested the species abundance at each time point between AMR and IMR, we observed that at each sampling time point, different species were robustly highly abundant: 13 at STP (11 enriched in AMR), 18 at GD1 (15 enriched in AMR), 13 at GD2 (8 enriched in AMR) and 14 at GD3 (9 enriched in AMR). For the full list, see **Supplementary Table 10**. Out of these species, a few patterns were noticeable: 1) early responders such as *Kosakonia, Herbaspirillum* and *Curtobacterium*, which differed most at STP with declining fold changes during germination; 2) those that were robustly different at every germination day but not at STP such as *P. putida*, *P. oryzihabitans*, *L. citreum*, *E. ludwigii* and *M. aurea*; and 3) late responders such as *S. matophilia* whose enrichment (in logFC) in AMR increased from 0.17 at STP to 2.35 at GD3, and *A. calcoaceticus* from –0.71 at STP to 2.39 at GD3. For the comparisons within a single variety, only a few species were robustly different. Five species were robustly different in Faro and Fandaga and one each in Fantex and Planet: *A. johnsonii* was robustly depleted under AMR in both Faro and Fandaga (Faro: logFC –5.34, *P*_adj_ = 2.66×10^-4^; Fandaga, logFC –4.51, *P*_adj_ = 1.37×10^-3^); *Herbaspirillum sp*. SJZ107 was enriched in Faro (logFC = 3.37, *P*_adj_ = 3.01×10^-4^) and Planet (logFC = 5.12, *P*_adj_ = 0.011), and *Kosakonia cowanii,* was enriched in Faro (logFC = 1.75, *P*_adj_ = 0.011) and Fantex (logFC = 1.90, *P*_adj_ = 1.38×10^-3^). Species that were enriched in specific varieties were: *Pseudomonas putida* (logFC = 2.46, *P*_adj_ = 1.96×10^-3^) and *Paenarthrobacter ilicis* (logFC = 3.24, *P*_adj_ = 0.043) in Faro and *Pseudomonas oryzihabitans* (logFC = 2.76, *P*_adj_ = 0.001), *Curtobacterium flaccumfaciens* (logFC = 1.72, *P*_adj_ = 5.44×10^-4^), *Acinetobacter bouvetii* (logFC = –4.79, *P*_adj_ = 6.58×10^-4^) and *Acinetobacter guillouiae* (logFC = –2.38, *P*_adj_ = 0.024) in Fandaga (see **Supplementary Table 10** for the full list of all robust species identified by ANCOM-BC2 for each variety). Altogether, these data show that AMR selected for fast-growing Gram-negative bacteria. Most of the species enriched under AMR belonged mainly to the Enterobacterales (*Kosakonia*, *Enterobacter*, *Erwinia*), *Pseudomonas, Stenotrophomonas* and *Herbaspirillum,* which could be because of the higher steeping and germination temperature of AMR (20 °C, vs 12–16 °C in IMR) and its long 18-hour dry rest that likely favoured the proliferation of these rapidly growing mesophilic taxa. From a malting standpoint, the rapid growth of these Gram-negative bacteria has been linked to poor lautering performance problems in malt (Laitila et al., 2007; Raulio et al., 2009; Tsitko et al., 2018), which we speculate could be the reason for the higher wort filtration times observed in AMR compared to IMR malts.

**Table 1.** Metagenomics-detected species that were enriched in AMR and IMR. The results are presented as the natural log fold change in AMR compared to IMR of observed abundance by ANCOM-BC2 of a species for the common time points and end-of-germination datasets. Positive values indicate that the species is higher abundant in AMR. Bolded Holm-adjusted *P*-values < 0.05 indicate species that passed the pseudo-count sensitivity analysis. Linear mixed-effects models used: for the common time points dataset: regime + variety + time point as fixed effects and the biological replicate (or Bucket identity) as a random effect; end of germination: regime + variety.

| **Species** | **Common time points** |  | **End of germination** |  |
|---|---|---|---|---|
|  | **Log-fold change** | **Adjusted *P*-value** | **Log-fold change** | **Adjusted *P*-value** |
| **Enriched in AMR** |  |  |  |  |
| *Herbaspirillum sp. SJZ107* | **3.86** | **8.78E-08** | **2.09** | **4.73E-03** |
| *Leuconostoc pseudomesenteroides* | 3.29 | 1.02E-01 | **2.92** | **5.68E-04** |
| *Pseudomonas putida* | **2.55** | **8.35E-15** | **2.47** | **4.92E-07** |
| *Leuconostoc citreum* | **2.32** | **2.05E-03** | 1.21 | 1.95E-01 |
| *Pseudomonas oryzihabitans* | **2.15** | **4.03E-06** | **1.95** | **8.08E-03** |
| *Kosakonia cowanii* | **1.98** | **1.38E-12** | **1.17** | **2.08E-02** |
| *Acinetobacter calcoaceticus* | **1.72** | **5.72E-05** | **1.76** | **8.96E-03** |
| *Erwinia sp. 198* | **1.62** | **8.52E-04** | **1.46** | **1.26E-02** |
| *Kosakonia sp. SMBL-WEM22* | **1.54** | **3.99E-07** | 1.12 | 1.10E-01 |
| *Stenotrophomonas maltophilia* | **1.48** | **6.82E-05** | **2.67** | **3.71E-05** |
| *Rhizopus arrhizus* | 0.90 | 1.69E-01 | **1.47** | **1.43E-02** |
| *Enterobacter ludwigii* | **1.42** | **1.01E-02** | **1.78** | **3.94E-04** |
| *Salmonella enterica* | **1.32** | **4.95E-03** | 0.83 | 1.00E+00 |
| *Kosakonia sp. CCTCC M2018092* | **1.18** | **3.27E-02** | 1.26 | 8.84E-02 |
| *Curtobacterium flaccumfaciens* | **1.10** | **3.56E-02** | 0.64 | 1.00E+00 |
| *Pseudomonas poae* | **1.09** | **1.44E-04** | 0.87 | 7.87E-01 |
| **Enriched in IMR** |  |  |  |  |
| *Acinetobacter johnsonii* | **–4.22** | **1.31E-09** | **–4.71** | **6.70E-08** |
| *Acinetobacter bouvetii* | **–3.32** | **4.83E-09** | Not tested | 1.00E+00 |
| *Massilia aurea* | **–1.90** | **1.42E-09** | –1.60 | 3.67E-02 |
| *Acinetobacter guillouiae* | **–1.67** | **2.66E-02** | **–2.54** | **7.52E-05** |
| *Acinetobacter bereziniae* | –0.73 | 1.00E+00 | **–1.56** | **5.78E-03** |
| *Stenotrophomonas sp. 169* | –1.15 | 1.76E-03 | **–1.52** | **6.73E-03** |
| *Dyadobacter luteus* | **–1.36** | **4.73E-03** | –1.74 | 1.28E-01 |
| *Chryseobacterium turcicum* | **–1.12** | **6.66E-03** | –0.74 | 1.00E+00 |

#### 3.3.2. Metaproteomic profiling of the barley-associated microbiome

Whereas metagenomics gives an indication of which microbes are present in a sample, it does not reveal which genes (and consequently which proteins) are being expressed. We therefore additionally performed a metaproteomics analysis to obtain functional information on the barley-associated microbiome. Using samples from the same time points as for metagenomics, microorganisms were extracted from the kernel surfaces, proteins were extracted, digested with trypsin, the resulting peptides were TMT-labelled and analysed in 18 individual LC-MS/MS runs. The search space (FASTA) for peptide identification was derived from the species list obtained by the shotgun metagenomics described in 3.3.1. Two experimental design (ED) strategies were applied for metaproteomics analysis: in ED1, all sampling time points of an individual barley variety were multiplexed (changes within a variety over time), whereas in ED2, the barley varieties were multiplexed for each individual sampling time point (changes between barley varieties at defined sampling timepoints). This design strategy was chosen because data-dependent MS analysis selects peptides for identification and quantification semi-stochastically, so that the same set of peptides cannot be guaranteed between consecutive MS analyses, leading to missing data points (Tabb et al., 2010; van Ooijen et al., 2018; Zhang et al., 2016). The usage of TMT compensated for this limitation as all relevant samples are analysed simultaneously within a TMT-multiplex. In case of IMR, 22,199 and 20,417 peptides were identified using ED1 and ED2, respectively, while for AMR, 27,108 and 24,958 peptides were identified in ED1 and ED2 (non-redundant peptides, 5% FDR). Microbial peptide origin was assigned by a lowest common ancestor (LCA) approach using Unipept (Vande Moortele et al., 2025). The majority of peptides (approximately 70% for IMR and 52% for AMR) were assigned to the kingdom Viridiplantae, i.e., to the host *Hordeum vulgare*, and removed from further analysis. In IMR, 13% and in AMR, 28% of the peptide sequences were classified as of microbial; of these, 13.5% were assigned to the taxonomic root (i.e., occurring in both prokaryotes and eukaryotes), and approximately 3.3% of the peptides could not be assigned to any taxonomic level probably because either the submitted amino acid sequence exceeded the length threshold of Unipept (5–50 amino acids long) or because the amino acid came from an organism whose reference sequence did not exist in the reference database. Taxonomically, the barley metaproteome spanned seven phyla (Ascomycota, Pseudomonadota, Bacillota, Bacteroidota, Mucoromycota, Basidiomycota and Actinomycetota), 15 Classes, 24 Orders, 37 Families and 62 genera (**Supplementary Table 11**). The number of species detected varied from 118 in IMR-ED1 to 128 in AMR-ED2. Interestingly, AMR had a higher number of identified LCAs shared among all four barley varieties (see **Figure 9A and B**). This suggests that AMR conditions shifted the active microbial taxa to more uniform communities across the varieties during germination, which also agrees with the trend of the Shannon diversity observed in the metagenomic data (**Supplementary Table 5**).

**Figure 9.**
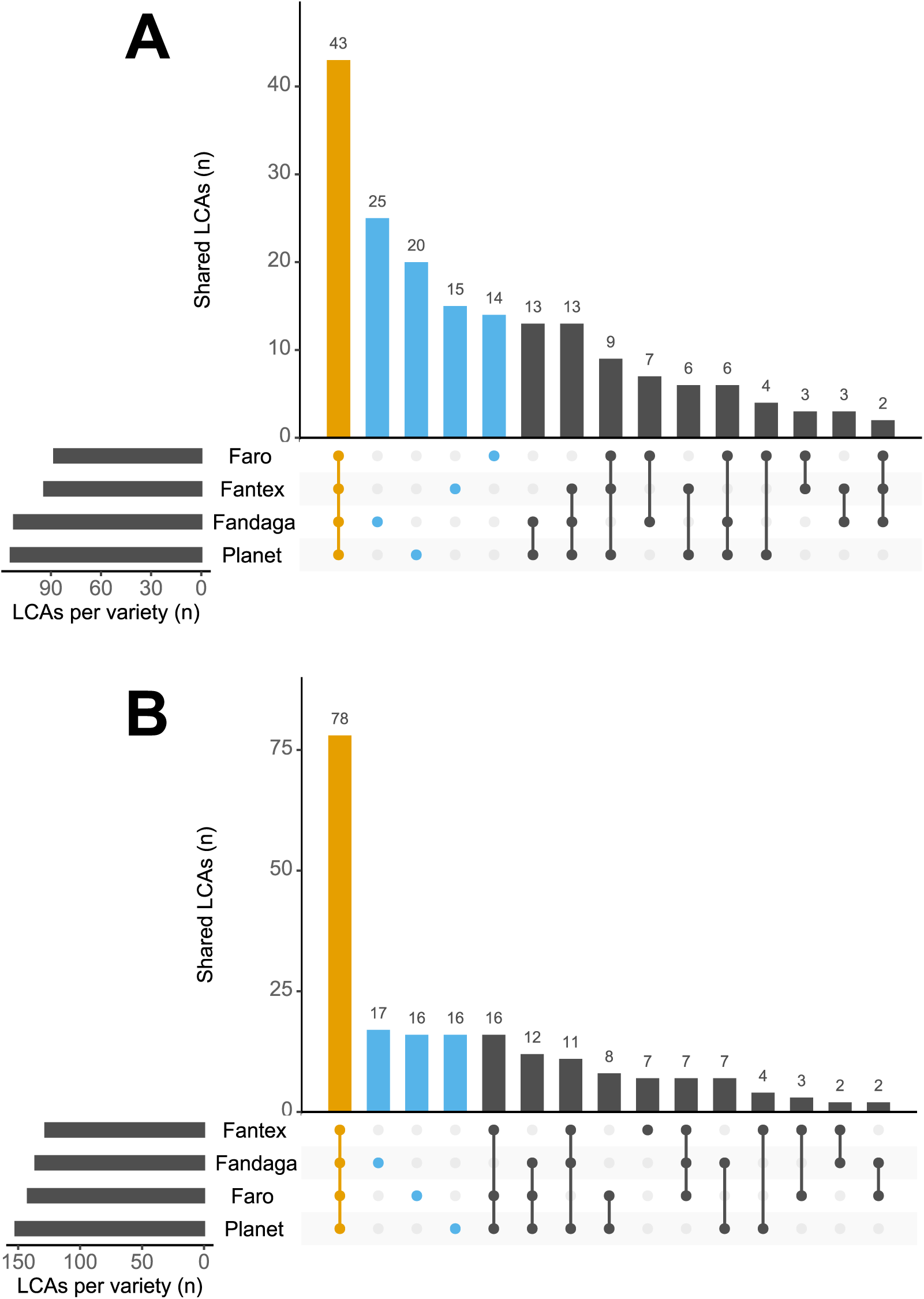
Shared and variety-specific microbial taxa detected by metaproteomics in the four barley varieties during malting. (**A**, **B**) Upset plots of the lowest common ancestors (LCAs) assigned to the peptides identified in the four barley varieties (Fandaga, Fantex, Faro and Planet). (**A**) displays the results of IMR (from STP to GD5), whereas (**B**) displays the results of AMR (from STP to GD3). Peptides were taxonomically assigned using Unipept; peptides without an LCA and those not of bacterial or fungal origin were excluded. The four barley varieties are indicated on the horizontal bars and sorted in increasing order of total number of LCAs identified per variety over the course of the germination phase of malting. The vertical bars show the number of LCAs in each combination of varieties marked by the connected dots below. Dots are colour-coded to show the intersection for all four varieties (orange), and the unique LCAs identified for each variety (blue). STP (end of steeping or start of germination); GD (Germination day).

Before further quantitative peptide analysis of the metaproteomics data, we checked whether the data contained unwanted variation from technical factors that could mask the biological signals of interest. We therefore performed a principal component analysis (PCA) on the VSN-normalised TMT peptide intensities using the InDaPCA algorithm (Podani et al., 2021) to account for the sparsity of the metaproteomics data between individual MS analyses. PCA analysis based on comparisons between mass spectrometry runs revealed batch-to-batch effects, as samples clustered according to the pattern, they were multiplexed within a single mass spectrometry experiment (see **Supplementary Figure 11A**). This means that differences observed when comparing samples across mass spectrometry runs are driven more by unwanted technical variation (likely caused by severe undersampling of the peptide space by the mass spectrometer) than by biological variation. Since TMT-labelling provides a route to compare samples within a label multiplex, we then examined the pattern of clusters formed at this level. As shown in **Supplementary Figures 11B** and **11C**, samples within a multiplex cluster by sampling time point and barley variety, the main sources of biological variation in our study. Thus, these results strongly suggest that for quantitative metaproteomics, a multiplexing strategy is mandatory to be able to overcome biases due to the semi-stochastic selection of peptides for identification and quantification by the mass spectrometer, given the very large set of proteins encountered in a microbiome.

Next, we asked whether the measured intensity of the peptides assigned to a microorganism correlates with the number of metagenomic reads. For each microorganism with a species-level assignment shared between the metaproteomics and metagenomics data (n = 142), the TMT reporter intensities of all detected peptides were summed and VSN-transformed, whereas the reads assigned to each species were normalised to the total number of species-assigned reads in the sample. The Spearman correlation between this sum of VSN-transformed peptide intensities of the microorganism and the TSS-normalised (total sum scaling) reads from the metagenomic data was calculated, resolved by malting regime, variety, time-point and replicate. The Spearman correlation coefficient between metaproteomics sum-of-peptides intensity and TSS-transformed number of reads ranged between - 0.1 and 0.33 for IMR (average, 0.136), whereas for the AMR, it ranged between 0.083 and 0.546 (average 0.217) (see **Supplementary Figure 12A–E**). When instead using the number of detected peptides over the complete course of malting within a barley variety instead of the sum of peptide intensities, the Spearman correlation for the IMR was 0.186 (min: 0.105, max: 0.258), while for the AMR, 0.337 (min: 0.303, max: 0.369, see **Supplementary Figure 13**). As such there is a tendency that a higher number of detected peptides correlates to a higher abundance of the respective microorganism (in contrast to sum-of-peptides intensity).

#### 3.3.3. Functional features of the barley-associated microbiome

Our main interest in metaproteomic data, however, was to identify differences in microbial protein abundancies between IMR and AMR (as well as over time) to gain insight into potential functional differences. From a malt production standpoint, degradation of barley cell wall polysaccharides is a major objective, as otherwise lautering and filtration problems arise from the increased viscosity caused by the release of high-molecular-weight β-glucans and arabinoxylans during mashing (Gastl et al., 2020; Sadosky et al., 2002). We therefore performed a Gene Ontology (GO) analysis to associate microbes with the relevant terms. In all four barley varieties, microbial proteins associated with xylan catabolic process (GO:0045493) and beta-glucan catabolic process (GO:0051275) (or their ancestor terms) were detected (see **Supplementary Table 12**).

In the case of IMR, in total 126 peptides (47 unique) mapping onto 38 unique proteins involved in xylan catabolism, and 71 peptides (31 unique) from 27 unique proteins related to beta-glucan catabolism were detected. These peptides were overwhelmingly from *Alternaria alternata* (36 unique peptides), the family Pleosporaceae (15 unique peptides), or the class Dothideomycetes (7 unique peptides), of which *Alternaria alternata* is also part. Other contributing microorganisms were *Pyrenophora teres* (8 unique peptides, which is also from the family Pleosporaceae)*, Paenibacillus nuruk*i (7 unique peptides), 2 unique peptides of a fungus from the genus *Aureobasidium* and single peptides from the domain Bacteria, the kingdom Pseudomonadati and the genus *Pantoea* (see **Supplementary Table 12**). In AMR, in total, 127 peptides (42 unique) from 53 proteins (some peptides map onto more than a single protein) relating to xylan catabolism, and 65 peptides (25 unique) from 22 proteins involved in beta-glucan catabolism were detected. Here as well, the majority of peptides were mapped onto *Alternaria alternata* (30 unique peptides), the family Pleosporaceae (8 unique peptides), or the class Dothideomycetes (7 unique peptides). *Paenibacillus nuruki* (8 unique peptides) and *Pyrenophora teres* (7 unique peptides), and the genus *Aureobasidium* (1 unique peptide) were also detected. However, in contrast to IMR, additionally, single unique peptides from *Acinetobacter*, *Beauveria bassiana*, and *Pedobacter psychrotolerans* were identified (see **Supplementary Table 12**).

We next asked if the peptides associated to these cytolytic proteins were differentially abundant over the germination days and if the malting regime had an influence. Generally, cytolytic proteins were lower abundant at GD1 as in contrast to STP in both IMR and AMR, but strikingly, at the end of germination in AMR the majority of cytolytic proteins were lower abundant than at the STP timepoint, which was not the case in IMR (see **Figure 10A–D** for peptides which show a statistically significant differential abundance between timepoints, for statistical analysis see **Supplementary Table 13** and volcano plots are shown in **Supplementary Figure 14A–D**). We speculate that the increased germination temperature in AMR compensated for the lower protein abundance, resulting in a cytolytic modification similar to IMR malts. Interestingly, analysis of GO terms showed enrichment of xylan catabolism and endo-1,4-beta-xylanase activity in Planet at GD4 in comparison with GD3 in IMR (q = 0.025 for both) which was also associated with an increase in endo-xylanase enzymatic activity (p = 0.012, see **Figure 4F**). Taken together, these results suggest that microbial enzymes contribute to the endo-xylanase activity of barley malt, likely with strong contribution of *Alternaria alternata*.

**Figure 10.**
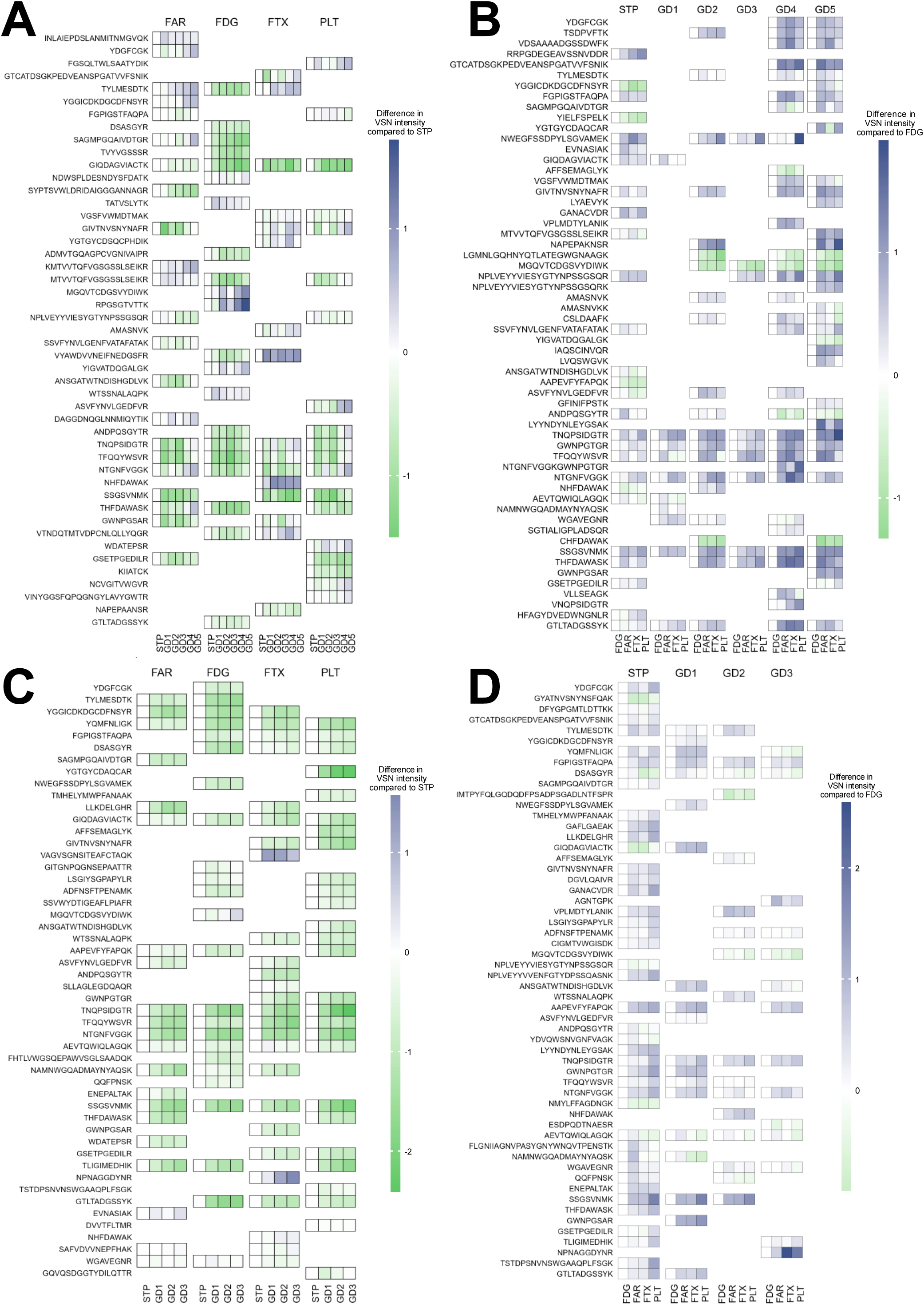
Metaproteomic analysis of peptides associated with GO terms relating to either beta-glucan or xylan catabolism during IMR and AMR. Heatmap showing the relative abundance of VSN-transformed peptide TMT intensities within the respective TMT multiplex (columns): **(A)** Intensities normalised to the STP timepoint during IMR using ED1 (multiplexing of timepoints of a specific barley variety), **(B)** Intensities normalised to the FDG variety during IMR using ED2 (multiplexing of all varieties at specific timepoints), **(C)** AMR using ED1, normalised to STP, and **(D)** AMR using ED2, normalised to FDG. Only peptides which exhibited a statistically significant difference within the multiplex (timepoints or variety) are shown. Positive differences (blue) indicate higher abundance of the respective peptide in comparison with STP or FDG, respectively. Statistics (adj. *P*-values, logFC, Uniprot. Acc. No., protein names, LCA, molecular function and biological processes are given in **Supplementary Table 9**. FDG, Fandaga; FTX, Fantex; FAR, Faro; PLT, Planet; STP (end of steeping or start of germination); GD (Germination day).

The data also suggest that β-glucanase enzymes are mostly derived from *Alternaria alternata* in IMR and AMR (10 and 11 unique peptide, respectively) with additional minor contributions from *Pyrenophora teres* and *Paenibacillus nuruki* (**Supplementary Table 12**). Previous research showed that thermostable β-glucan solubilase activity originates from the barley husk (Yin and MacGregor, 1988) and has been suggested to stem from fungi (Yin et al., 1989). It is therefore tempting to speculate that *Alternaria alternata* contributes these enzymes, however, further experiments are required to determine which of these enzymes, if any, correspond to thermostable enzyme(s) described by Yin and MacGregor (Yin and MacGregor, 1988), particularly as wort composition during mashing also influences the protein thermostability (De Schepper et al., 2021) and thus predicted computational melting temperatures from the protein amino acid sequence might be misleading and were thus not attempted here.

#### 3.3.4. *Lautering-defective malt shows a higher abundance of peptides originating* from a biofilm-forming bacterial genus than in the non-defective batch

Next, we used the quantitative and functional microbial information from the metaproteomics analysis to compare the 2023 Planet crop, which showed no lautering defect, with the lautering-defective 2021 crop to test whether their microbial community compositions and functions differ. First, we examined whether there were any differences in the microbial functional features of these two crop years. The 2021 crop showed a significant enrichment of xylanase catabolic process (GO:0045493) and endo-1,4--xylanase activity (GO:0031176) in peptides more abundant at STP than at GD1 and GD3 (see **Supplementary Table 12**), with most of the peptides attributed to *Alternaria alternata*. Hence, the lautering defect of the 2021 Planet crop is likely not caused by a deficiency in microbes contributing to cell wall degradation.

We were further interested in which microorganisms the differentially abundant peptides belonged to and whether these changed over time, particularly as we showed above that the number of identified peptides correlates with microbial abundance. The 2023 crop showed strong increases in the number of detected peptides at GD3 for *Acinetobacter*, Erwiniaceae, *Pantoea*, Phaffomycetales and *Wickerhamomyces anomalus*, whereas the lautering-defective 2021 crop showed more abundant peptides from *Duffyella gerundensis*, Microbacteriaceae, Micrococcales, *Paenibacillus nuruki* and *Pseudomonas* (**Supplementary Table 14**). At this time point, the lautering-normal 2023 crop had 78 peptides with higher abundance belonging to the genus *Pseudomonas* (out of 956 more abundant peptides in total), while the lautering-defective 2021 crop had 274 peptides originating from the genus *Pseudomonas* (out of 952 more abundant total peptides) (**Supplementary Figure 15, Supplementary Table 14**). This is a significantly higher proportion (exact binomial test: *P* < 2.6 x 10^-5^) and it is therefore tempting to speculate that the predominant detection of *Pseudomonas*, a genus of known strong biofilm formers (Fazli et al., 2014; Mann and Wozniak, 2012; Masák et al., 2014; Siddiqui and Khan, 2017), at the end of germination in the 2021 crop contributes to the observed filtration problems. However, this requires validation in future experiments, ideally accounting for microbial community dynamics, especially as the number of higher abundant peptides belonging to the genus *Pseudomonas* was initially higher at the STP time point in the 2023 crop (n = 205) than in the 2021 crop (n = 42).

In summary, the metaproteomics data suggest that the individual barley microflora can positively influence brewing-relevant malt quality through xylan and glucan catabolism but can also negatively affect malt quality by contributing to filtration problems via biofilm forming microbes.

### 3.4. AMR reduces the overall energy demand and carbon footprint of the malting process

As our results and previous studies indicate that AMR produces malt of similar quality to IMR within a shorter time, we assessed AMR’s energy and CO_2_ savings potential compared to IMR. The energy expenditure for the two regimes was assessed based on the design of the pilot-scale malting facility with a maximum production capacity of 500 kg of malt (starting raw barley capacity between 600– 650 kg) and the regime parameters described in Section 2.1. The facility is supplied with electricity from the national grid and with natural gas, which is used solely to produce steam, mainly for drying. The electrical energy consumption E_p_ (Wh) of the water and air motors, the electrical heaters, and the rotation of the germination arm and spindle was calculated as *E_p_* = *Pt*, where t is the total germination time of each regime (h) and P (kW) is the apparent power of the device. The energy required for air humidification and cooling during the germination phase was determined using the Mollier diagram to characterise the states of both the incoming air and the air entering the germination vessel. For simplicity, we assumed that 1) the incoming air had a dry-bulb temperature of 20 °C and a relative humidity of 60%; 2) the humidification system was 100% efficient, so that the humidified air was fully saturated; and 3) the incoming air had the same state throughout germination, thereby disregarding fluctuations in air temperature and relative humidity as well as recirculation of exhaust air from the germination vessels to conserve energy. The energy (Wh) for air humidification and cooling was calculated using the following equations.

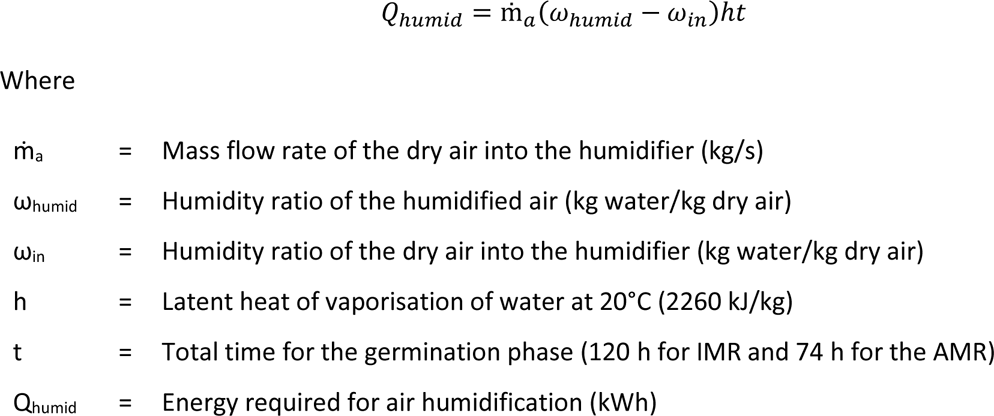

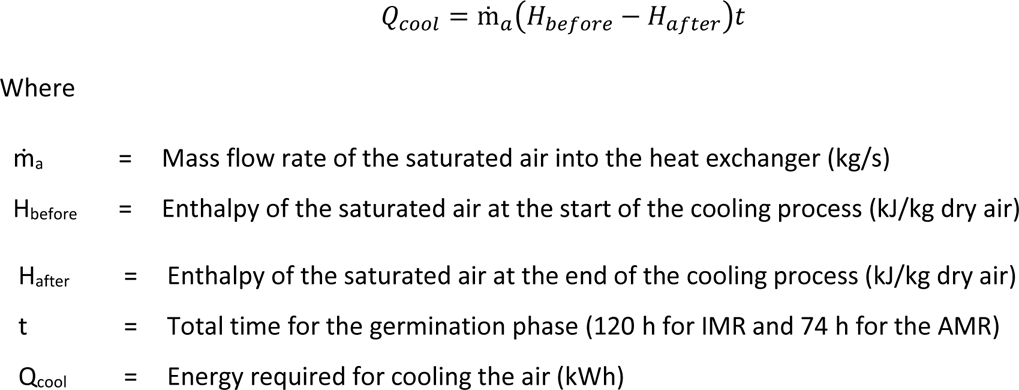

The final temperature of the cooled air (i.e., the temperature of the air entering the grain bed) was assumed to be 2 °C below the germination set-point temperature. The energy needed to evaporate water from the green malt during kilning was estimated as follows. First, we determined the batch size of green malt obtained from 600 kg of barley at 12% moisture content, with final moisture contents of 44% and 47% for IMR and AMR green malt, respectively (see **Supplementary Table 15**). Nevertheless, we also calculated the energy expenditure for equal final moisture contents (44%) of the green malts from both regimes (see **Supplementary Table 15**). Second, we calculated the amount of water to be evaporated during withering, drying and kilning (see **Figure 1A and 1B**), assuming water contents of 6% at the end of withering and drying (30 h) and 4% at the end of kilning (5 h), respectively. Finally, the theoretical thermal energy required to evaporate this water was calculated using the enthalpy of vaporisation of water at 25 °C (0.707 kWh/kg water). However, the actual energy requirement may be higher than calculated, as the drying process does not proceed ideally in its final stages owing to the hygroscopic nature of barley malt (Johnston, 1954). Based on our calculations, AMR uses approximately 5–9% less electrical energy (depending on the final moisture content of the green malt, see **Table 2**) than the reference IMR to produce 500 kg of malt, owing to the lower energy demand for humidification and for cooling the humidified air. At the average EU electricity price for non-household consumers of €0.1902 per kWh (Eurostat, 2025), this corresponds to savings of about €8–15 per 500 kg of malt and to a reduction in CO_2_ emissions of about 5.7–10.6 kg per kWh per 500 kg of malt, based on the official 2024 ENGIE emission factor (131.73 g/kWh, https://business.engie.be/en/faq/contract/co2-emissions).

**Table 2.**
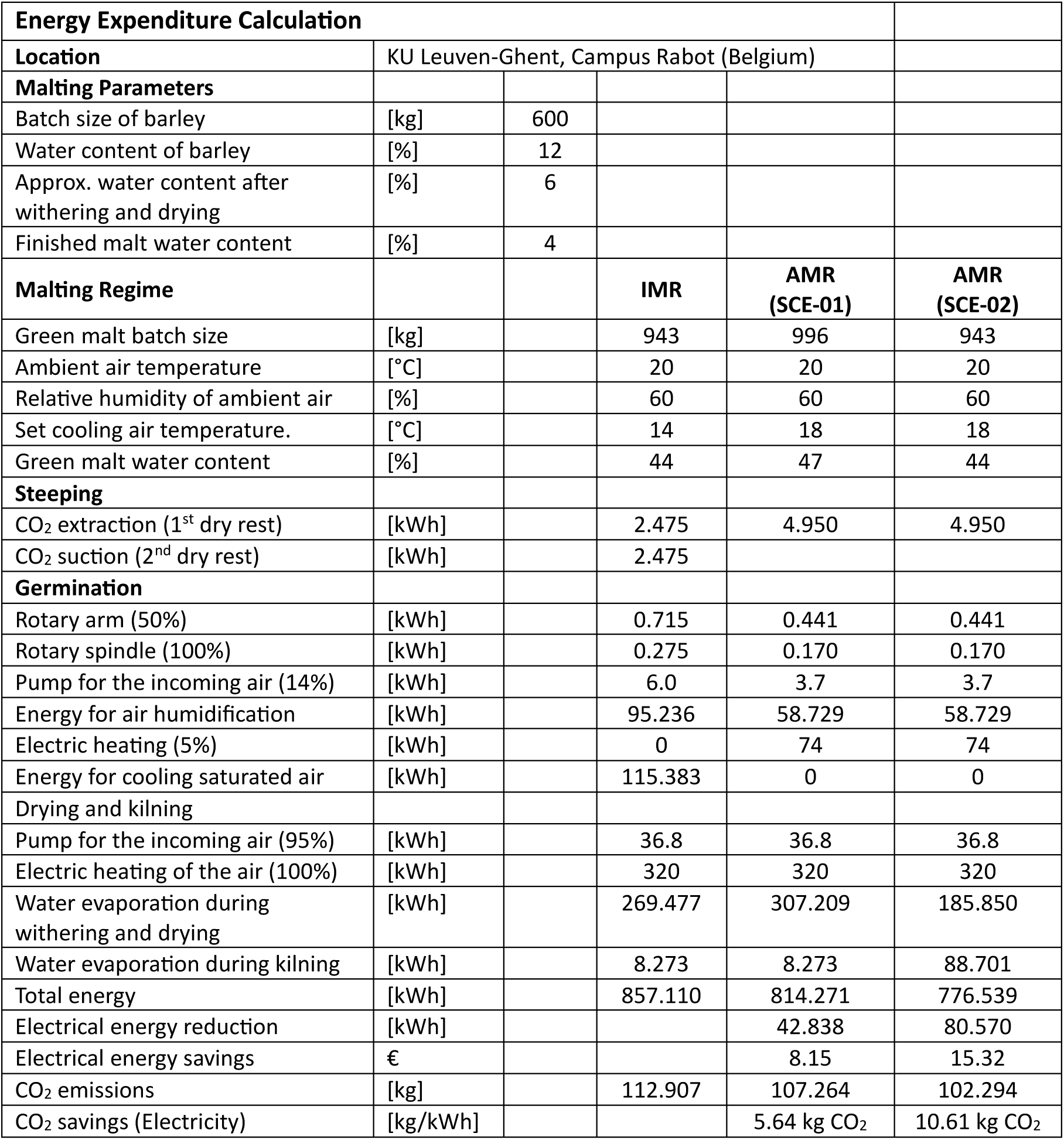
Simplified calculation spreadsheet of the energy expenditure when using the standard industrial malting regime and an accelerated malting regime with the 600–650 kg barley batch capacity pilot-scale malting plant at KU Leuven-Ghent, Campus Rabot. *SCE-01 and SCE-02 indicate scenarios 1 and 2, where the final moisture content of the AMR-green malt is greater than that of the IMR (sce-01) and where they are equal (sce-02)*.

## 4. Discussion

Taken together, our study results show that AMR with a germination phase shortened by 2 days was able to produce high-quality malts similar to a standard malting regime. All AMR malts had acceptable cytolytic modification; they exceeded the commonly accepted friability threshold of 85% and had acceptable wort viscosities and β-glucan degradation. Although β-glucanase development was slower during AMR, the enzymes produced were sufficient, as most varieties reached a similar final percentage of β-glucan degradation to that in IMR. The lower α-amylase, β-amylase and β-glucanase activities and the lower diastatic power of AMR malts were consistent with the plateau in enzyme formation reported for elevated germination temperatures, yet they did not translate into longer saccharification times during wort production. Therefore, within the scope of the analyses performed in the study, the shorter germination time did not compromise the enzymatic capacity required for brewing.

In contrast to earlier work, AMR did not result in a suppression of proteolysis. One possible contributing factor might be the higher moisture content of AMR green malts (around 47% vs 44% in IMR), which can lead to enhanced proteolytic modification (Narziß, 1966). Because most Kolbach indices were at or above the upper limit of the usual specification range, particularly for the Fandaga cultivar, the 3-day germination period of AMR could be shortened further to increase time savings and energy savings. However, such optimal germination conditions seem to be variety-dependent, and future research would need to be conducted to also test for the influence of provenance and crop year.

The lautering performance results illustrate that malt quality is not determined by the malting regime alone. The pronounced filtration deficit of the 2021 AMR malts was not reproduced with the 2023 Planet crop for which AMR even improved the wort filtration volumes. This indicates that the raw material and its associated microbiome interact with the malting regime and influence the resulting malt quality. This interpretation is supported by the metagenomic data which showed that each barley variety carried its own microbial community and that the malting regime influences the composition over time. Further, we show that fungi, in particular *Alternaria alternata* and related Pleosporaceae, contributed xylan catabolic enzymes, indicating a beneficial role in barley cell wall modification, whereas the predominance of the genus Pseudomonas possibly negatively contributes to wort lautering performance. The barley-associated microbiome can therefore act both as a contributor to and a risk to malt quality and would need to be controlled especially when germinating at elevated temperatures.

Several methodological limitations should also be considered. The malting trials were performed at pilot scale with barley mainly from a single crop year, and the crop year comparison was limited to one variety. The proposed link between Pseudomonas and impaired lautering performance is correlative and requires further experimental validation. Finally, the energy assessment relies on simplifying assumptions (constant ambient air conditions, no air recirculation and theoretical evaporation enthalpies), so the absolute savings of 5–9% in electrical energy should be regarded as estimates for the pilot plant rather than values directly transferable to industrial malthouses. In addition, our calculations were based on warm weather conditions during summer and thus should be re-evaluated based on different weather conditions, geographical locations and seasons as these all affect the energy requirements during steeping and germination to achieve the target temperature set points.

In conclusion, AMR is a promising approach to achieve significant time and energy savings in malting, but the results of our metagenomics and -proteomics studies make it clear that the microbiome needs to be controlled to avoid negative influence on the resulting malt quality.

## Acknowledgements

The authors thank the following Master’s thesis students: Mr David De Donder, who assisted in the optimisation of the pilot-scale malting system after its installation and Ms Imani Deputter, who assisted in malt production and enzymatic analyses. The authors thank the Flemish Supercomputer Center (VSC), funded by the Research Foundation – Flanders (FWO) and the Flemish Government, for providing the infrastructure we used to analyse the metagenomics and metaproteomics data. The public domain images for the graphical abstract were sourced from https://openclipart.org/detail/347952/minion; https://bioart.niaid.nih.gov/bioart/638; https://bioart.niaid.nih.gov/bioart/639 and https://bioart.niaid.nih.gov/bioart/161.

## CRediT author statement

**Steven Kasozi Galiwango** – Formal analysis, Investigation, Methodology, Software, Visualization, Writing – original draft, Writing – review and editing; **Eoin O’Cinneide** – Formal analysis, Investigation, Methodology, Software; **Lars Pierco** – Formal analysis, Investigation; **Elif Göksu Kızılyar** – Formal analysis, Investigation; **Wouter De Sutter** – Formal analysis, Investigation, Methodology; **Agata Soszka** – Formal analysis, Investigation; **Jolien Philips** – Formal analysis, Investigation; **Gust Heuvinck** – Formal analysis, Investigation; **Gert De Rouck** – Conceptualization, Funding acquisition, Methodology, Resources, Supervision, Writing – review and editing; **Ilse Van de Voorde** – Conceptualization, Funding acquisition, Resources, Supervision, Writing – review and editing; **Florian Weiland** – Conceptualization, Formal analysis, Funding acquisition, Methodology, Project administration, Software, Supervision, Visualization, Writing – original draft, Writing – review and editing

## Funding

This work was supported by the KU Leuven Impuls Fund [grant number IMP/20/030] (project code: 3E211142).

## Declaration of interests

The authors declare no competing interests.

## Declaration of generative AI and AI-assisted technologies in the manuscript preparation process

During the preparation of this work, the authors used Claude (Opus 5.5, Anthropic) for language editing, formatting assistance, and troubleshooting of R scripts used for data visualisation (where also Gemini was used). After using these tools, the authors reviewed and edited the content where necessary and take full responsibility for the content of the published article.

## Data Availability

The raw mass spectrometry files of metaproteomics data generated in this study are available on ProteomeXchange via the identifier PXD086227. The barley metagenomic sequencing data is available in the NCBI SRA database accessible via https://www.ncbi.nlm.nih.gov/sra/PRJNA1541012. The raw data for enzymatic analysis and malt quality assessment were deposited on Zenodo together with their corresponding data analysis R scripts and can be accessed via the following link: https://doi.org/10.5281/zenodo.23890431. The data analysis scripts for metagenomics data can be accessed using the following Zenodo link: https://doi.org/10.5281/zenodo.24121796, whereas those for metaproteomics data analysis can be accessed via the following Zenodo link: https://doi.org/10.5281/zenodo.23995407.

## Notes

https://proteomecentral.proteomexchange.org/?pxid=PXD086227

https://www.ncbi.nlm.nih.gov/sra/PRJNA1541012

https://doi.org/10.5281/zenodo.23890431

https://doi.org/10.5281/zenodo.24121796

https://doi.org/10.5281/zenodo.23995407

### Competing Interest Statement

The authors have declared no competing interest.

