## Supplementary_Figures for "Barley malting for beer production – Impact of temperature on the barley-associated microbiome, malt quality and energy expenditure"

### Slide 1
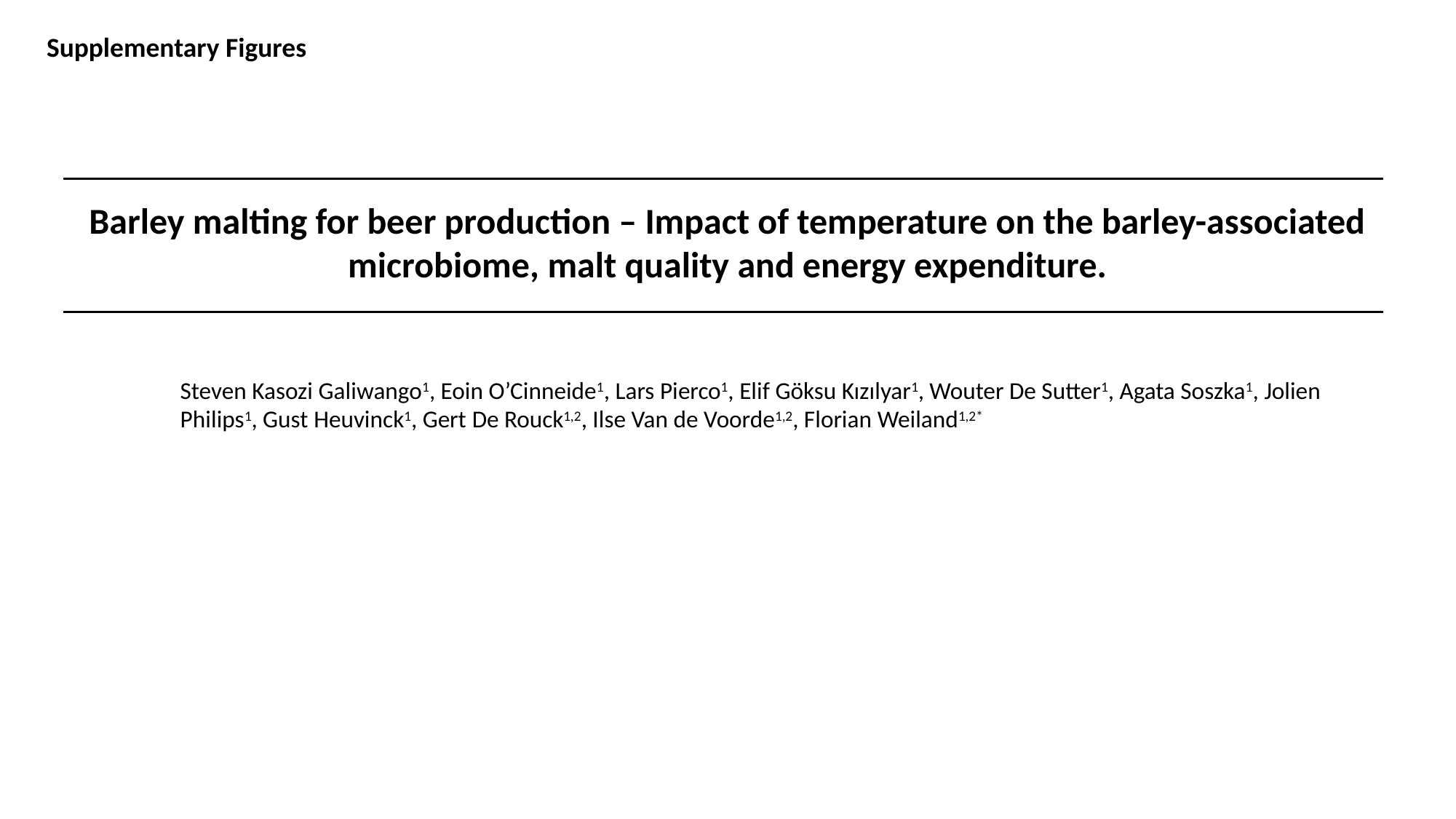

Supplementary Figures
Barley malting for beer production – Impact of temperature on the barley-associated microbiome, malt quality and energy expenditure.
Steven Kasozi Galiwango^1^, Eoin O’Cinneide^1^, Lars Pierco^1^, Elif Göksu Kızılyar^1^, Wouter De Sutter^1^, Agata Soszka^1^, Jolien Philips^1^, Gust Heuvinck^1^, Gert De Rouck^1,2^, Ilse Van de Voorde^1,2^, Florian Weiland^1,2*^

### Slide 2
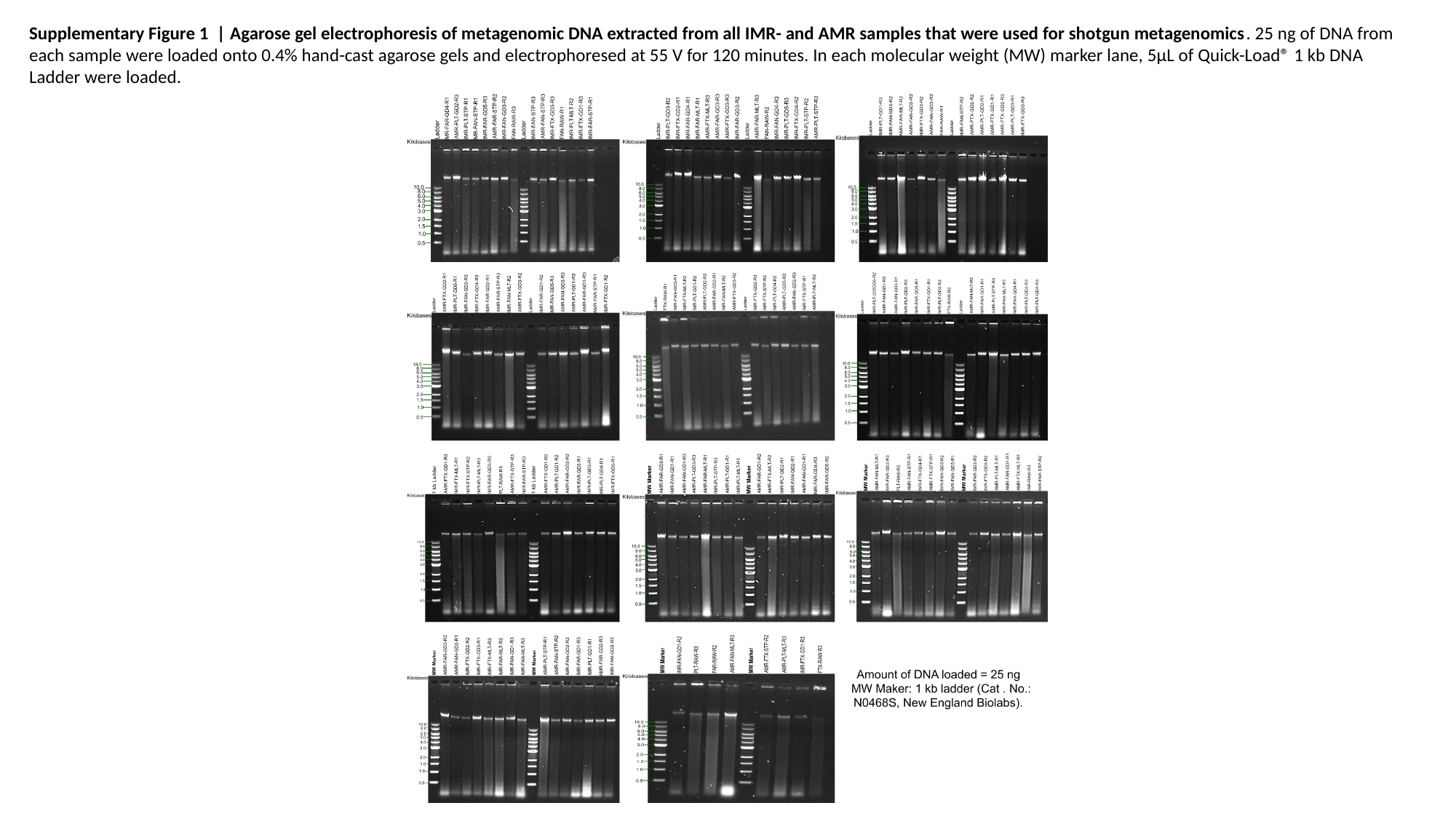

Supplementary Figure 1 | Agarose gel electrophoresis of metagenomic DNA extracted from all IMR- and AMR samples that were used for shotgun metagenomics. 25 ng of DNA from each sample were loaded onto 0.4% hand-cast agarose gels and electrophoresed at 55 V for 120 minutes. In each molecular weight (MW) marker lane, 5μL of Quick-Load® 1 kb DNA Ladder were loaded.

### Slide 3
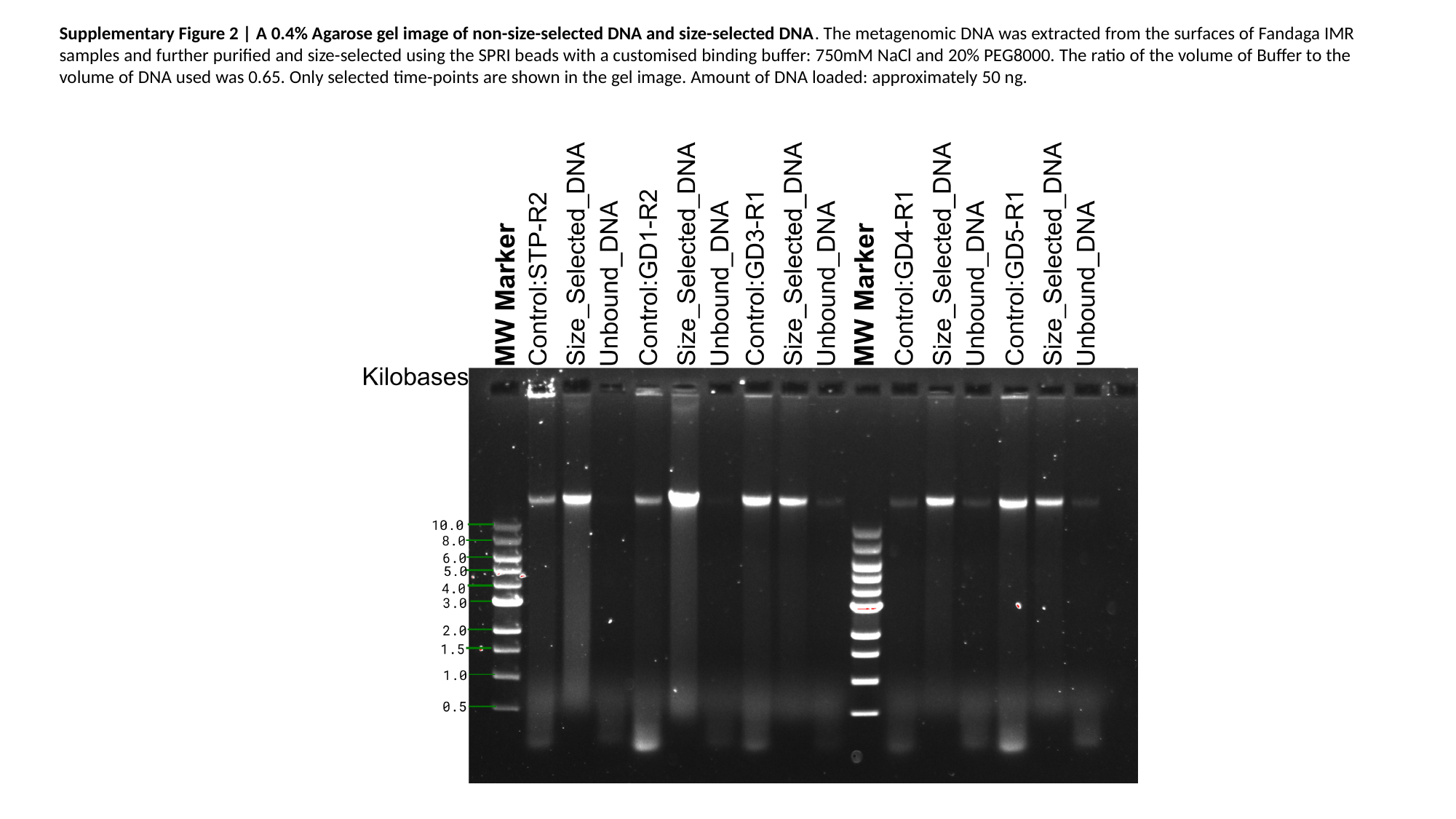

Supplementary Figure 2 | A 0.4% Agarose gel image of non-size-selected DNA and size-selected DNA. The metagenomic DNA was extracted from the surfaces of Fandaga IMR samples and further purified and size-selected using the SPRI beads with a customised binding buffer: 750mM NaCl and 20% PEG8000. The ratio of the volume of Buffer to the volume of DNA used was 0.65. Only selected time-points are shown in the gel image. Amount of DNA loaded: approximately 50 ng.

### Slide 4
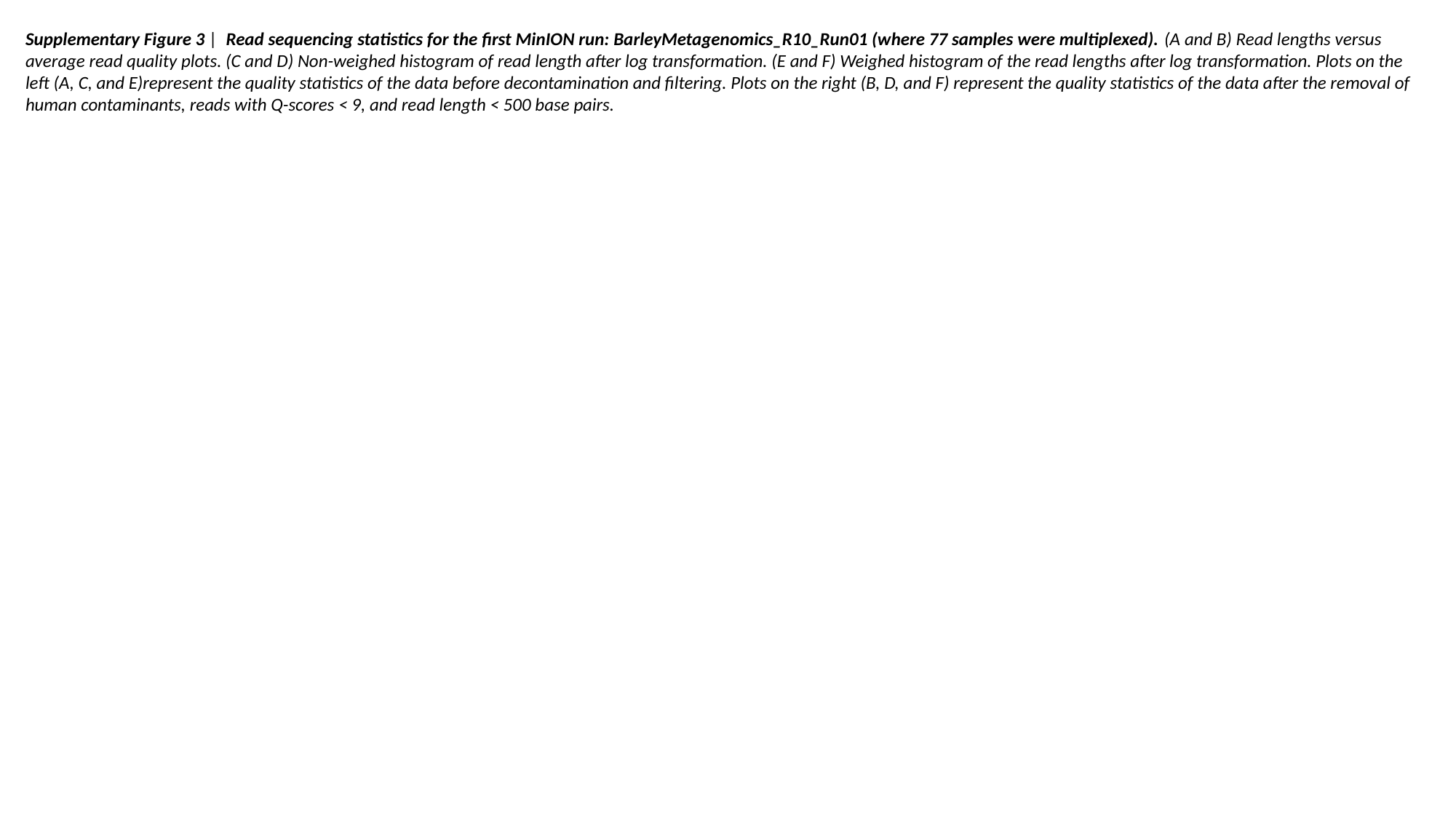

Supplementary Figure 3 | Read sequencing statistics for the first MinION run: BarleyMetagenomics_R10_Run01 (where 77 samples were multiplexed). (A and B) Read lengths versus average read quality plots. (C and D) Non-weighed histogram of read length after log transformation. (E and F) Weighed histogram of the read lengths after log transformation. Plots on the left (A, C, and E)represent the quality statistics of the data before decontamination and filtering. Plots on the right (B, D, and F) represent the quality statistics of the data after the removal of human contaminants, reads with Q-scores < 9, and read length < 500 base pairs.

### Slide 5
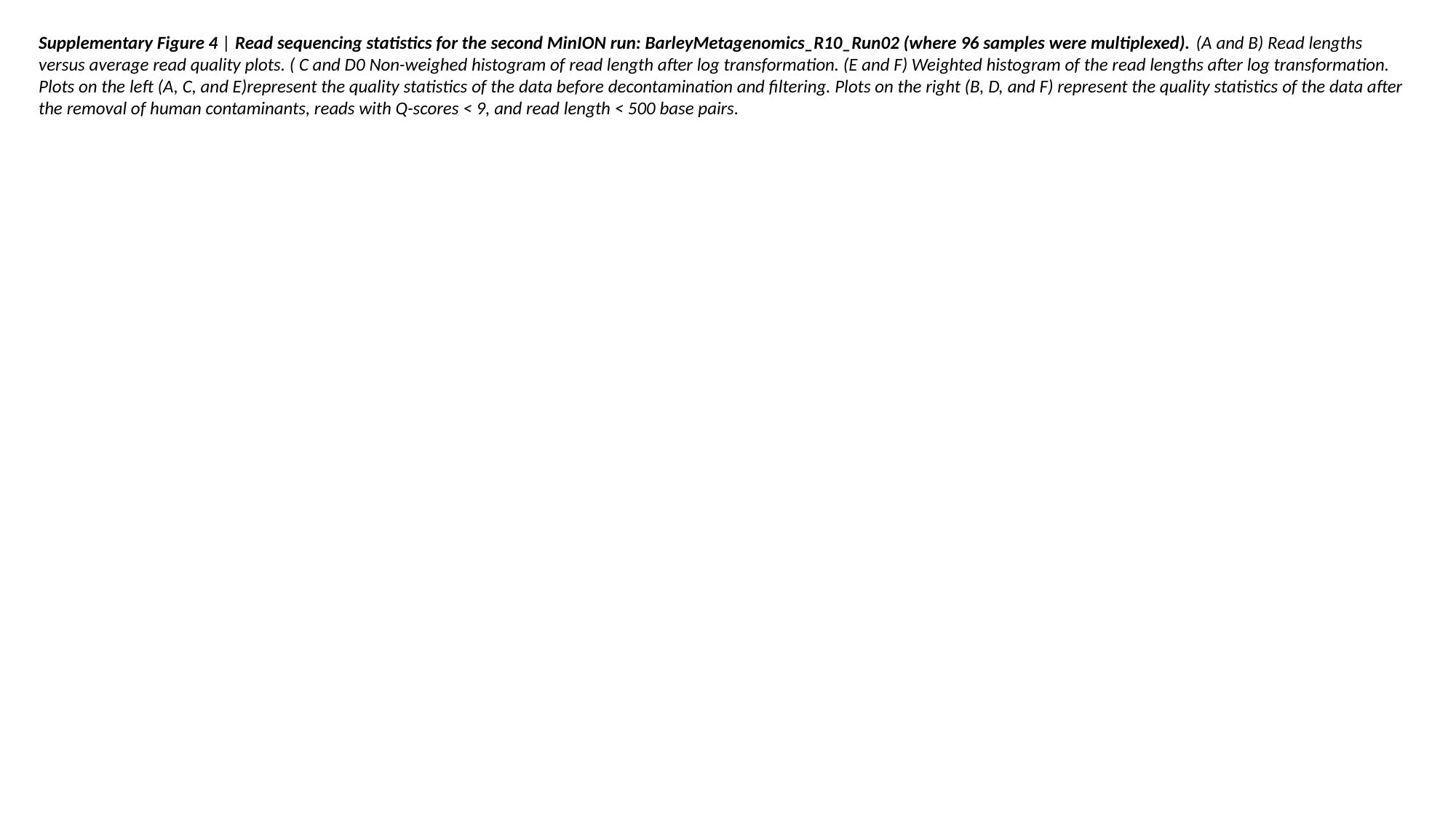

Supplementary Figure 4 | Read sequencing statistics for the second MinION run: BarleyMetagenomics_R10_Run02 (where 96 samples were multiplexed). (A and B) Read lengths versus average read quality plots. ( C and D0 Non-weighed histogram of read length after log transformation. (E and F) Weighted histogram of the read lengths after log transformation. Plots on the left (A, C, and E)represent the quality statistics of the data before decontamination and filtering. Plots on the right (B, D, and F) represent the quality statistics of the data after the removal of human contaminants, reads with Q-scores < 9, and read length < 500 base pairs.

### Slide 6
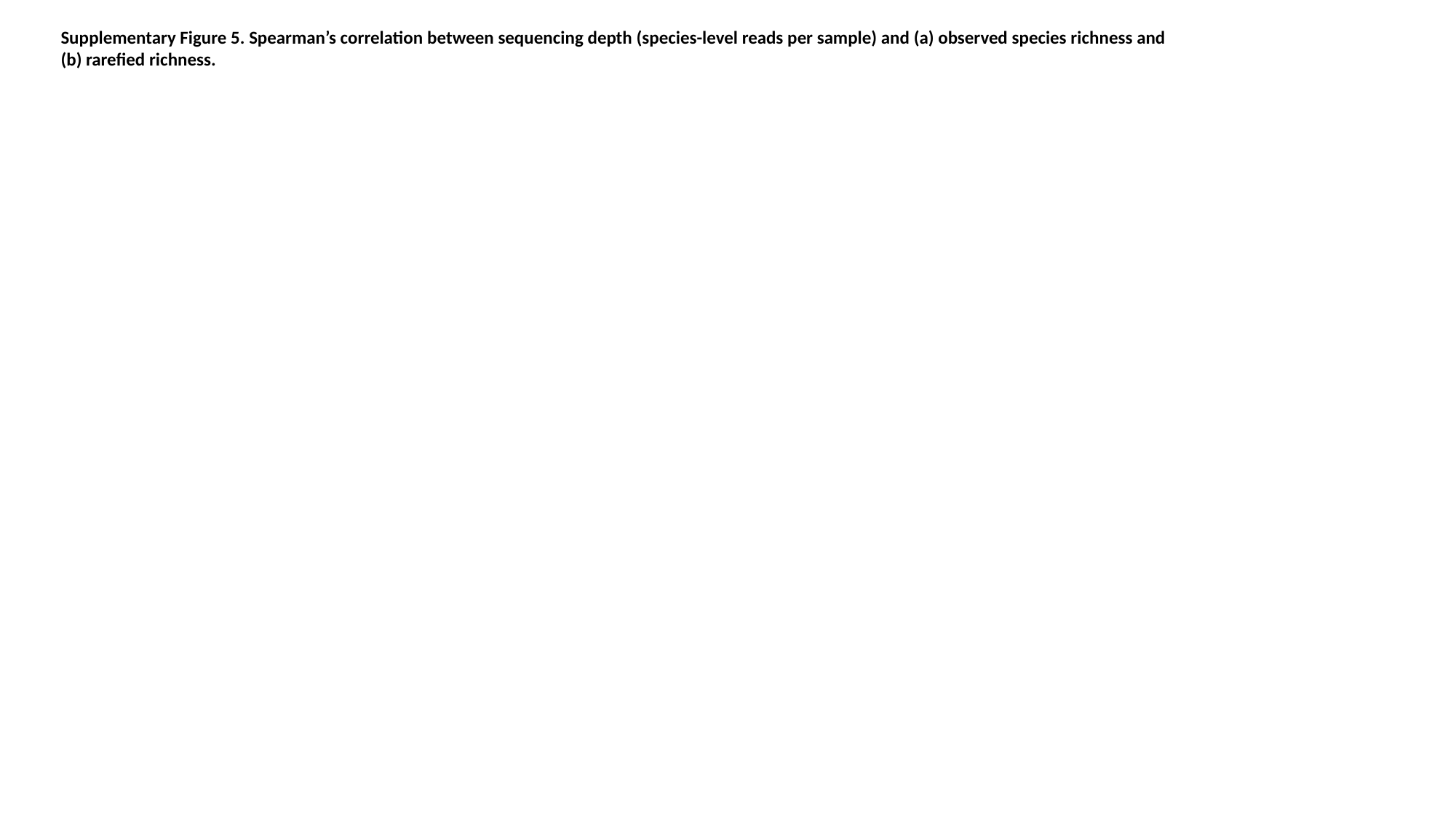

Supplementary Figure 5. Spearman’s correlation between sequencing depth (species-level reads per sample) and (a) observed species richness and (b) rarefied richness.

### Slide 7
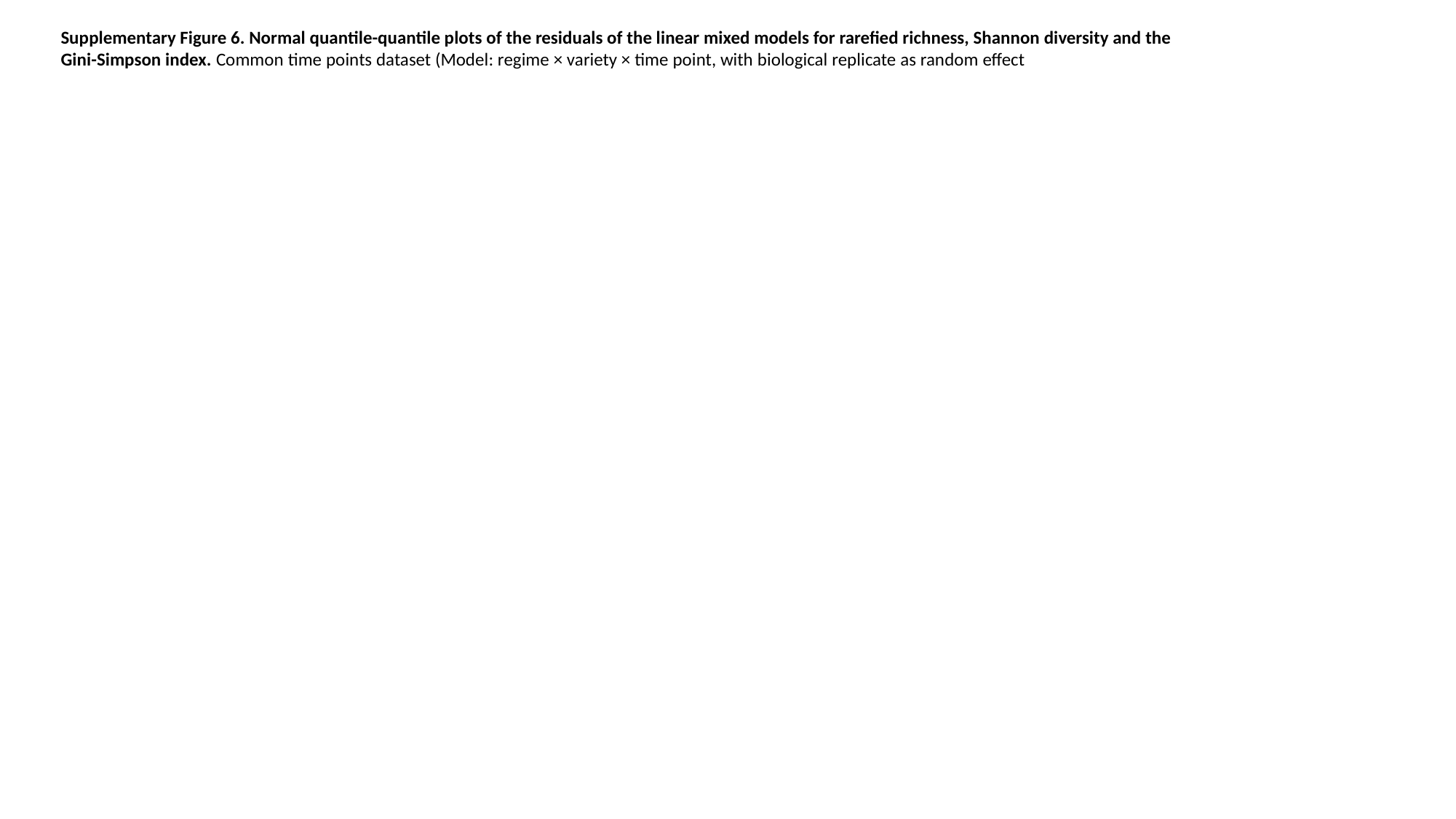

Supplementary Figure 6. Normal quantile-quantile plots of the residuals of the linear mixed models for rarefied richness, Shannon diversity and the Gini-Simpson index. Common time points dataset (Model: regime × variety × time point, with biological replicate as random effect

### Slide 8
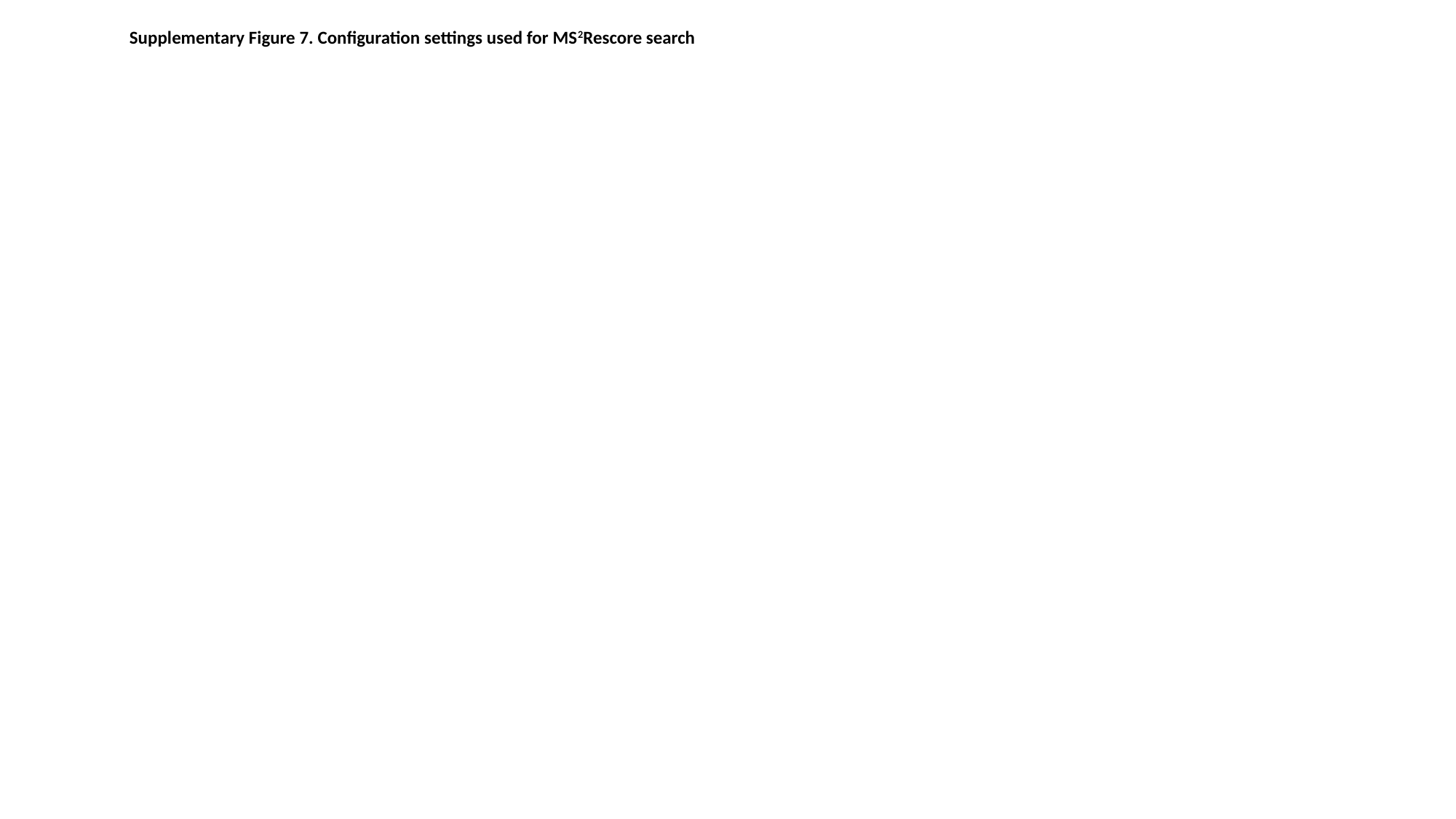

Supplementary Figure 7. Configuration settings used for MS^2^Rescore search

### Slide 9
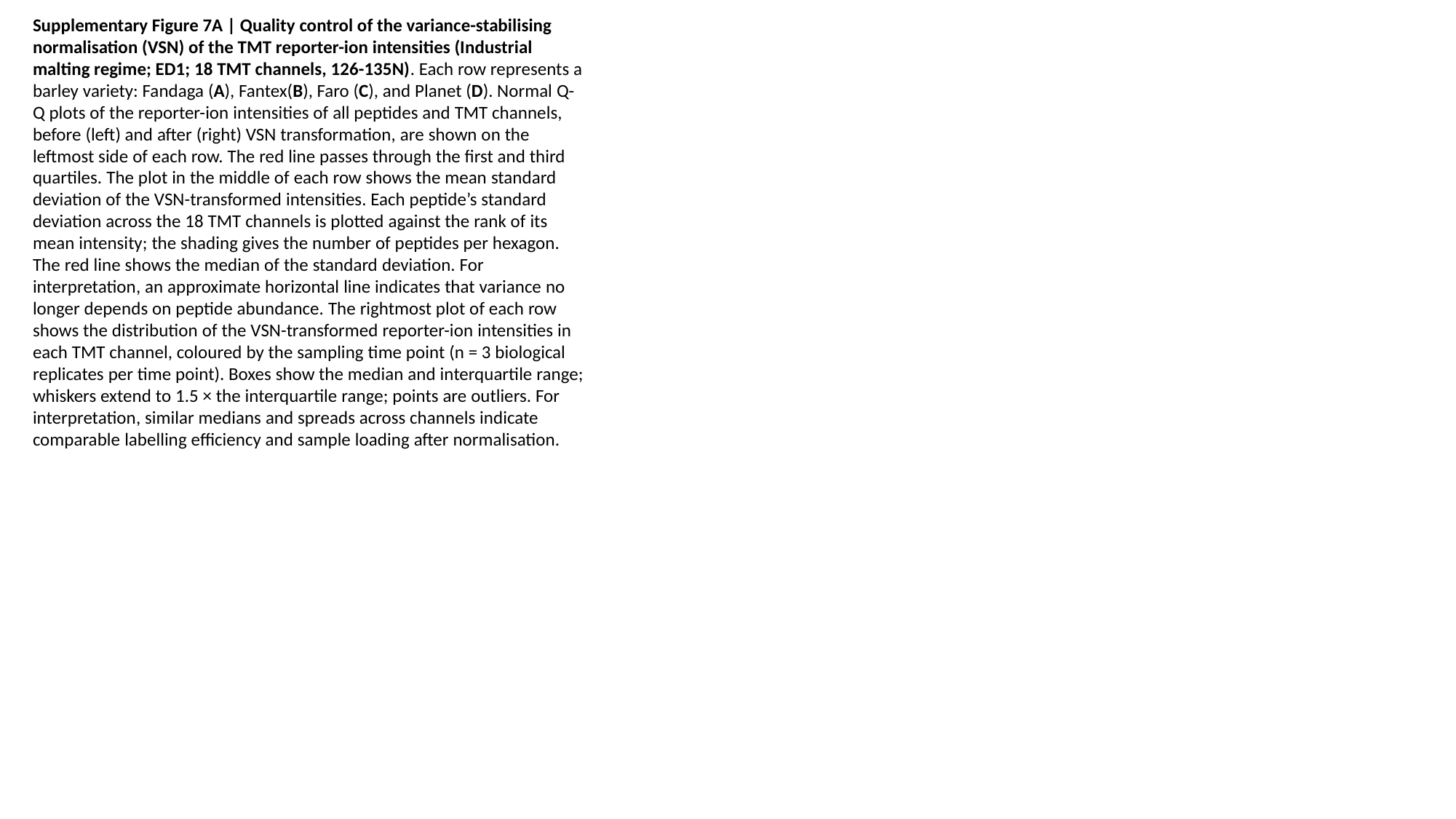

Supplementary Figure 7A | Quality control of the variance-stabilising normalisation (VSN) of the TMT reporter-ion intensities (Industrial malting regime; ED1; 18 TMT channels, 126-135N). Each row represents a barley variety: Fandaga (A), Fantex(B), Faro (C), and Planet (D). Normal Q-Q plots of the reporter-ion intensities of all peptides and TMT channels, before (left) and after (right) VSN transformation, are shown on the leftmost side of each row. The red line passes through the first and third quartiles. The plot in the middle of each row shows the mean standard deviation of the VSN-transformed intensities. Each peptide’s standard deviation across the 18 TMT channels is plotted against the rank of its mean intensity; the shading gives the number of peptides per hexagon. The red line shows the median of the standard deviation. For interpretation, an approximate horizontal line indicates that variance no longer depends on peptide abundance. The rightmost plot of each row shows the distribution of the VSN-transformed reporter-ion intensities in each TMT channel, coloured by the sampling time point (n = 3 biological replicates per time point). Boxes show the median and interquartile range; whiskers extend to 1.5 × the interquartile range; points are outliers. For interpretation, similar medians and spreads across channels indicate comparable labelling efficiency and sample loading after normalisation.

### Slide 10
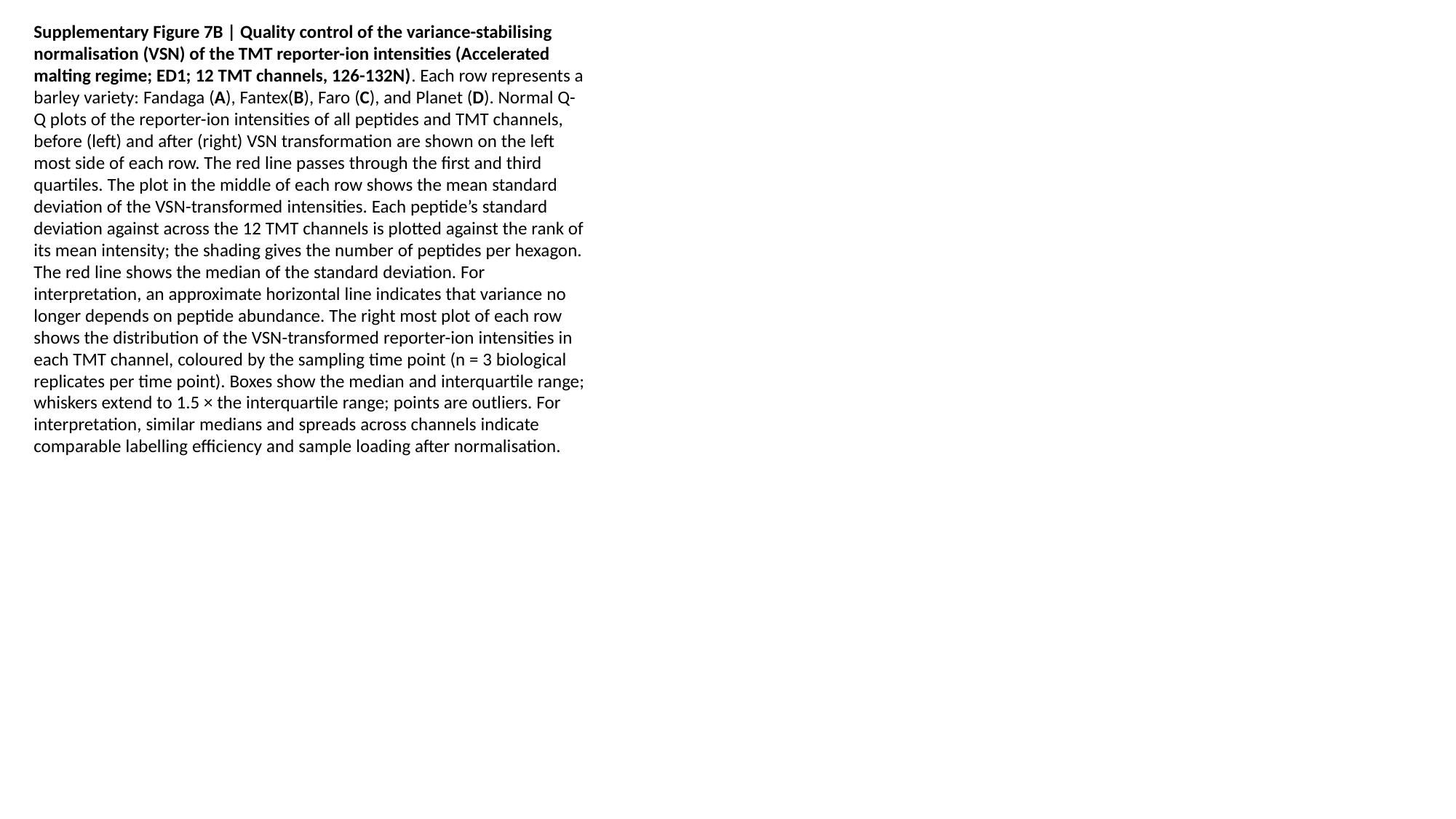

Supplementary Figure 7B | Quality control of the variance-stabilising normalisation (VSN) of the TMT reporter-ion intensities (Accelerated malting regime; ED1; 12 TMT channels, 126-132N). Each row represents a barley variety: Fandaga (A), Fantex(B), Faro (C), and Planet (D). Normal Q-Q plots of the reporter-ion intensities of all peptides and TMT channels, before (left) and after (right) VSN transformation are shown on the left most side of each row. The red line passes through the first and third quartiles. The plot in the middle of each row shows the mean standard deviation of the VSN-transformed intensities. Each peptide’s standard deviation against across the 12 TMT channels is plotted against the rank of its mean intensity; the shading gives the number of peptides per hexagon. The red line shows the median of the standard deviation. For interpretation, an approximate horizontal line indicates that variance no longer depends on peptide abundance. The right most plot of each row shows the distribution of the VSN-transformed reporter-ion intensities in each TMT channel, coloured by the sampling time point (n = 3 biological replicates per time point). Boxes show the median and interquartile range; whiskers extend to 1.5 × the interquartile range; points are outliers. For interpretation, similar medians and spreads across channels indicate comparable labelling efficiency and sample loading after normalisation.

### Slide 11
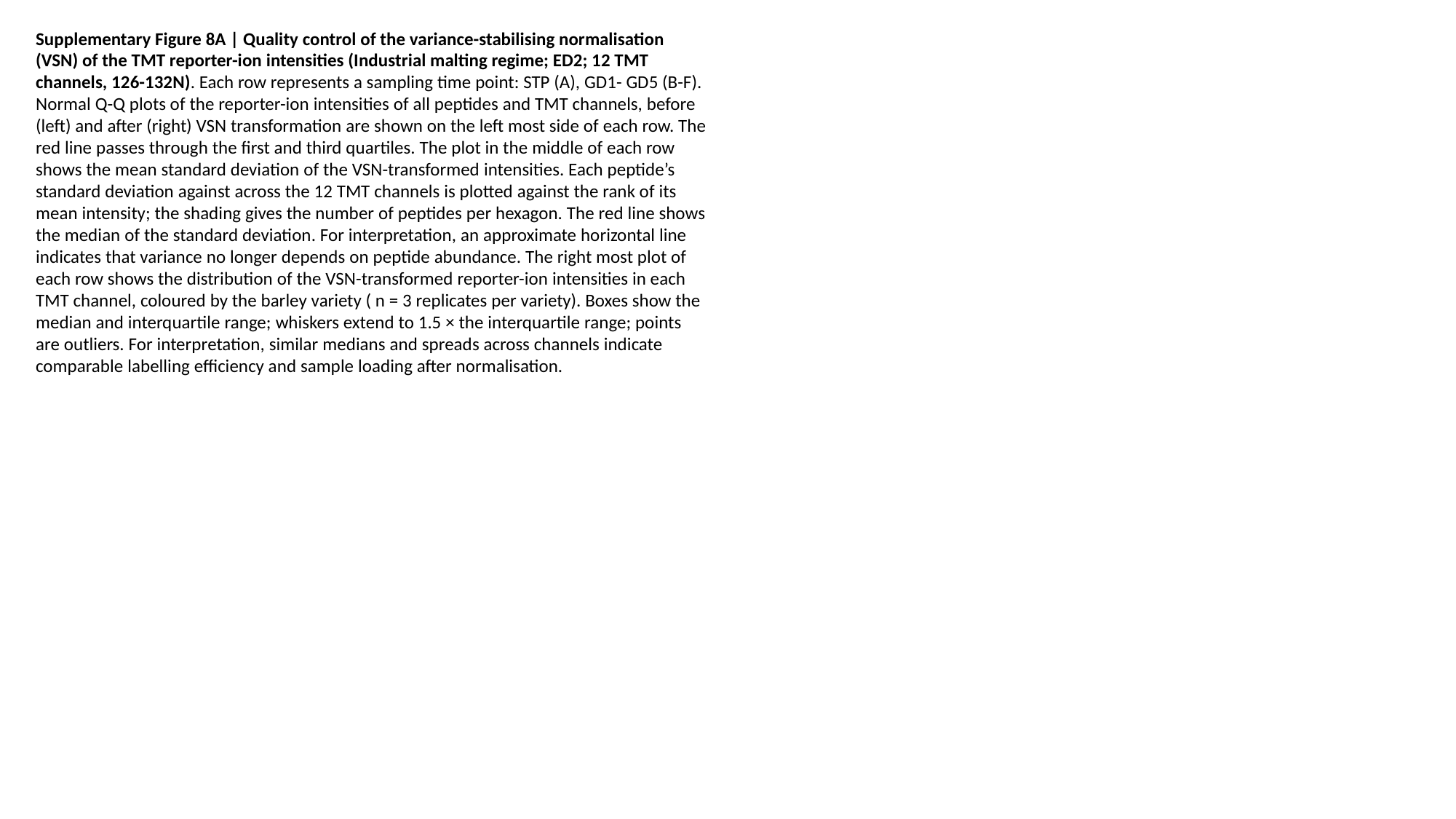

Supplementary Figure 8A | Quality control of the variance-stabilising normalisation (VSN) of the TMT reporter-ion intensities (Industrial malting regime; ED2; 12 TMT channels, 126-132N). Each row represents a sampling time point: STP (A), GD1- GD5 (B-F). Normal Q-Q plots of the reporter-ion intensities of all peptides and TMT channels, before (left) and after (right) VSN transformation are shown on the left most side of each row. The red line passes through the first and third quartiles. The plot in the middle of each row shows the mean standard deviation of the VSN-transformed intensities. Each peptide’s standard deviation against across the 12 TMT channels is plotted against the rank of its mean intensity; the shading gives the number of peptides per hexagon. The red line shows the median of the standard deviation. For interpretation, an approximate horizontal line indicates that variance no longer depends on peptide abundance. The right most plot of each row shows the distribution of the VSN-transformed reporter-ion intensities in each TMT channel, coloured by the barley variety ( n = 3 replicates per variety). Boxes show the median and interquartile range; whiskers extend to 1.5 × the interquartile range; points are outliers. For interpretation, similar medians and spreads across channels indicate comparable labelling efficiency and sample loading after normalisation.

### Slide 12
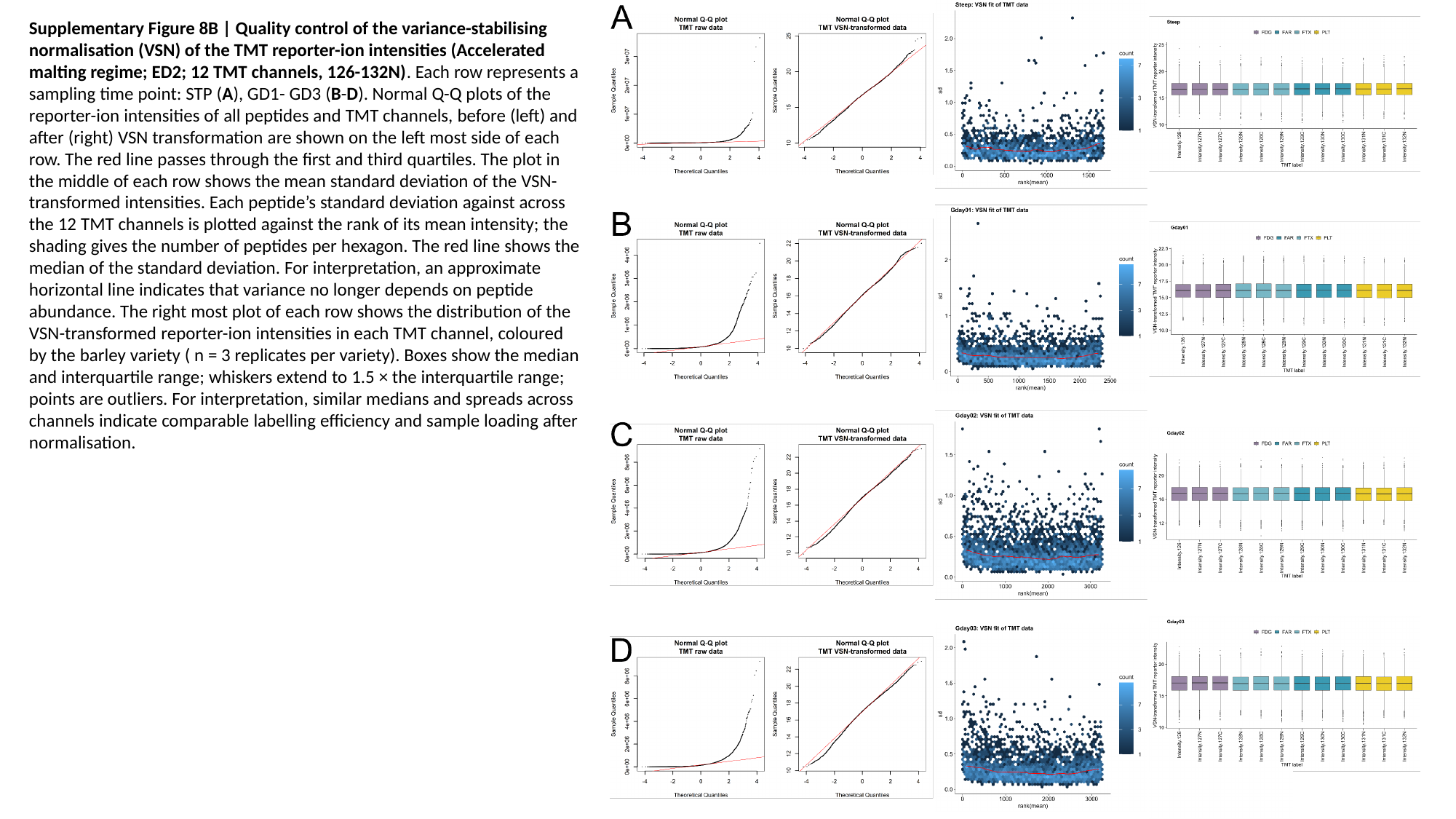

Supplementary Figure 8B | Quality control of the variance-stabilising normalisation (VSN) of the TMT reporter-ion intensities (Accelerated malting regime; ED2; 12 TMT channels, 126-132N). Each row represents a sampling time point: STP (A), GD1- GD3 (B-D). Normal Q-Q plots of the reporter-ion intensities of all peptides and TMT channels, before (left) and after (right) VSN transformation are shown on the left most side of each row. The red line passes through the first and third quartiles. The plot in the middle of each row shows the mean standard deviation of the VSN-transformed intensities. Each peptide’s standard deviation against across the 12 TMT channels is plotted against the rank of its mean intensity; the shading gives the number of peptides per hexagon. The red line shows the median of the standard deviation. For interpretation, an approximate horizontal line indicates that variance no longer depends on peptide abundance. The right most plot of each row shows the distribution of the VSN-transformed reporter-ion intensities in each TMT channel, coloured by the barley variety ( n = 3 replicates per variety). Boxes show the median and interquartile range; whiskers extend to 1.5 × the interquartile range; points are outliers. For interpretation, similar medians and spreads across channels indicate comparable labelling efficiency and sample loading after normalisation.

### Slide 13
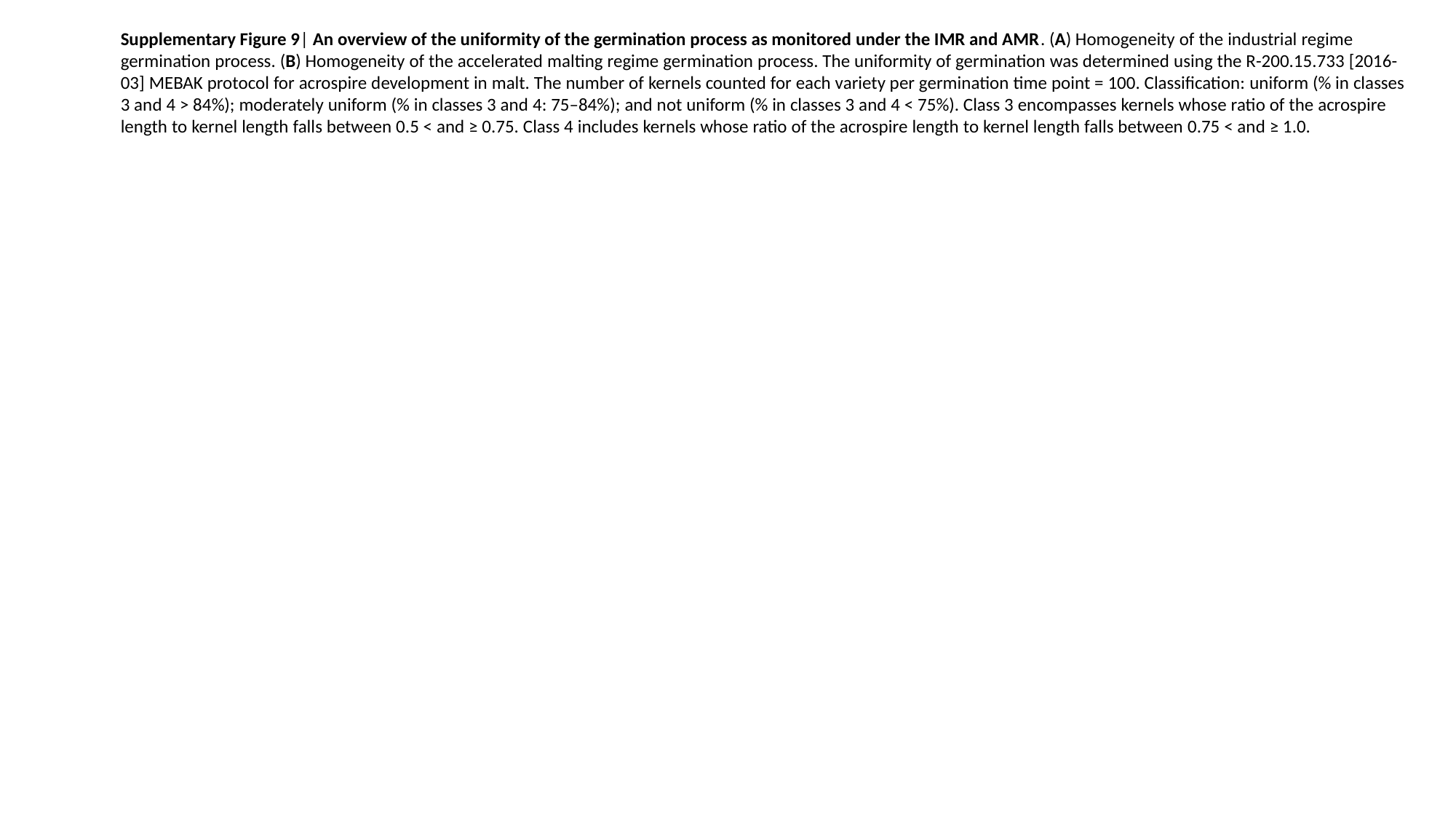

Supplementary Figure 9| An overview of the uniformity of the germination process as monitored under the IMR and AMR. (A) Homogeneity of the industrial regime germination process. (B) Homogeneity of the accelerated malting regime germination process. The uniformity of germination was determined using the R-200.15.733 [2016-03] MEBAK protocol for acrospire development in malt. The number of kernels counted for each variety per germination time point = 100. Classification: uniform (% in classes 3 and 4 > 84%); moderately uniform (% in classes 3 and 4: 75–84%); and not uniform (% in classes 3 and 4 < 75%). Class 3 encompasses kernels whose ratio of the acrospire length to kernel length falls between 0.5 < and ≥ 0.75. Class 4 includes kernels whose ratio of the acrospire length to kernel length falls between 0.75 < and ≥ 1.0.

### Slide 14
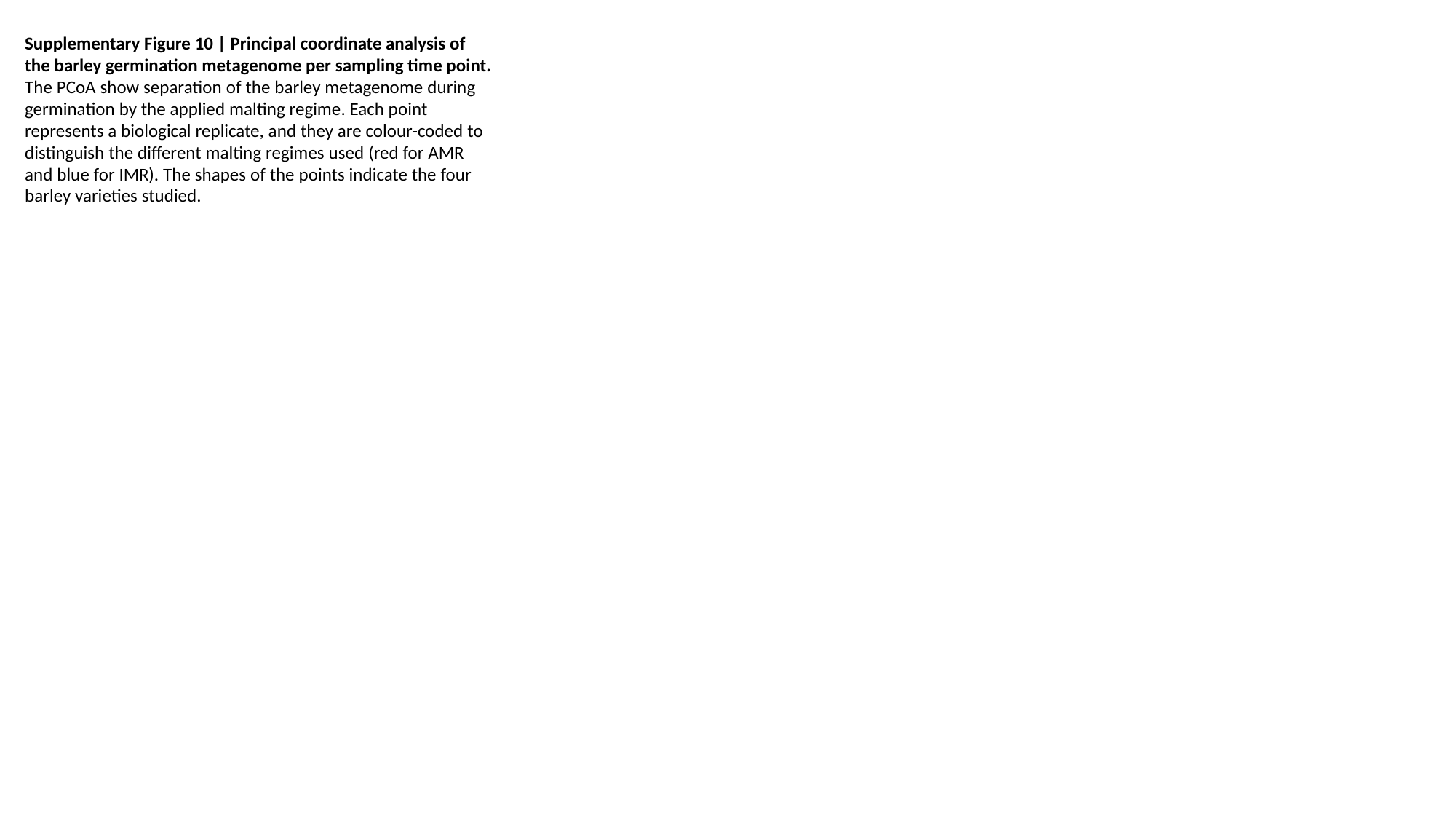

Supplementary Figure 10 | Principal coordinate analysis of the barley germination metagenome per sampling time point. The PCoA show separation of the barley metagenome during germination by the applied malting regime. Each point represents a biological replicate, and they are colour-coded to distinguish the different malting regimes used (red for AMR and blue for IMR). The shapes of the points indicate the four barley varieties studied.

### Slide 15
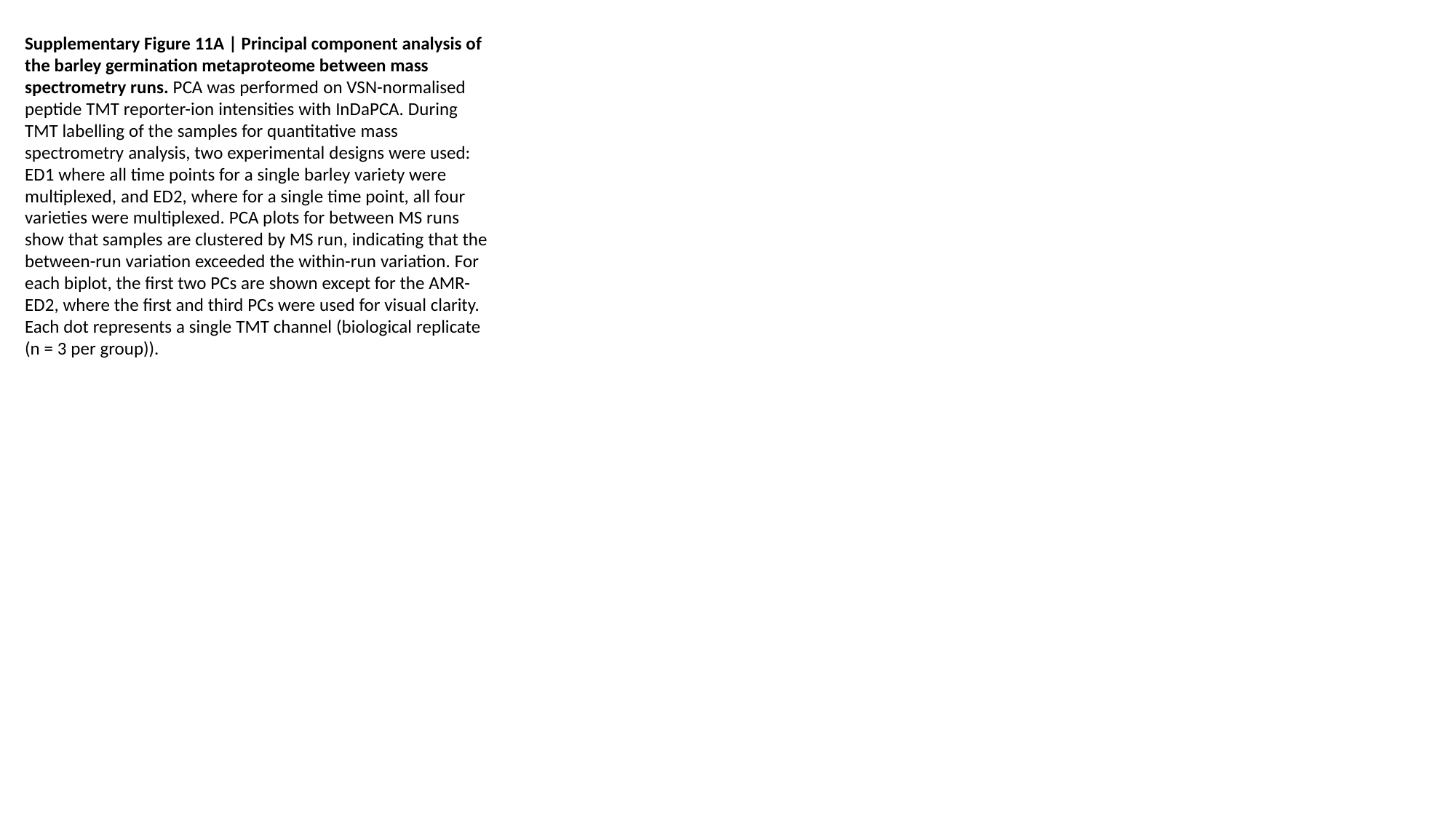

Supplementary Figure 11A | Principal component analysis of the barley germination metaproteome between mass spectrometry runs. PCA was performed on VSN-normalised peptide TMT reporter-ion intensities with InDaPCA. During TMT labelling of the samples for quantitative mass spectrometry analysis, two experimental designs were used: ED1 where all time points for a single barley variety were multiplexed, and ED2, where for a single time point, all four varieties were multiplexed. PCA plots for between MS runs show that samples are clustered by MS run, indicating that the between-run variation exceeded the within-run variation. For each biplot, the first two PCs are shown except for the AMR-ED2, where the first and third PCs were used for visual clarity. Each dot represents a single TMT channel (biological replicate (n = 3 per group)).

### Slide 16
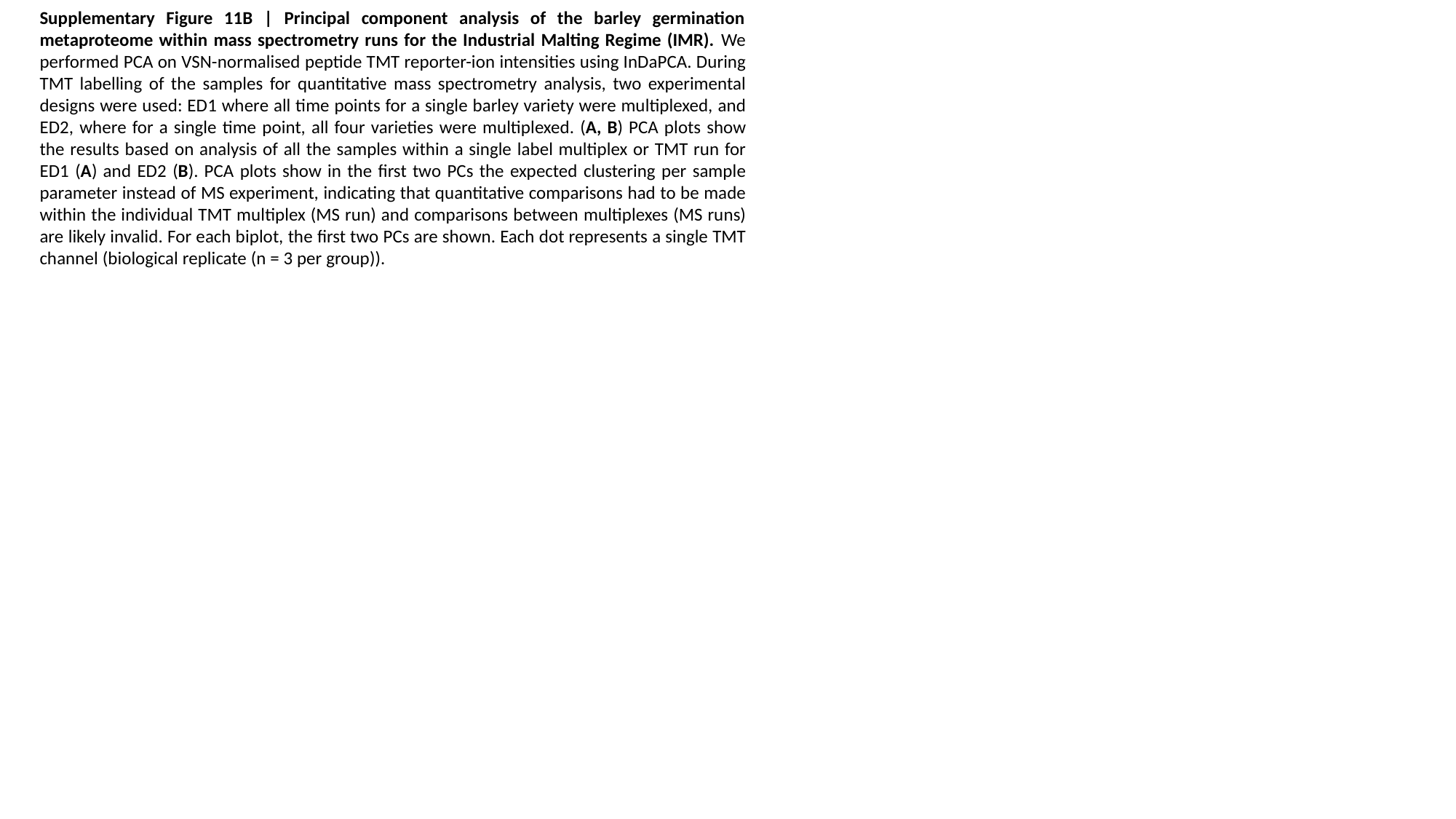

Supplementary Figure 11B | Principal component analysis of the barley germination metaproteome within mass spectrometry runs for the Industrial Malting Regime (IMR). We performed PCA on VSN-normalised peptide TMT reporter-ion intensities using InDaPCA. During TMT labelling of the samples for quantitative mass spectrometry analysis, two experimental designs were used: ED1 where all time points for a single barley variety were multiplexed, and ED2, where for a single time point, all four varieties were multiplexed. (A, B) PCA plots show the results based on analysis of all the samples within a single label multiplex or TMT run for ED1 (A) and ED2 (B). PCA plots show in the first two PCs the expected clustering per sample parameter instead of MS experiment, indicating that quantitative comparisons had to be made within the individual TMT multiplex (MS run) and comparisons between multiplexes (MS runs) are likely invalid. For each biplot, the first two PCs are shown. Each dot represents a single TMT channel (biological replicate (n = 3 per group)).

### Slide 17
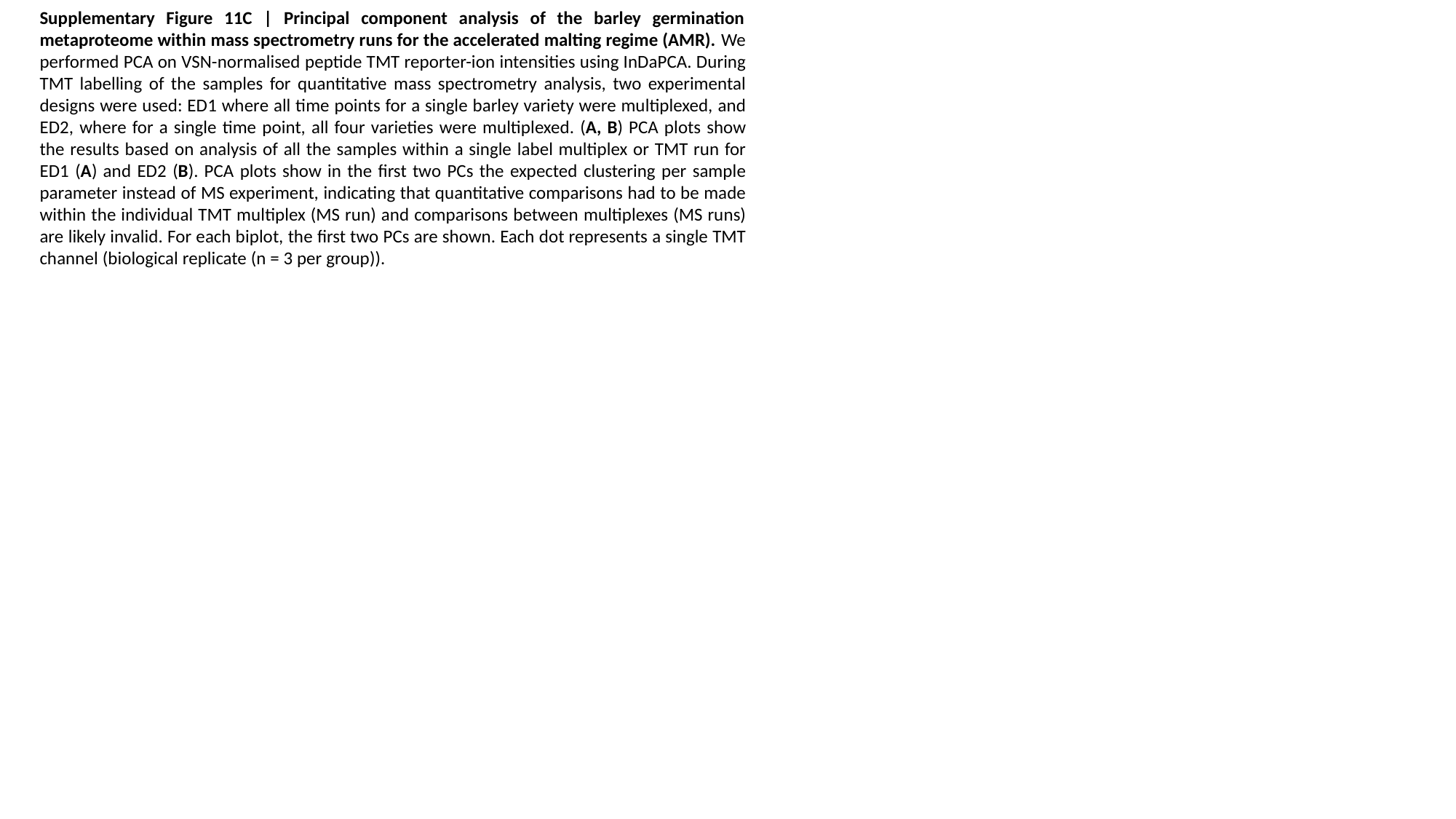

Supplementary Figure 11C | Principal component analysis of the barley germination metaproteome within mass spectrometry runs for the accelerated malting regime (AMR). We performed PCA on VSN-normalised peptide TMT reporter-ion intensities using InDaPCA. During TMT labelling of the samples for quantitative mass spectrometry analysis, two experimental designs were used: ED1 where all time points for a single barley variety were multiplexed, and ED2, where for a single time point, all four varieties were multiplexed. (A, B) PCA plots show the results based on analysis of all the samples within a single label multiplex or TMT run for ED1 (A) and ED2 (B). PCA plots show in the first two PCs the expected clustering per sample parameter instead of MS experiment, indicating that quantitative comparisons had to be made within the individual TMT multiplex (MS run) and comparisons between multiplexes (MS runs) are likely invalid. For each biplot, the first two PCs are shown. Each dot represents a single TMT channel (biological replicate (n = 3 per group)).

### Slide 18
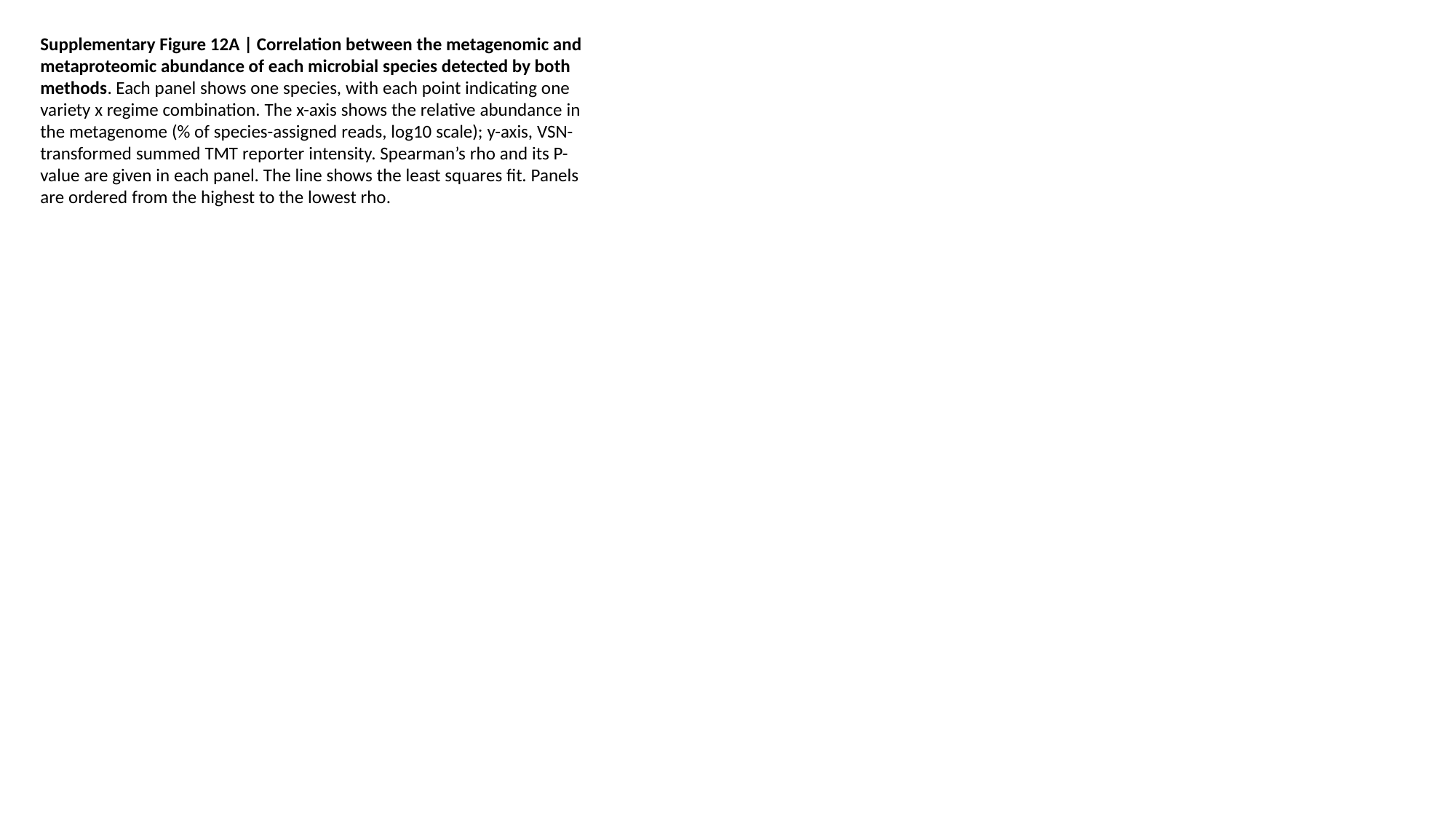

Supplementary Figure 12A | Correlation between the metagenomic and metaproteomic abundance of each microbial species detected by both methods. Each panel shows one species, with each point indicating one variety x regime combination. The x-axis shows the relative abundance in the metagenome (% of species-assigned reads, log10 scale); y-axis, VSN-transformed summed TMT reporter intensity. Spearman’s rho and its P-value are given in each panel. The line shows the least squares fit. Panels are ordered from the highest to the lowest rho.

### Slide 19
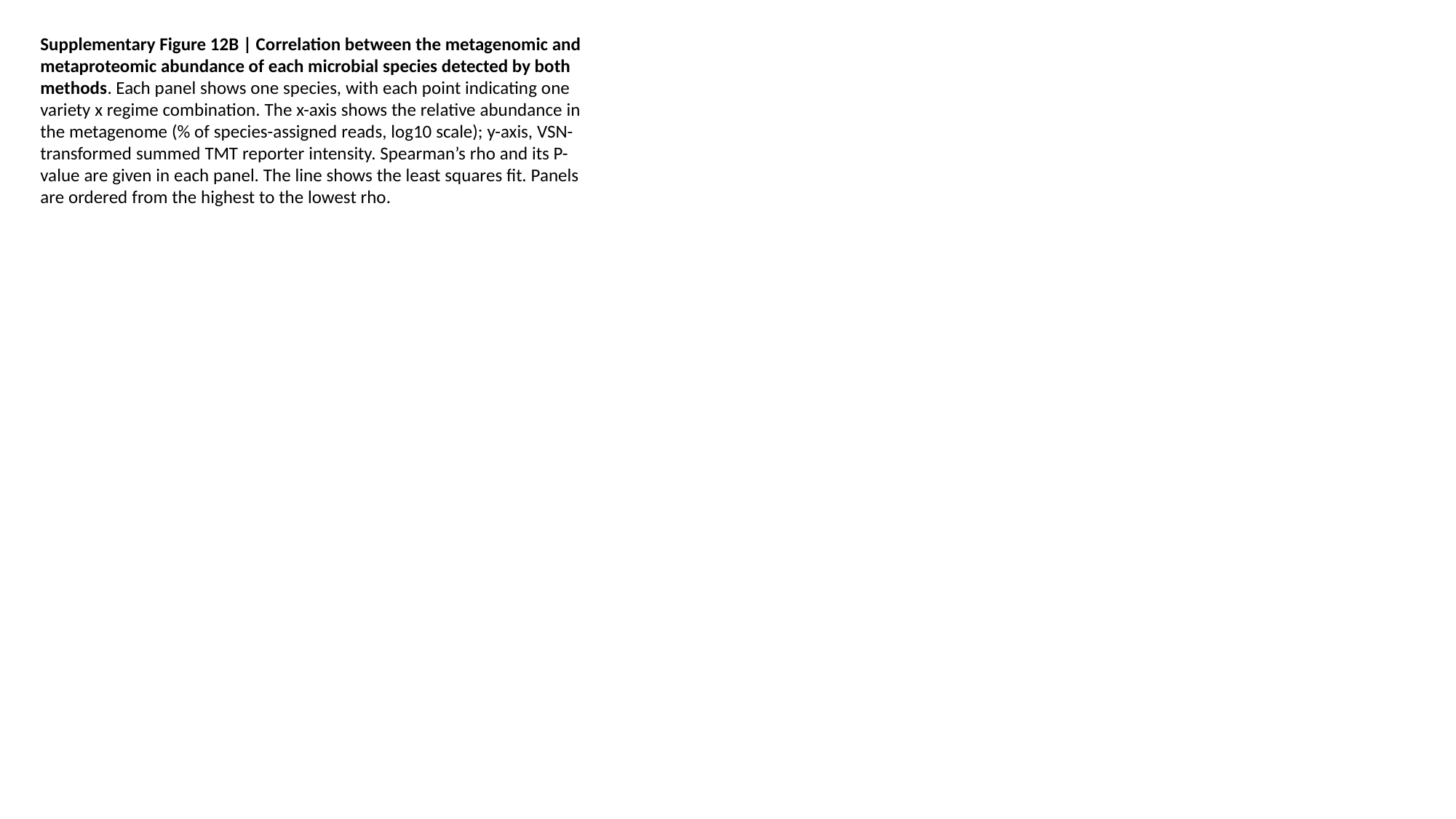

Supplementary Figure 12B | Correlation between the metagenomic and metaproteomic abundance of each microbial species detected by both methods. Each panel shows one species, with each point indicating one variety x regime combination. The x-axis shows the relative abundance in the metagenome (% of species-assigned reads, log10 scale); y-axis, VSN-transformed summed TMT reporter intensity. Spearman’s rho and its P-value are given in each panel. The line shows the least squares fit. Panels are ordered from the highest to the lowest rho.

### Slide 20
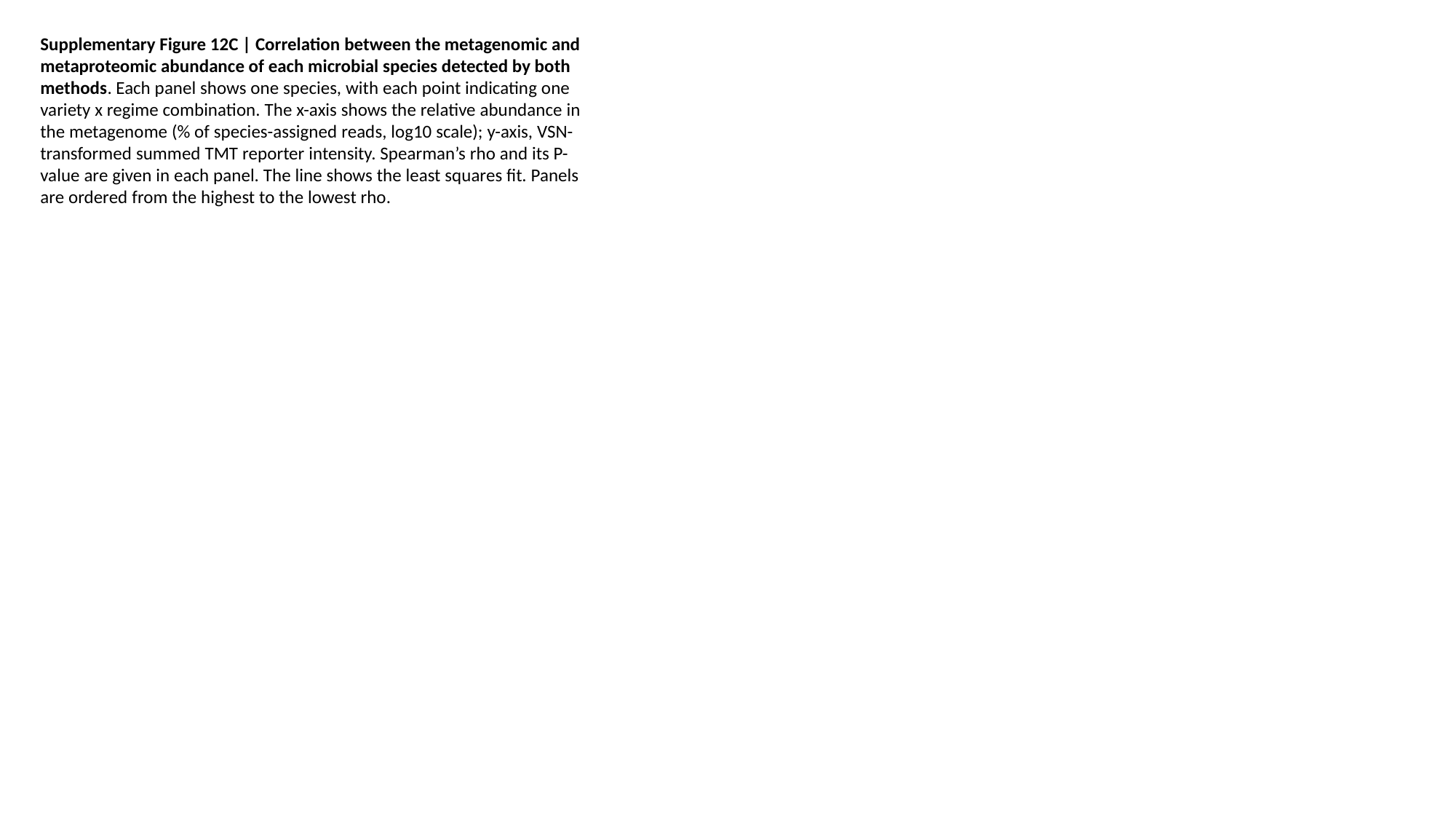

Supplementary Figure 12C | Correlation between the metagenomic and metaproteomic abundance of each microbial species detected by both methods. Each panel shows one species, with each point indicating one variety x regime combination. The x-axis shows the relative abundance in the metagenome (% of species-assigned reads, log10 scale); y-axis, VSN-transformed summed TMT reporter intensity. Spearman’s rho and its P-value are given in each panel. The line shows the least squares fit. Panels are ordered from the highest to the lowest rho.

### Slide 21
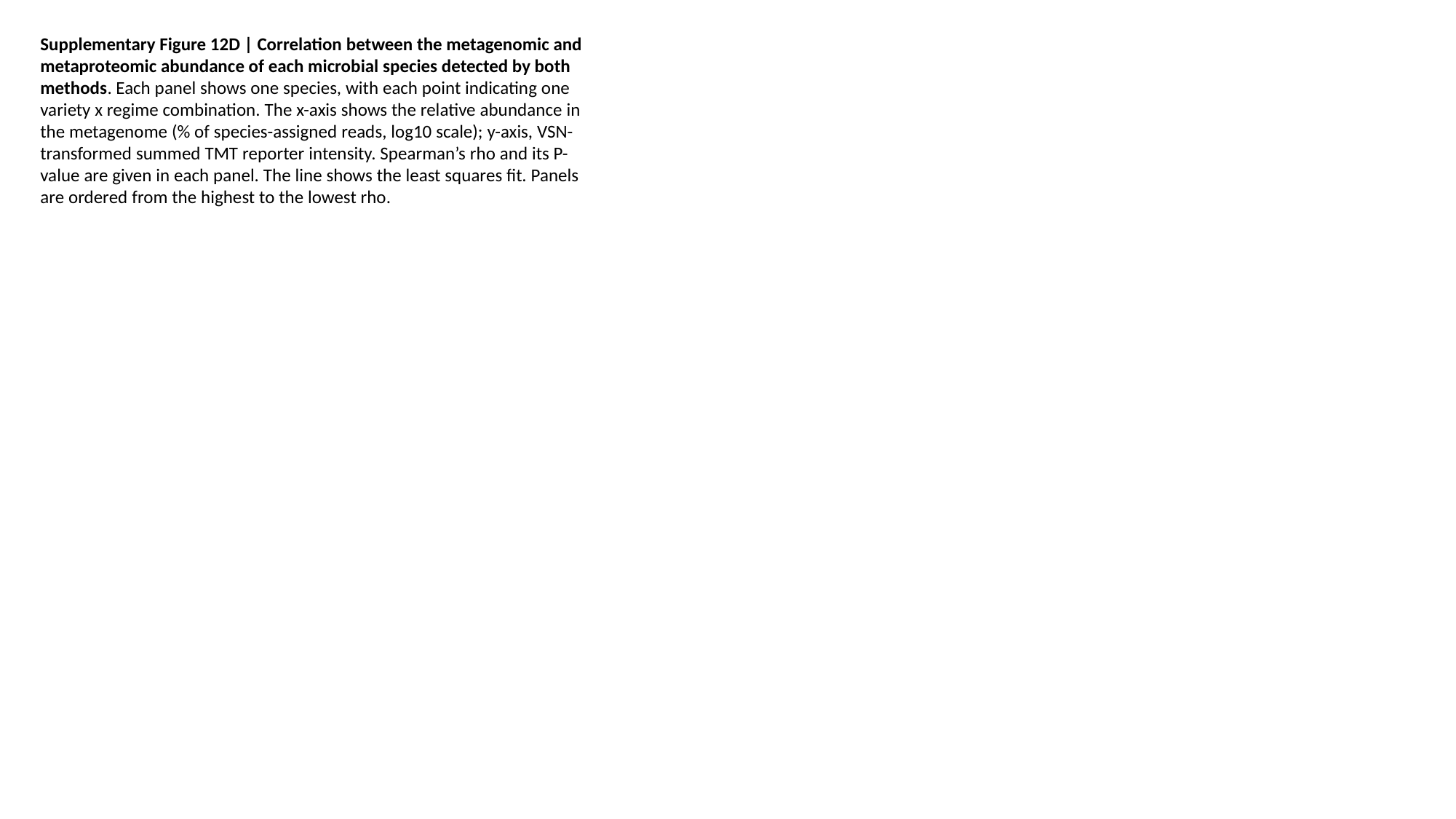

Supplementary Figure 12D | Correlation between the metagenomic and metaproteomic abundance of each microbial species detected by both methods. Each panel shows one species, with each point indicating one variety x regime combination. The x-axis shows the relative abundance in the metagenome (% of species-assigned reads, log10 scale); y-axis, VSN-transformed summed TMT reporter intensity. Spearman’s rho and its P-value are given in each panel. The line shows the least squares fit. Panels are ordered from the highest to the lowest rho.

### Slide 22
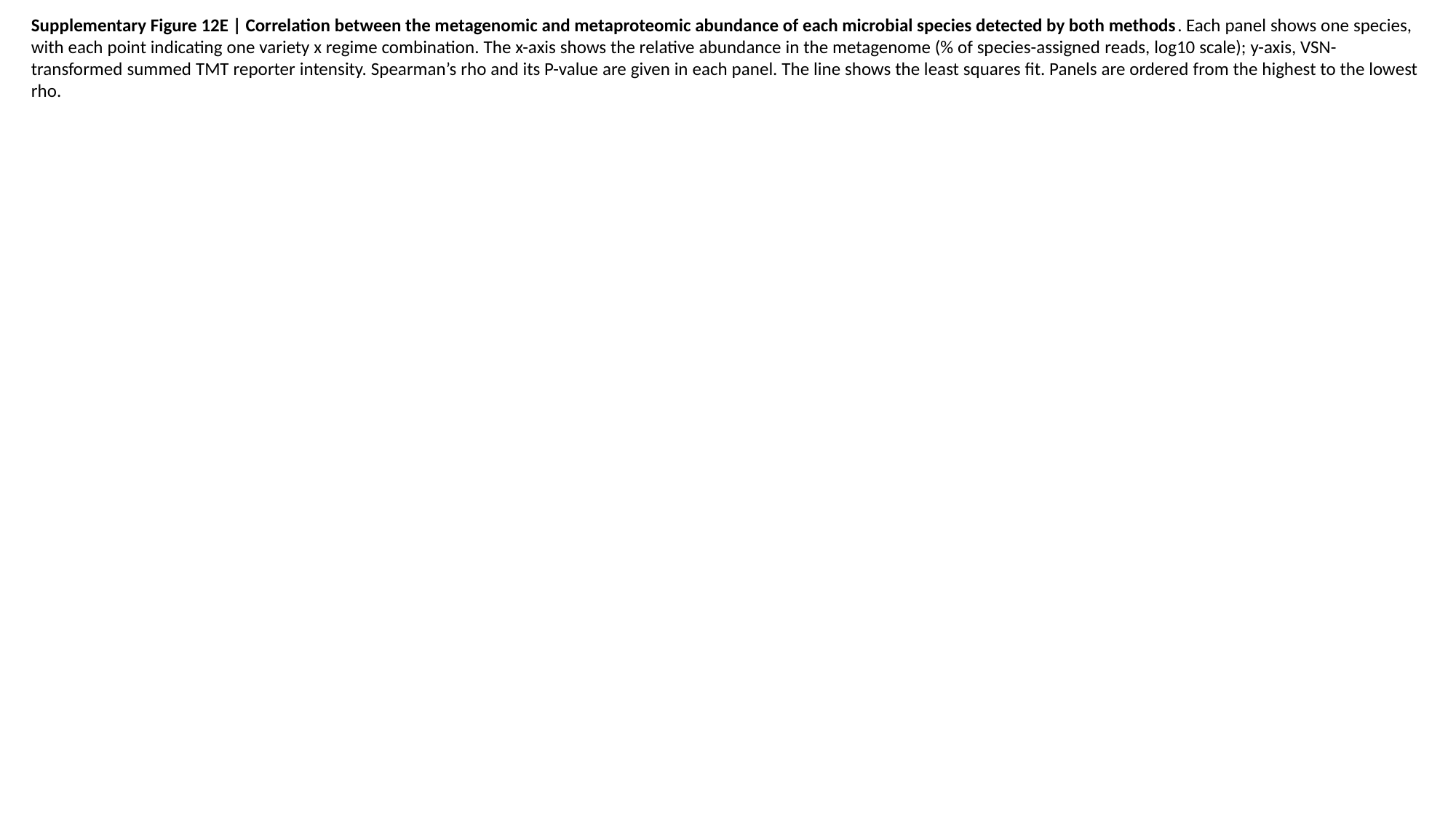

Supplementary Figure 12E | Correlation between the metagenomic and metaproteomic abundance of each microbial species detected by both methods. Each panel shows one species, with each point indicating one variety x regime combination. The x-axis shows the relative abundance in the metagenome (% of species-assigned reads, log10 scale); y-axis, VSN-transformed summed TMT reporter intensity. Spearman’s rho and its P-value are given in each panel. The line shows the least squares fit. Panels are ordered from the highest to the lowest rho.

### Slide 23
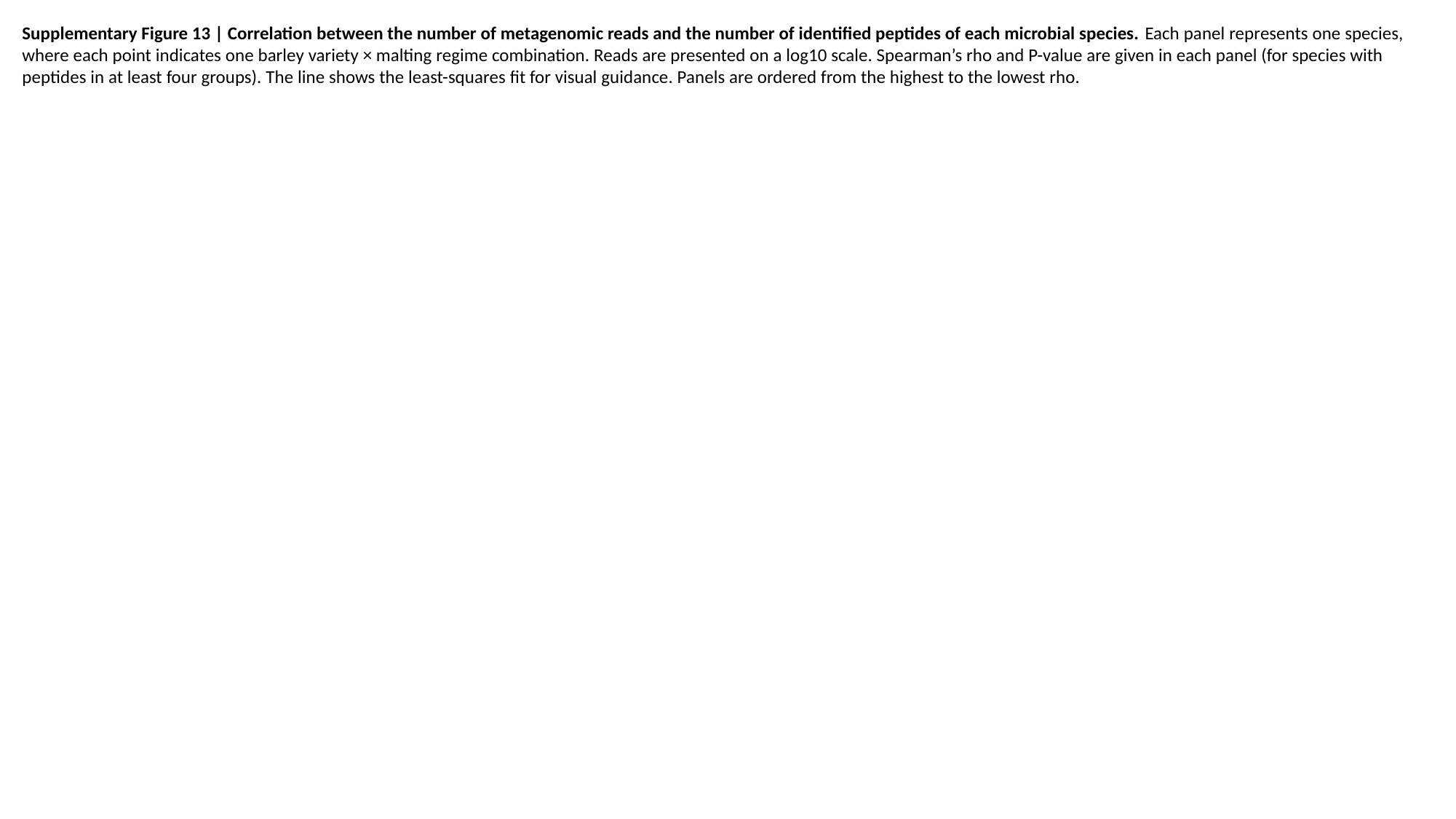

Supplementary Figure 13 | Correlation between the number of metagenomic reads and the number of identified peptides of each microbial species. Each panel represents one species, where each point indicates one barley variety × malting regime combination. Reads are presented on a log10 scale. Spearman’s rho and P-value are given in each panel (for species with peptides in at least four groups). The line shows the least-squares fit for visual guidance. Panels are ordered from the highest to the lowest rho.

### Slide 24
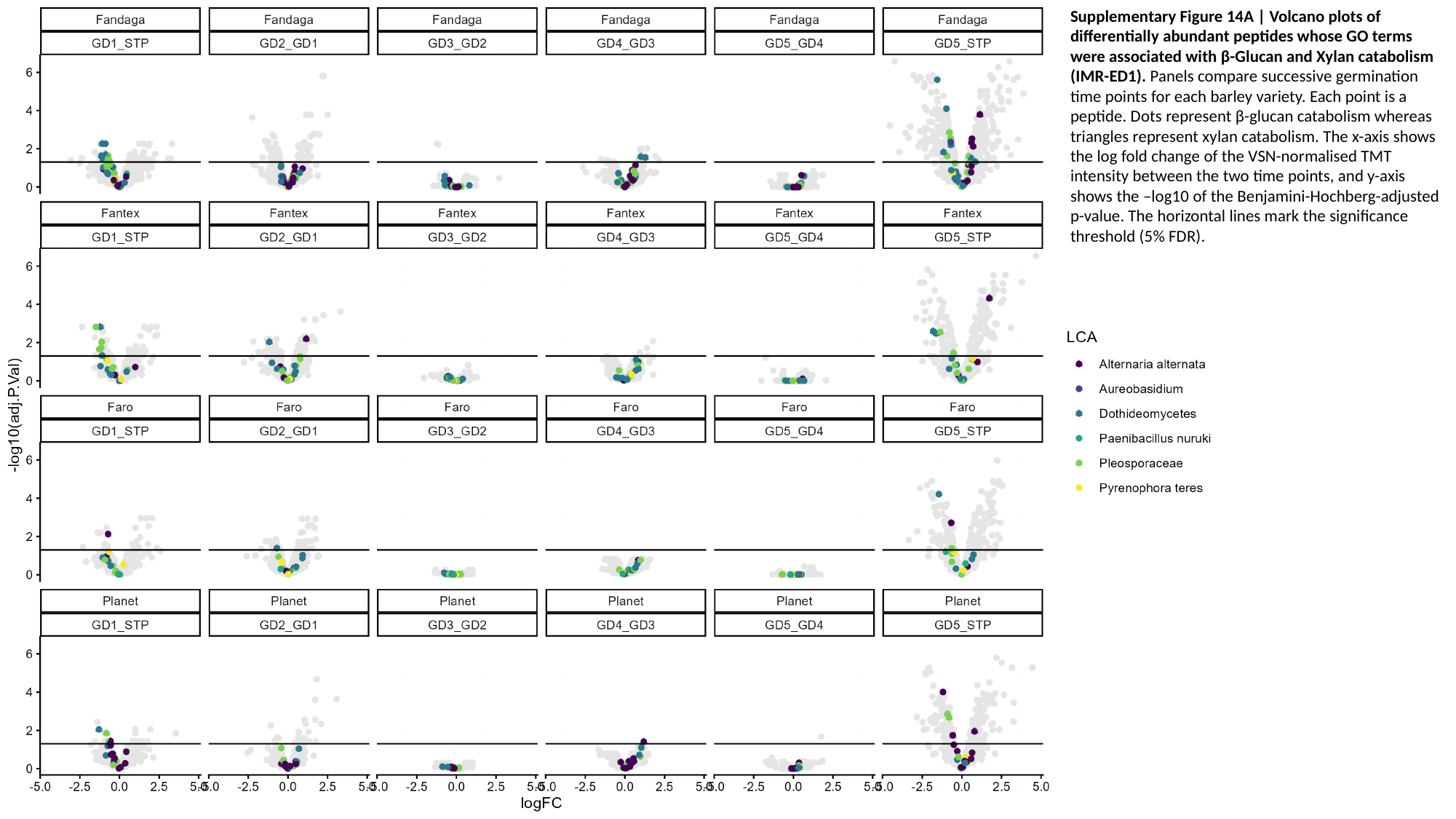

Supplementary Figure 14A | Volcano plots of differentially abundant peptides whose GO terms were associated with β-Glucan and Xylan catabolism (IMR-ED1). Panels compare successive germination time points for each barley variety. Each point is a peptide. Dots represent β-glucan catabolism whereas triangles represent xylan catabolism. The x-axis shows the log fold change of the VSN-normalised TMT intensity between the two time points, and y-axis shows the –log10 of the Benjamini-Hochberg-adjusted p-value. The horizontal lines mark the significance threshold (5% FDR).

### Slide 25
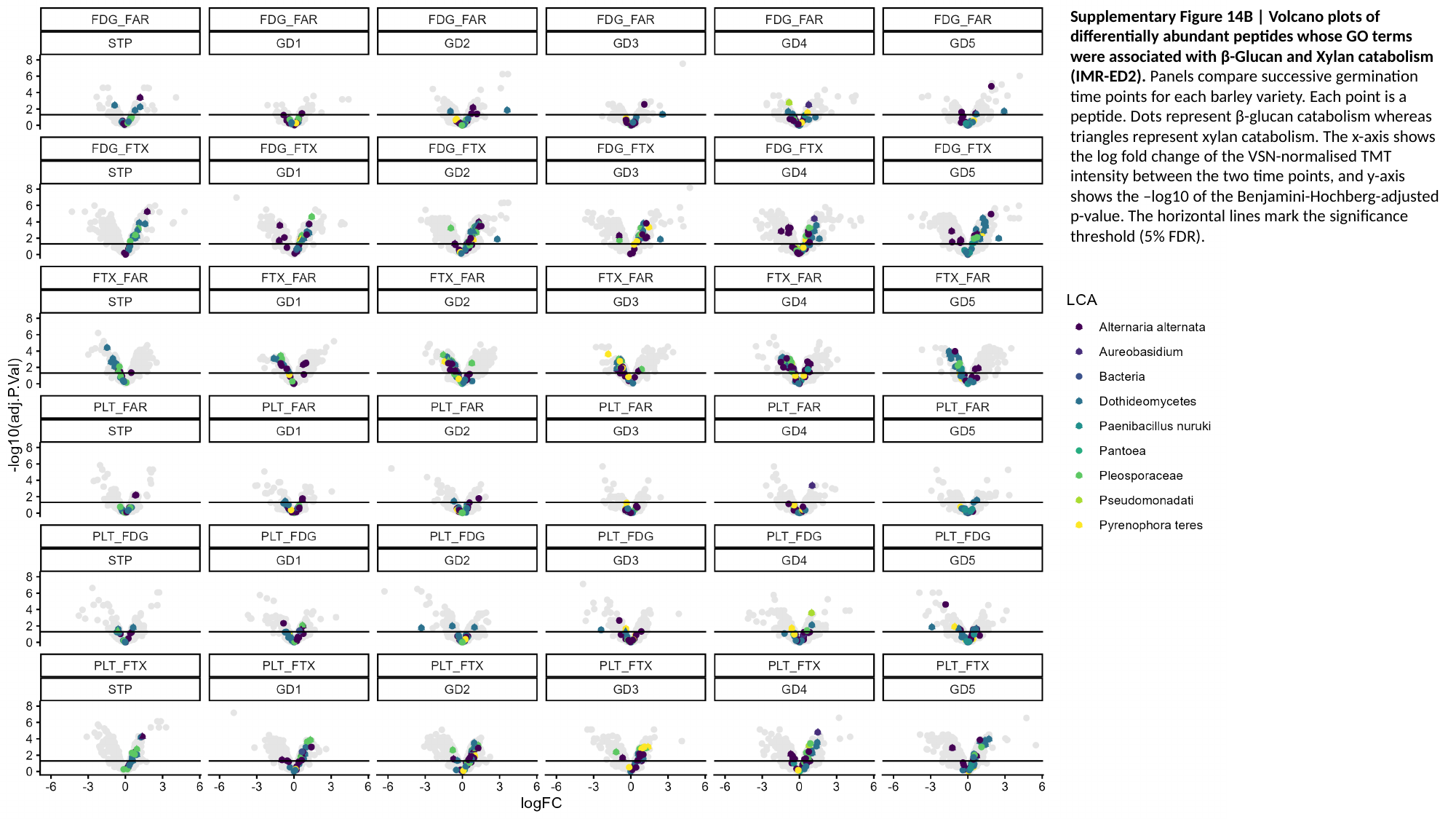

Supplementary Figure 14B | Volcano plots of differentially abundant peptides whose GO terms were associated with β-Glucan and Xylan catabolism (IMR-ED2). Panels compare successive germination time points for each barley variety. Each point is a peptide. Dots represent β-glucan catabolism whereas triangles represent xylan catabolism. The x-axis shows the log fold change of the VSN-normalised TMT intensity between the two time points, and y-axis shows the –log10 of the Benjamini-Hochberg-adjusted p-value. The horizontal lines mark the significance threshold (5% FDR).

### Slide 26
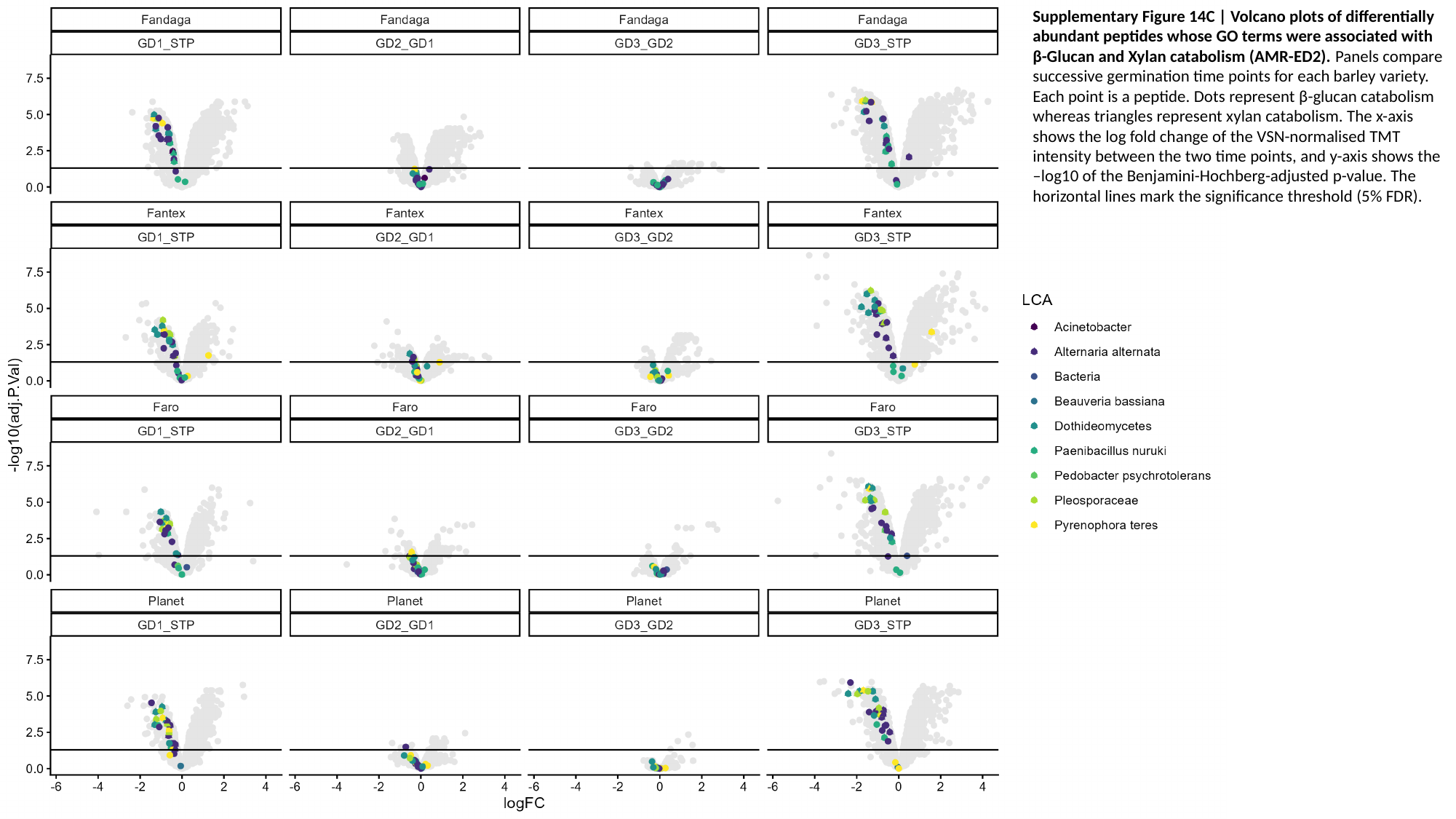

Supplementary Figure 14C | Volcano plots of differentially abundant peptides whose GO terms were associated with β-Glucan and Xylan catabolism (AMR-ED2). Panels compare successive germination time points for each barley variety. Each point is a peptide. Dots represent β-glucan catabolism whereas triangles represent xylan catabolism. The x-axis shows the log fold change of the VSN-normalised TMT intensity between the two time points, and y-axis shows the –log10 of the Benjamini-Hochberg-adjusted p-value. The horizontal lines mark the significance threshold (5% FDR).

### Slide 27
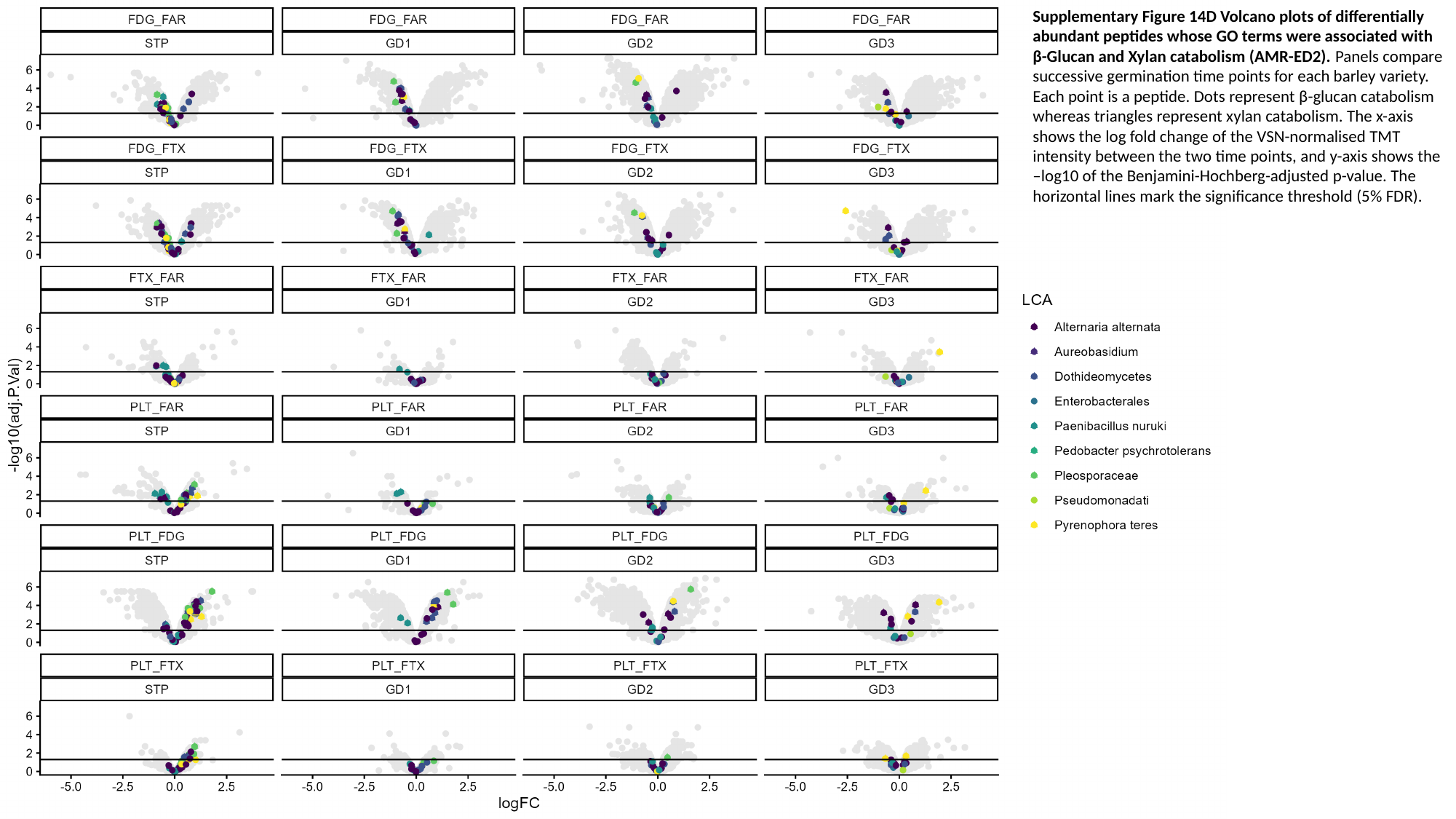

Supplementary Figure 14D Volcano plots of differentially abundant peptides whose GO terms were associated with β-Glucan and Xylan catabolism (AMR-ED2). Panels compare successive germination time points for each barley variety. Each point is a peptide. Dots represent β-glucan catabolism whereas triangles represent xylan catabolism. The x-axis shows the log fold change of the VSN-normalised TMT intensity between the two time points, and y-axis shows the –log10 of the Benjamini-Hochberg-adjusted p-value. The horizontal lines mark the significance threshold (5% FDR).

### Slide 28
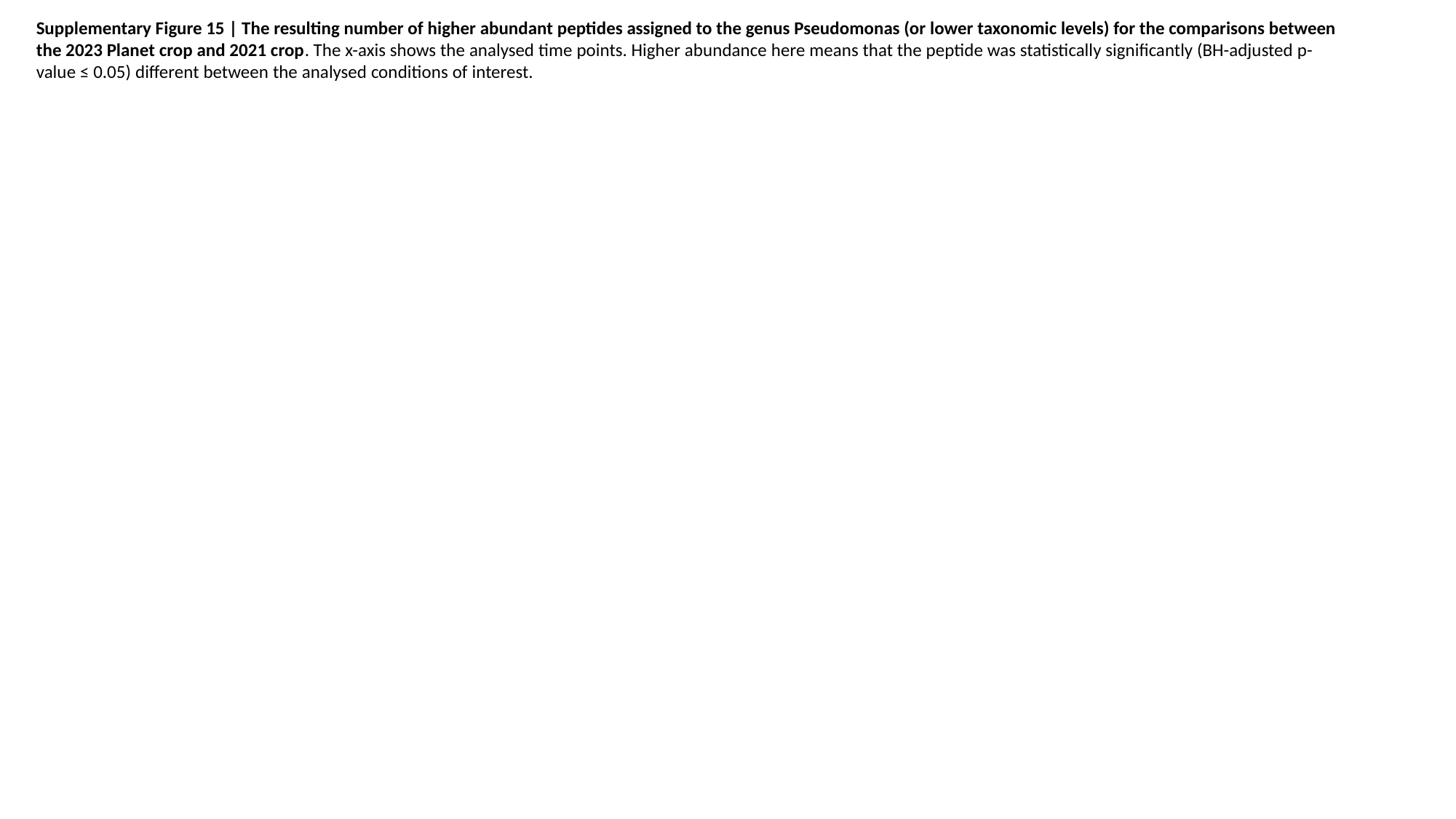

Supplementary Figure 15 | The resulting number of higher abundant peptides assigned to the genus Pseudomonas (or lower taxonomic levels) for the comparisons between the 2023 Planet crop and 2021 crop. The x-axis shows the analysed time points. Higher abundance here means that the peptide was statistically significantly (BH-adjusted p-value ≤ 0.05) different between the analysed conditions of interest.
